# A single-cell multi-omic atlas spanning the adult rhesus macaque brain

**DOI:** 10.1101/2022.09.30.510346

**Authors:** Kenneth L. Chiou, Xingfan Huang, Martin O. Bohlen, Sébastien Tremblay, Diana R. O’Day, Cailyn H. Spurrell, Aishwarya A. Gogate, Trisha M. Zintel, Cayo Biobank Research Unit, Madeline G. Andrews, Melween I. Martínez, Lea M. Starita, Michael J. Montague, Michael L. Platt, Jay Shendure, Noah Snyder-Mackler

## Abstract

Cataloging the diverse cellular architecture of the primate brain is crucial for understanding cognition, behavior and disease in humans. Here, we generated a brain-wide single-cell multimodal molecular atlas of the rhesus macaque brain. Altogether, we profiled 2.58M transcriptomes and 1.59M epigenomes from single nuclei sampled from 30 regions across the adult brain. Cell composition differed extensively across the brain, revealing cellular signatures of region-specific functions. We also identified 1.19M candidate regulatory elements, many novel, allowing us to explore the landscape of *cis*-regulatory grammar and neurological disease risk in a cell-type-specific manner. Together, this multi-omic atlas provides an open resource for investigating the evolution of the human brain and identifying novel targets for disease interventions.

---

The cellular and molecular origins of complex human thought and behavior remain largely a mystery. Historically, proposed explanations have centered on the large relative size [1, 2, 3], high cell numbers [4], or the large cortical surface area and thickness [5] of the human brain. These explanations in isolation, however, fail to explain the many uniquely human faculties, nor do they explain the extreme variety and complexity of impairments that accompany human neurodevelopmental, neuropsychiatric, and neurodegenerative disorders [6]. The human brain is composed of myriad cell types and this cellular heterogeneity contributes to our cognitive and behavioral complexity [7, 8]. Supporting this hypothesis is the observation that the number of distinct cell types in the brain is positively correlated with behavioral complexity across vertebrates [9]. In recent decades, it has been proposed that certain aspects of higher human cognition are supported by specific cell types such as von Economo neurons [10] and “mirror neurons” [11], which have been hypothesized to support intuition and empathy, respectively. These propositions, however, remain largely untested due to gaps in our understanding of the cellular landscape of the human brain and, crucially, differences in cell-type composition and regional heterogeneity among the brains of humans, nonhuman primates, and other animals.

In recent years, the application of rapidly developing single-cell technologies to the brain has begun to address these gaps. Single-cell molecular surveys of targeted regions of the mouse and human brain, for example, have revealed specialized species-specific cell types—e.g., rosehip neurons in humans [12]—and regional biases in cell-type distribution and function [13]. Such atlases are yielding unprecedented cross-species insights into the cellular architecture supporting the structure and function of the brain [14, 15], but the general paucity of comparative nonhuman primate brain atlases has left a conspicuous gap [16]. Moreover, much effort has focused on single molecular modalities (e.g., transcriptomics), typically in only one or a few regions, leaving a lacuna in our understanding of the molecular mechanisms underlying cell function across much of the primate brain.

Here, we generated a 4.2 million cell (combined) transcriptomic and epigenomic atlas across the brain of the rhesus macaque (*Macaca mulatta*), the most widely used nonhuman primate model organism for studies of human perception, cognition, aging, and neurological disease [17]. These single-cell profiles derive from 30 distinct brain regions that collectively represent major cortical, subcortical, and cerebellar areas involved in sensory, cognitive, emotional, and motor functions. Many of these regions are also implicated in one or more clinically relevant neurological disorders. By integrating measures of gene expression and chromatin accessibility, we discover molecular signatures that define cell types across the macaque brain, characterize their distribution and molecular function across disparate anatomical regions, and nominate sets of *cis*-regulatory regions that likely contribute to mature cell fate and function across the brain.

## Results

### A molecular taxonomy of cell types across the primate brain

We generated single-nucleus RNA sequencing (snRNA-seq) data from 30 distinct regions across the cortex, subcortex, cerebellum, and brainstem (*N*=5 animals, 3 female) using sci-RNA-seq3 combinatorial indexing [18, 19] (**Fig. 1A**, **table S1**). With the original sci-RNA-seq3 protocol [20], we generated 1,008,204 single-nucleus transcriptomes from 110 age-, sex-, and hemisphere-matched samples representing 28 brain regions of 10 year-old (mid-adult aged) macaques (N=3 animals; 2 female). Over the course of the study, we implemented improvements in nuclei isolation and preservation [21] which increased nuclear transcriptome recovery by 60% (median unique molecular indices [UMIs], before=202, after=320) and, consequently, the number of nuclei passing our UMI threshold. With the improved protocol, we generated an additional 1,702,081 singlenucleus transcriptomes from the right hemisphere of two animals, the vast majority (N=1,579,908) of which were sampled from 27 brain regions of a single 10 year-old female macaque. Altogether, after applying quality control filters (**Methods**, **fig. S1-S2**), we recovered transcriptome profiles for 2,583,967 nuclei (median UMI per cell=265, median genes expressed per cell=221, **table S2**).

**Fig. 1.**
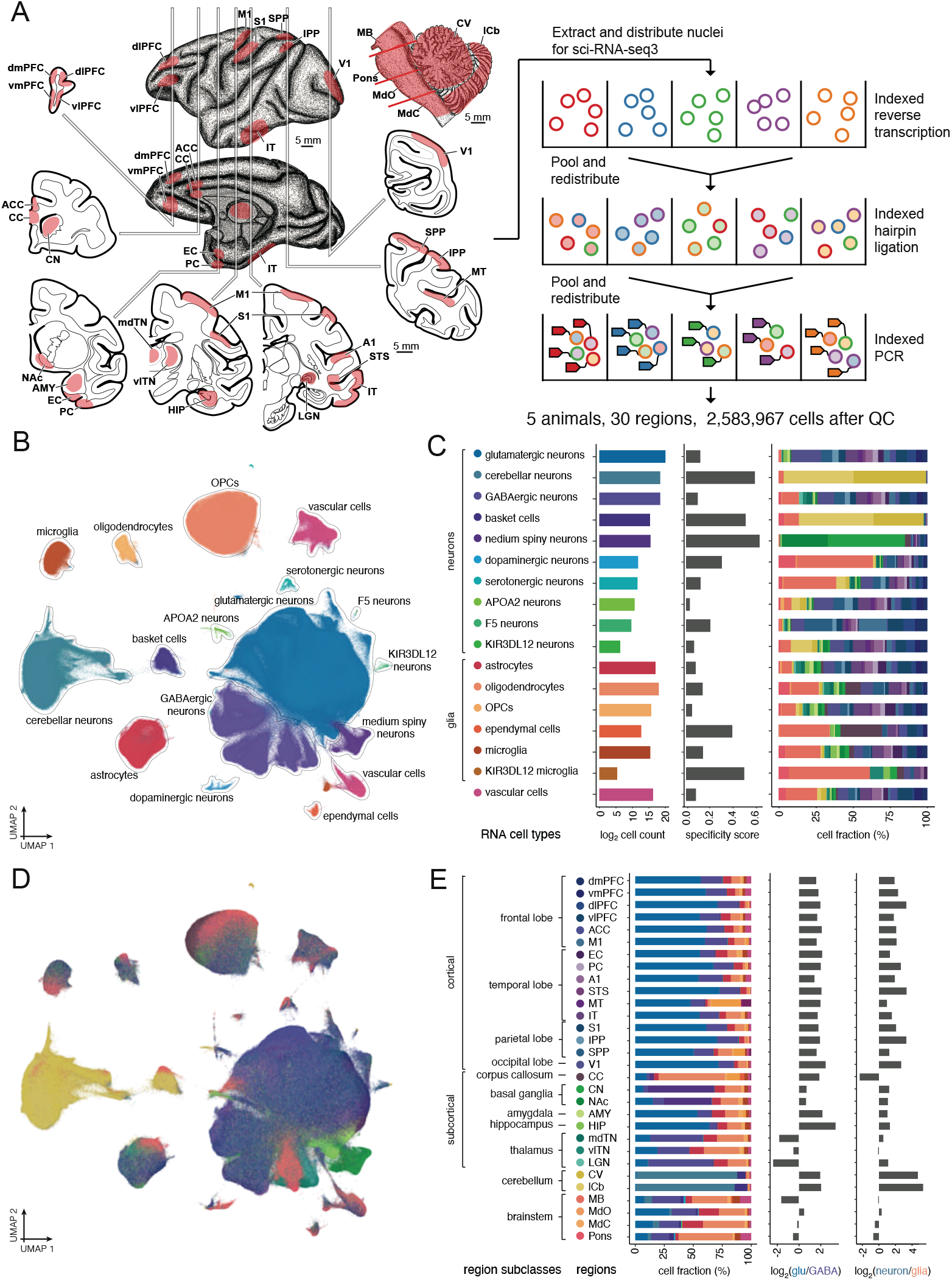
Experimental setup and summary of the Macaque Brain Atlas snRNA-seq dataset. **A**, Schematic of biopsied brain regions for sci-RNA-seq3 experiment. A full list of sampled regions is provided in **table S1**. **B**, UMAP visualization of all snRNA-seq profiled cells colored by cell type (with color code shown in panel C). **C**, Barplots showing the log_2_ transformed cell counts (left), regional specificity score (middle) and regional composition (right, with color code shown in panel E) of each cell type. **D**, UMAP visualization of all snRNA-seq cells colored by cell type (with color code shown in panel E). **E**, Barplots showing the cell type composition (left, with color code shown in panel C), log_2_ transformed ratio of glutamatergic neurons and GABAergic neurons (middle) and log_2_ transformed ratio of neurons and glial cells (right) of each region. Regions are organized by the regional subclass to which they belong.

Controlling for batch effects across sequencing runs (**Methods**, **fig. S3**), we jointly clustered single cell profiles across all sampled brain regions to identify 17 molecularly distinct cell types, which we refer to as ‘cell classes’ (**Fig. 1B-C**). Based on established cell markers (**fig. S4**, **table S3**), we annotated these 17 cell classes as either: (*i*) neuronal cells, including cortical glutamatergic neurons (*CAMK2A*), cortical GABAergic neurons (*GAD1, GAD2*), basket cells (*GRID2, SORCS3*), other cerebellar neurons (primarily granule cells; *GRM4*), medium spiny neurons (*DACH1, PPP1R1B, BCL11B*), serotonergic neurons (*TPH2*), dopaminergic neurons (TH, *DBH);* or (*ii*) non-neuronal cells, including microglia (*DOCK2*), oligodendrocyte precursor cells (OPCs; *VCAN*), astrocytes (*ALDH1A1, GFAP*), oligodendrocytes (MOG, *MBP*), vascular cells (*CFH*), and ependymal cells (*FOXJ1*). Our broad survey also captured four rare, possibly novel cell populations that to our knowledge, have not yet been identified in other studies: three *RBFOX3* + (NeuN+) neuron-like populations (marker genes: *APOA2, N=7,055* cells; *F5, N=880; KIR2DL1/2, N=84)* and one *RBFOX3–* microglialike population (marker gene: *KIR3DL1/2+*, N=44 cells, also *P2RY12+/PTPRC+/ENTPD1*+). Given their rarity, we removed these four cell populations from downstream analyses. Hierarchical clustering of cell classes by the top 50 principal components of gene expression largely recapitulated broad ontogenetic relationships, with most neuronal classes clustering together (dopaminergic neurons being the exception) and the two mesoderm-derived classes (microglia and vascular cells) clustering together (**fig. S5A**).

By sampling across a broad range of anatomical regions within the same individuals, we were able to characterize cellular composition across 30 distinct brain regions—to our knowledge, the most regionally expansive primate single-cell brain atlas to date (**Fig. 1D-E**). The distribution of major cell classes were balanced between sexes and hemispheres (**fig. S6**), but differed extensively across regions, reflecting the cellular makeup underlying region-specific functions (**Fig. 1E**). Unsupervised hierarchical clustering of brain regions according to cell-class composition for the most part conformed to broader anatomical categorizations, with regions of the cortex, subcortex, brainstem, and cerebellum usually grouping together (**fig. S5B**), which was also the case when clustering regions based on the top 50 principal components of gene expression (**fig. S5B**). Two of these four broad regional classes were comprised primarily of a single cell class: in the cortex (N=16 regions, **table S4**), glutamatergic neurons were the most abundant cell type (mean=63.7% of all cells per sample) and outnumbered GABAergic neurons by almost fourfold (**Fig. 1E**; mean=17.4%), while the cerebellum (N=2 regions) was composed almost entirely of cerebellar neurons (mean=85.1%). In contrast, the subcortex (N=8 regions) and brainstem (N=4 regions), were more heterogeneous with respect to their cellular composition, with samples from these regions containing roughly equal proportions of glutamatergic neurons (mean_subcortex_ = 25.1%; mean_brainstem_ = 25.5%), GABAergic neurons (mean_subcortex_ = 20.2%; mean_brainstem_ = 23.0%), and oligodendrocytes (mean_subcortex_ = 18.5%; mean_brainstem_ = 25.5%). We further subdivided the cortical and subcortical samples into ‘region subclasses’ based on neuroanatomical groups (**table S1**), in which there was more limited variation in cellular composition (**Fig. 1E**). For instance, in the subcortex, medium spiny neurons (MSN) comprised around half of the cells in the basal ganglia (nucleus accumbens [NAc] mean=44.7%; caudate nucleus [CN] mean=60.0% MSNs), while the thalamus was enriched for GABAergic neurons (lateral geniculate nucleus [LGN] mean=55.7%; mediodorsal thalamic nucleus [mdTN] mean=43.8%; ventrolateral thalamic nucleus [vlTN] mean=28.6%).

Our broad survey also captured two rarer, but important, cell classes: dopaminergic and serotonergic neurons. These two neurons collectively represented less than 0.3% of all profiled cells (dopaminergic=0.14%; serotonergic=0.12% of all cells) and 0.5% of all neurons (dopaminergic=0.19%; serotonergic=0.17% of all neurons), suggesting that targeted approaches that enrich for these cells (e.g., [22, 23]) are necessary to identify transcriptional variation among subtypes. Dopaminergic neurons, which are found primarily in the substantia nigra pars compacta at low frequency (1.1% of cells sampled in the midbrain vs. mean 0.1% in other sampled regions), are involved in a range of important processes, including voluntary movement, reinforcement learning, and addiction, and their loss is a neuropathological hallmark of Parkinson’s disease [24]. We found that serotonergic neurons were most abundant in the brainstem (mean 0.35% in the 4 brainstem regions vs. mean 0.09% in other sampled regions), where they play a major role in sleep, mood, and appetite, and are key targets of pharmacological therapies for major depressive disorder in humans [25].

### Regional variation in cell subtype composition

To characterize heterogeneity within cell classes, we partitioned the dataset and repeated preprocessing and clustering separately for each of the 17 cell classes. Collectively, we identified 112 distinct clusters (**fig. S7**, **table S5**) that captured neuronal and non-neuronal diversity across the primate brain (**Fig. 2A**). We refer to clusters at this level as ‘cell subtypes’. We identified extensive heterogeneity in glutamatergic (39 subtypes) and GABAergic (20 subtypes) neurons primarily found in the cortex and some regions of the subcortex (e.g., hippocampus, thalamus), while neurons derived from other non-cortical brain regions (e.g., cerebellum, striatum) were transcriptionally distinct and relatively homogeneous within those regions (**Fig. 2A**). This is due in part to the large number of specialized neurons present in some of these regions, including granule and Purkinje cells in the cerebellum, and medium spiny neurons in the basal ganglia (**table S5**).

**Fig. 2.**
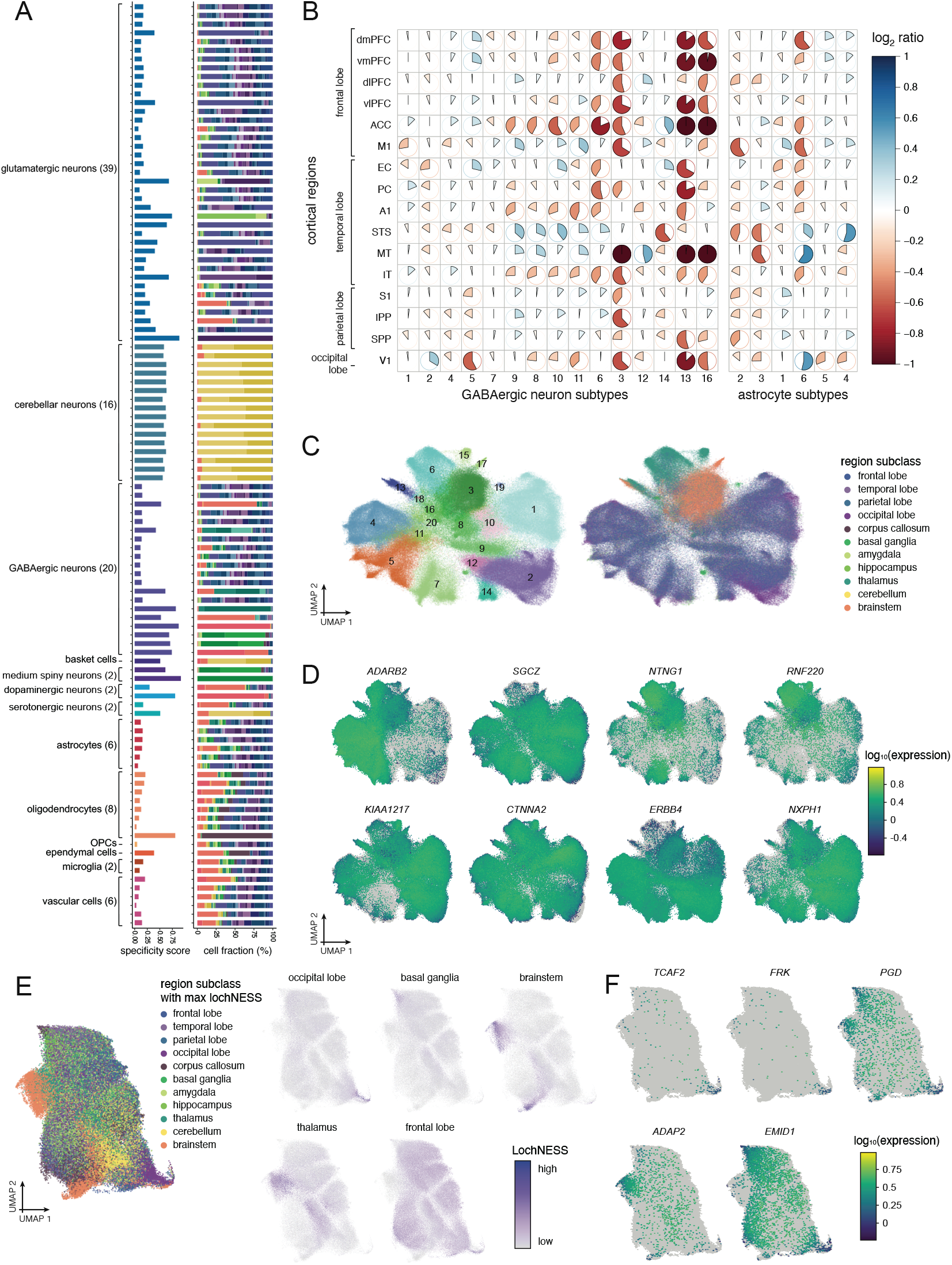
Cell subtype distribution and variation across the brain. **A**, Barplots showing the region specificity score (i.e., Jensen-Shannon divergence statistic) and composition for cell subtypes (with color code shown in **Fig. 1E**). **B**, Heatmap showing scalled log_2_ ratios of GABAergic neuron and astrocyte subtype compositions within cortical region, compared to the average across all regions. Cell subtypes with at least 100 cells profiled are shown in the order of abundance (x-axis, left to right) in the cortical regions organized by region subclasses (y-axis). The color and direction of each pie corresponds to relative enrichment (blue, clockwise) and depletion (red, anti-clockwise) of a cell subtype in a region. Log_2_ ratios were capped at positive and negative 2 prior to scaling. **C**, UMAP visualizations of GABAergic neurons colored by cell subtype (left) and regional subclass (right). **D**, UMAP visualizations of GABAergic neurons colored by cell subtype marker gene expression. **E**, UMAP visualization of astrocytes colored by the region with the highest lochNESS, indicating enrichment of a region subclass in the cell’s transcriptional vicinity. LochNESS distribution in a few example regions (occipital lobe, basal ganglia, brainstem, thalamus and frontal lobe) are highlighted in separate panels as examples. **F**, UMAP visualizations of astrocytes colored by lochNESS-derived region-related marker genes.

Our systematic approach also allowed us to characterize and compare the regional cellular distribution of non-neuronal subtypes, including those of glia, which have not often been the focus on most single cell atlases to date (**Fig. 2A**, **table S6**). Overall, we identified 6 astrocyte, 2 microglial, 7 oligodendrocyte, and 6 vascular cell subtypes, the latter including endothelial cells, smooth muscle cells, pericytes, and both perivascular and meningeal fibroblasts (**fig. S7**, **table S5**) [26]. We compared cell subtypes to published datasets using a NNLS approach [19], and found broad correspondence with subtypes observed in human cortical [14], human brain vascular [26], and macaque hippocampal atlases [27] (**fig. S8A-E**).

To identify cell subtypes that were specific or biased towards a single region or set of regions, we calculated a measure of “regional specificity” using the Jensen-Shannon divergence statistic (**Methods**, [28, 29]). Overall, glial subtypes were more evenly distributed across all regions compared to neuronal subtypes (**Fig. 2A**). This is reflected in lower Jensen-Shannon specificity scores for glial subtypes (mean=0.20; median=0.15; range=[0.04,0.81]) compared to cortical neurons (mean=0.31; median=0.18; range=[0.08,0.89]). A number of cell subtypes, both neuronal and nonneuronal, were highly region-specific. For instance, oligodendrocyte subtype 8, the rarest oligodendrocyte subtype (N=3,439 cells; 1.5% of oligodendrocytes) overwhelmingly derived from the highly myelinated corpus callosum (93.0% of these cells; **Fig. 2A**, **table S6**). Among cortical neurons, GABAergic interneuron subtypes generally exhibited a lower median regional specificities than to glutamatergic neuron subtypes, although there were a number of interneuron subtypes specific to the thalamus (cluster 6) or brainstem (clusters 3 and 16), discussed below.

Given that the regional specificity of excitatory neuronal subtypes has been explored in depth in other studies [30, 31], we focus here instead on populations that are vital for neuronal signal transduction but for which cellular diversity has not previously been explored across the macaque brain. Specifically, we concentrated on the regional diversity of interneurons, because they are important components of long-range circuitry and have been characterized in a few regions across mice, monkeys, and humans [12, 15, 14, 32], allowing us to both benchmark our atlas but also extend current knowledge to understudied regions. We also examine regional distribution among astrocytes, which are crucial for maintaining neuronal homeostasis [33] and are implicated in neurological disorders [34], but have been relatively understudied at the single-cell level.

We pursued three main approaches to dissect the regional heterogeneity within interneuron and astrocyte subtypes, discussed in further detail below: 1) quantification of cell subtype composition to identify nuanced differences in detailed regions within the cortex; 2) identification of regionally specific gene expression programs by analyzing region specific subtypes of interneurons; and 3) in the case of minimal region specific subtypes, leveraging a recently developed statistic to identify regionspecific gene expression patterns in astrocyte subtypes in a cell subtype-agnostic fashion.

Within specific regions of the cortex, cell subtype composition differences become more subtle and require focused quantification. As a first approach, for every sufficiently abundant interneuron and astrocyte subtype in the cortex (>100 cells), we calculated the log_2_ transformed ratio of cell subtype composition in a region, compared to the average composition of that subtype across all cortical regions (**Fig. 2B**). Within the five most abundant interneuron subtypes, we note general balance across all cortical regions, but also observe a relative enrichment of cluster 2 (*PVALB+)* in the occipital lobe (primary visual cortex [V1]) with depletion in regions within the temporal lobe, and depletion of cluster 5 (*ADARB/PAX6+)* in V1. In the superior temporal sulcus (STS) and middle temporal visual area (MT), there is a strong depletion of astrocyte subtype 3 (*LUZP2/GPC5*+) but an enrichment of subtype 6 (*KCNIP4/RBFOX1*+).

Interneurons are the primary drivers of inhibitory control through the release of GABA (γ-aminobutyric acid) and thus strongly impact neural circuitry. Inappropriate development of GABAergic interneurons and subsequent loss of inhibitory regulation contributes to disorders of neurodevelopment, including epilepsy and autism [35, 36]. Despite their importance, the molecular identities and distribution of interneuron subtypes across the adult primate brain remain relatively unknown outside of a few regions [15, 32, 37]. Our snRNA-seq sample captured 371,548 GABAergic interneurons corresponding to 20 subtypes. As a second approach, we focused on gene markers of the region specific interneuron subtypes. Eleven interneuronal subtypes were primarily found in the cortex and could be assigned to four primary interneuron groups that are conserved between mouse and human brains [32], marked by *SST, PVALB, VIP*, and *LAMP5* expression (**fig. S9**). Compared to the cortex, the brainstem and thalamus had a unique distribution of interneuron subtypes (**Fig. 2C**). Thalamic interneurons, which use feed-forward inhibition to relay and tune visual responses to thalamocortical neurons, expressed high levels of *NTNG1* and *RNF220* (**Fig. 2D**), which is indicative of long-range interneurons in the first-order relay nuclei of the thalamus [38]. Sampling across the striatum, which is a critical part of the reward pathway and the largest part of the basal ganglia, a recent single-cell study identified a molecularly unique primate interneuron [32], which was most similar to our GABAergic cluster 18 and represented 15% of interneurons in the CN (**Fig. 2A**).

Astrocytes, the second most abundant non-neuronal cell type in our dataset, are multifaceted support cells of the brain that perform a variety of tasks related to neuronal homeostasis. These tasks can vary across brain regions [33] and astrocyte dysfunction has been linked to neurological diseases, including Alzheimer’s disease [39]. Given these regional differences, we examined whether astrocyte subtypes exhibited regional biases in macaque, similar to what has been observed in the mouse brain [40]. However, while astrocyte subtypes were widely distributed across multiple regions, the cell clusters did not correspond neatly to regions of origin, making claims about inter-region differences in cell composition difficult to systematically analyze across the many regions profiled. To address this complexity, as a third approach, we adapted our recently developed statistic, lochNESS [41], to quantitatively measure regional enrichment in each cell’s “neighborhood” of transcriptionally similar cells. Briefly, for each cell, we tally the number of cells from each brain region in its neighborhood and calculate a focal regional enrichment score (**fig. S10A**, **Methods**). We illustrate the utility of this approach by calculating the lochNESS score on astrocytes at the level of brain region subclasses. Each cell had 11 lochNESS scores calculated, one for each region subclass, with each such score quantifying the enrichment of the given region subclass in a cell’s transcriptional vicinity. We then identified the most enriched region subclass in a cell’s neighborhood and examined the regional heterogeneity agnostic to the cluster-assigned subtype labels (**Fig. 2E**). We also extended the lochNESS to identify genes whose expression can be predicted by lochNESS scores for given regions. To do so, we modeled the lochNESS score of each region in each cell as a function of gene expression with generalized linear regression (**Methods**, **table S7**). The resulting set of genes with significant positive associations with a region’s lochNESS score have higher expression in, and are putatively markers for, cell subtypes in that region.

Using this approach, we identified markers for astrocytes in specific regions (e.g., *TCAF2* and *FRK* in the occipital lobe) and in combinations of regions (e.g., *PGD* in the brainstem, basal ganglia, and thalamus), that we would not have identified if we focused solely on discrete, computationally-defined clusters (**Fig. 2F**). This strategy thus facilitates the identification of more complex regionspecific gene expression patterns. For example, *EMID1*, which is a marker for a subpopulation of astrocyte-like NG2 cells [42], is more highly expressed in astrocytes in the cortex but not in the thalamus, brainstem, or cerebellum. In contrast, *ADAP2*, which is involved in protection from RNA virus infections [43], is highly specific to a subset of astrocytes found in the thalamus (**Fig. 2F**). LochNESS can thus provide a more nuanced approach to identifying regionally-biased cell subtypes and gene expression than conventional clustering. While we focused on astrocytes in this example, lochNESS could be iteratively applied to regions within a subclass in each cell class, e.g. for all glutamatergic neurons across all cortical regions or oligodendrocytes across all subcortical regions (**fig. S10B-D**).

### Joint analysis of single-nucleus transcriptomic and epigenomic data

To complement our transcriptomic dataset and identify key regulatory genomic regions in brain cells, we applied sci-ATAC-seq3 [44, 45] to profile single-nucleus ATAC sequencing (snATAC-seq) epigenomes from nearly all of the brain regions represented in our snRNA-seq dataset. To maximize comparability among datasets, we used 110 of the same age-, sex-, and hemisphere-matched tissue samples (representing the same three animals) profiled in our snRNA-seq dataset. To ensure that the snRNA-seq and snATAC-seq datasets captured the same heterogeneous populations of cells, we homogenized tissue samples on dry ice prior to separately preparing separate nuclei isolations for each library type (**Methods**). Together, the snATAC-seq samples represented 28 of the 30 regions (N=3 animals; midbrain [MB] and MT snATAC-seq data were not generated). After quality control (**Methods**, **fig. S11**), the total number of nuclei profiled was 1,587,880 and ranged from 5,100 (in the closed medulla [MdC]) to 114,410 (in the inferior temporal cortex [IT]) nuclei per region (median=63,739 nuclei per region). We called peaks on a per-sample basis and combined them across all samples based on genomic overlap, resulting in (after filtering) a combined set of 1,192,873 candidate *cis*-regulatory elements (cCREs) spanning 24.4% (725 Mb) of the genome.

We first applied UMAP dimensionality reduction and Leiden clustering to the batch-corrected epigenomic data (**Fig. 3A**) and identified 42 clusters which, based on promoter accessibility, could be assigned to most major cell classes found across the brain (**Fig. 3B**). However, given that unsupervised approaches to cell-type identification are consistently more sensitive using single cell/nucleus RNA-seq data [46], we drew from our transcriptionally defined cell annotations in order to assign cell labels to our snATAC-seq nuclei. To integrate the datasets, we used the graph-linked unified embedding (GLUE) approach [47] and generated a unified transcriptomic and epigenomic embedding of 4,171,847 nuclei (**Fig. 3C-D**). Subsequent cell-type predictions based on our multimodal integration assigned the majority of snATAC-seq nuclei to a cell class (73.7% with confidence ≥ 0.95; **fig. S12**), and captured all of the major cell classes (**Fig. 3D**) with the exception of serotonergic and dopaminergic neurons, which are relatively rare and fairly specific to the MB (which as noted above was not sampled in our snATAC-seq data). The regional distribution of cell classes captured from snATAC-seq and snRNA-seq data were highly concordant, both within regions (**Fig. 3E**) and overall (**Fig. 3F**), which demonstrates that our homogenization and nuclei isolation protocols captured the same heterogeneous populations of cells in the same regions across both modalities.

**Fig. 3.**
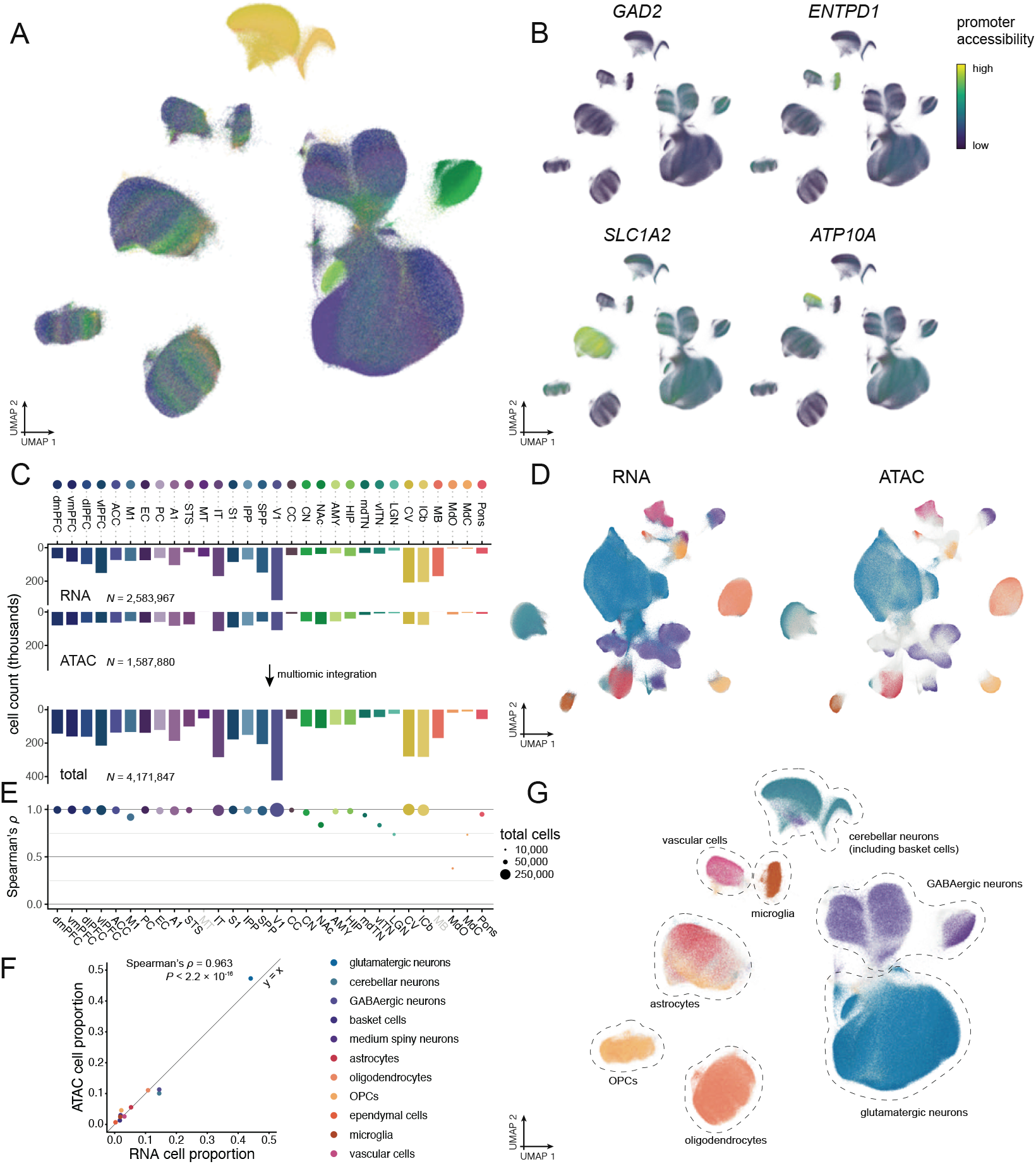
Generation of the Macaque Brain Atlas sci-ATAC-seq dataset and identification of cell classes. **A**, UMAP visualization of all snATAC-seq cells colored by brain region (with color code shown in C). **B**, UMAP visualizations of promoter accessibility scores of cell markers (*GAD2*: GABAergic neurons, *ENTPD1:* microglia, *SLC1A2*: astrocytes, *ATP10A:* vascular cells) reveal high specificity. **C**, Barplots showing nuclei counts by brain region of the snRNA-seq, snATAC-seq, and integrated datasets. **D**, UMAP visualizations of integrated multimodal data, with cell classes colored separately for (left) snRNA-seq and (right) snATAC-seq nuclei (with color code shown in panel F). **E**, Spearman’s rank correlation coefficients showing the correlation between cell-class proportions in the snRNA-seq and snATAC-seq datasets within each region (representing data generated from the same homogenized sample). **F**, Scatterplot showing the correlation between cell-class proportions in the overall snRNA-seq and snATAC-seq datasets (combined across brain regions). **G**, Integration-derived cell-class annotations visualized over the same snATAC-seq UMAP visualization shown in panel A (with color code shown in panel F).

### The gene regulatory landscape of the rhesus macaque brain

We leveraged the scRNA-based cell class annotations (**Fig. 3G**) to explore heterogeneity in cell type-specific gene regulation across the brain. To do so, we partitioned all unique snATAC-seq reads by predicted cell class (**Fig. 3G**), then called peaks separately for each partition using a similar peak-calling approach to that used for the overall dataset, thereby generating an inventory of putative cCREs derived from each cell class in isolation (**Methods**). Across 11 cell classes with snATAC-seq-assigned nuclei, we identified an average of 210,572 peaks per cell class, ranging from 99,323 in microglia to 425,738 in cortical GABAergic neurons (**Fig. 4A**). On average for any given cell class, these peaks covered 7.7% of the genome and 28.8% were found >2 Kb from the nearest gene or promoter (**Fig. 4A**).

**Fig. 4.**
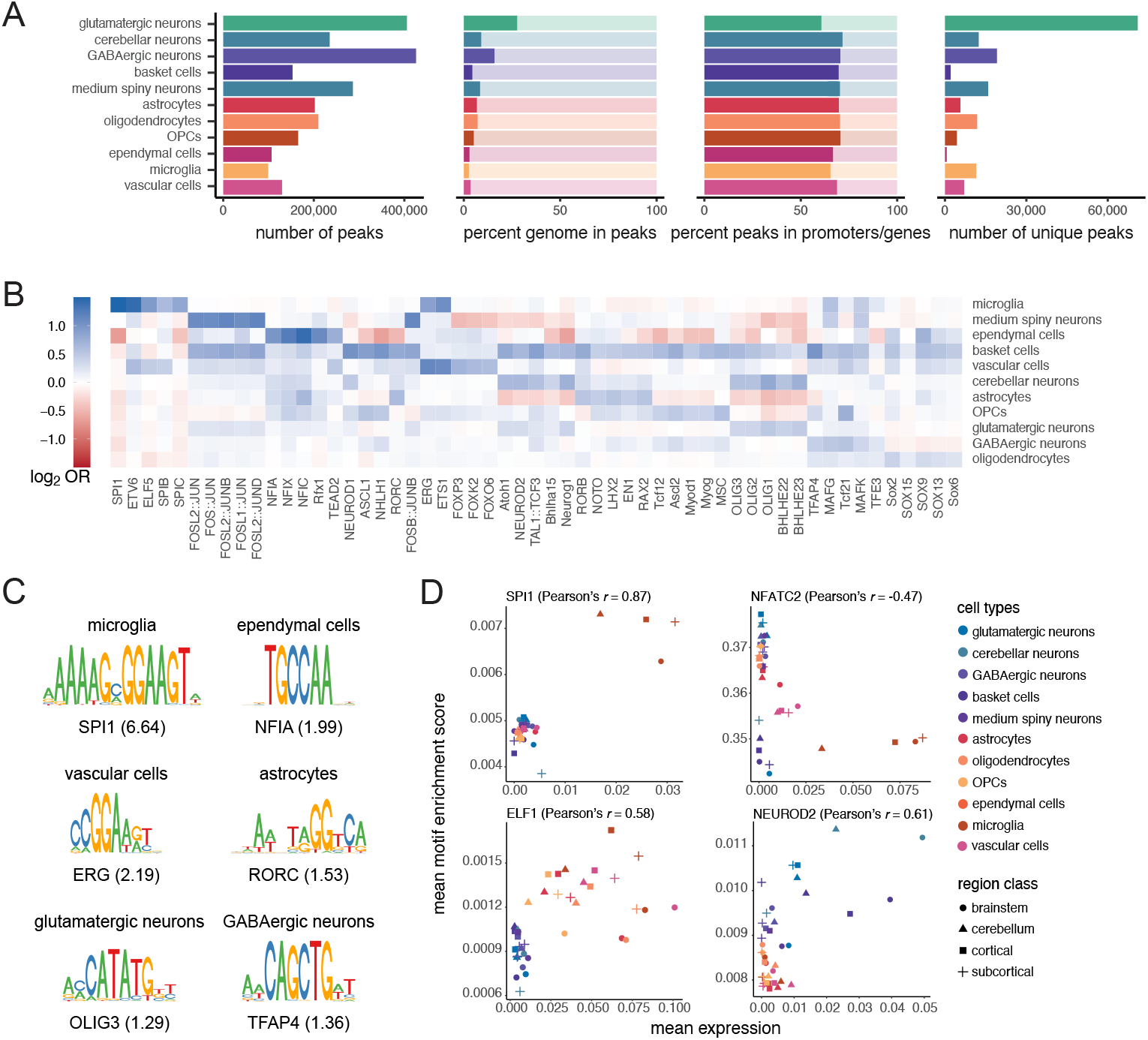
Enrichment of transcription factor binding site motifs in candidate regulatory elements. **A**, Barplots showing summary statistics for peak sets called separately on reads derived from cells assigned to each of 11 cell classes. **B**, Heatmap showing enrichment (log_2_ odds ratio) [OR] of transcription factor (TF) binding motifs among cell classes. The top five most-enriched nonredundant TF motifs (all *P_adj_* < 0.05) are shown per cell class, ordered from left to right by increasing *P_adj_*. Log_2_ OR color ranges are capped at ±1.5. **C**, Position weight matrices of the most-enriched TF motifs for six example cell classes. Odds ratios are shown in parentheses. **D**, Scatterplots showing correlation between snATAC-seq accessibility of TF binding motifs and snRNA-seq gene expression of corresponding TF genes within cell classes in regional classes for four example TFs.

#### Transcription factor regulatory networks

Multi-modal integration of cell specific snATAC-seq and snRNA-seq data allowed us to examine the *cis*- and trans-regulatory links between chromatin accessibilty and gene expression within individual cell types. We first examined putative trans-regulatory factors within cell classes and subtypes. Transcription factors (TFs) are key trans-regulatory proteins that control cell differentiation and function during neurodevelopment [48, 49, 50, 51] and have been implicated in myriad neurodegenerative diseases [52, 53, 54]. The extremely high cell-type specificity of some nuclear TFs have also made them useful targets for identifying and enriching rarer cell types prior to single-cell sequencing [55, 56, 57].

To identify candidate *trans*-acting regulatory networks in each cell class, we carried out TF binding motif enrichment analysis on each set of cell-class-specific peaks, defined as the subset of a cell class’s cCREs that did not overlap with any peaks called in other cell classes (**Methods**, **Fig. 4A**). Cell-class-specific cCREs were highly enriched for many TF binding motifs that are likely involved in cell-specific gene regulation (**Fig. 4B**, **table S8**), including many motifs previously implicated (**Fig. 4C**). For instance, microglial cCREs contained 6.6-fold more binding sites of the nuclear TF SPI1 (also known as PU.1) than expected by chance (P_adj_=1.22×10^-284^; **Fig. 4B-C**). In addition to such canonical examples, we identified numerous motifs that distinguish relatively similar cell classes. For instance, the TF binding motif for NFE2, from the NRF TF family, was most enriched (odds ratio [OR] > 2) in cCREs in both medium spiny neurons (OR=3.07, P_adj_=1.77×10^-87^) and basket cells (OR=2.34, P_adj_=8.76×10^-7^), while the binding motif for NEUROD1 was most enriched (OR>2) in cCREs of basket cells (OR=2.04, P_adj_=3.53×10^-29^), where this TF is necessary for basket cell terminal differentiation and, consequently, axon growth and inhibitory circuit formation [58].

We also characterized TF binding motif enrichment at the cell subtype level. To do this, we extended our multimodal integration and label-transferring approach to each cell class independently by tabulating the reads per-cell falling within cell-class-specific cCREs described above for all cells of a given cell class (**Fig. 3D**). We then integrated the data with corresponding snRNA-seq data of the same cell class using GLUE (**Methods**). The resulting integrated embeddings for each cell class were then used as the basis for predicting cell subtypes, which we carried out on all snATAC-seq cells within each class (**fig. S13**).

Since cell subtypes are preselected to already share broadly similar chromatin accessibility profiles, identifying peaks that are specific to a single subtype—similar to our approach at the cell-class level—was not feasible and left most cell subtypes with no or very few unique peaks to analyze. As an alternative strategy, we carried out differential accessibility analyses among cell subtypes to identify peaks that were predictive of each individual cell subtype within a given cell class (**Methods**). We then identified TF binding motifs enriched in highly differentially accessible regions within cell subtypes (**table S9**). For example, we observed numerous TF binding motifs (N=433, *P_adj_* < 0.05) that were enriched within highly accessible peaks in Purkinje cells, a GABAergic neuron type of the cerebellum that is implicated in autism spectrum disorders (ASD). In our snRNA-seq dataset, of all tested diseases [59], genes associated with autism (DOID:12849) were overrepresented (Fisher’s exact test, OR=10.2, *P_adj_*=8.52×10^-16^) among the top 100 Purkinje-cell marker genes, including *RORA* (fold-change [FC]=331.9), *AUTS2* (FC=43.1), and *SHANK2* (FC=13.8) (**table S10**). Correspondingly, we found that TF motifs enriched in differentially accessible peaks included RORA, four members of the EGR family (EGR1–EGR4), and CTCF (EGR1, EGR3, and CTCF were among the top 5 TF motifs ranked by OR; RORA ranked 182^nd^). RORA is a regulator of circadian rhythm that exhibits decreased expression in ASD brains and may play a role in ASD pathogenesis [60, 61]. EGR-family TFs have been implicated in the disruption of human-specific developmental programs in autism [62]. CTCF is an insulator protein that regulates chromatin structure and may play a critical role in maintaining dendrite structure in Purkinje cells [63] and is also a risk gene for ASD [64].

Given that families of TFs have similar binding motifs, it is often difficult to identify the specific TF in a given family that is responsible for enrichment in cell typespecific cCREs. To identify the most likely TF, we therefore employed our recently-developed approach [45, 65] that uses the computationally paired snRNA-seq and snATAC-seq data. In brief, this approach relies on the assumption that TFs will be highly expressed in cell types where they play a key role, while their associated motif should be enriched (or depleted) in that cell’s cCREs, indicating TF activation (or repression). Overall, we compared the accessibility of 369 TF binding motifs and their corresponding gene’s expression across the cell classes in four region subclasses, with 189 TFs showing positive Pearson’s correlation between gene expression and accessibility of the cognate motif, and 180 showing negative correlation (**fig. S14A**, **table S11**). Among the TFs with largest positive or negative Pearson’s correlation values were strong cell-class-specific activators and repressors (**Fig. 4D**, **fig. S14B**). For instance, *SPI1*, which has been identified as a candidate gene for Alzheimer’s disease via various functional genetics approaches [66], shows a strong activating effect with high expression of the *SPI1* gene and high accessibility for the SPI1 binding motif in microglia. In contrast, NFATC2 has a repressing effect in microglia and vascular cells, as shown by high expression of the *NFATC2* gene associated with lower NFATC2 motif binding in those cell types. We also found evidence for a clear distinction between neurons and non-neuronal cells at two TFs, with ELF1 functioning as a non-neuronal specific activator and NEUROD2 as a neuron-specific activator. Additionally, we note that FLI1, an activator in vascular and microglia cell types, and ELF1 have motif sequences similar to SPI1 (**fig. S14C**), but their activating effects impact a broader set of cell types.

#### The *cis*-regulatory landscape of brain cell variation

We next sought to characterize *cis*-regulatory interactions between cCREs and proximate genes in the rhesus macaque brain. We used two complementary analyses to scan for interactions using our integrated multi-modal dataset. First, we used the regulatory inference framework of GLUE [47], which leverages the unified feature embedding (i.e., joint integration of snRNA-seq genes and snATAC-seq peaks in a common data space) generated during GLUE integration to assess similarity between peaks and genes. Putative regulatory interactions are defined as a high cosine similarity between peak and gene feature embeddings in the unified data space, with statistical significance assessed by permutation [47]. Second, we used a metacell-based approach to aggregate snRNA-seq transcriptomes and snATAC-seq epigenomes into multimodal metacells based on k-means clustering of the unified cell embeddings, then used logistic regression to model the relationship between gene expression and chromatin accessibility within a given metacell [67]. In contrast to the GLUE regulatory score, the logistic regression analysis enabled us to differentiate between positive and negative regulatory interactions between peaks and genes. We considered peak-gene pairs to be putatively regulatory if *P_adj_* < 0.05 for both analyses (**Fig. 5A**, **fig. S15**). For each cell class, we also scanned for differentially accessible peaks using both a regularized logistic regression and a *t*-test, testing accessibility in a given cell class against accessibility in all other cell classes. We consider cCREs with differentially high accessibility (regularized LR coefficient > 0, log_2_ fold-change > 0, and *t*-test *P_adj_* < 0.05, **fig. S16**) to be candidate regulators of cell-type-specific genes (**Fig. 5A**).

**Fig. 5.**
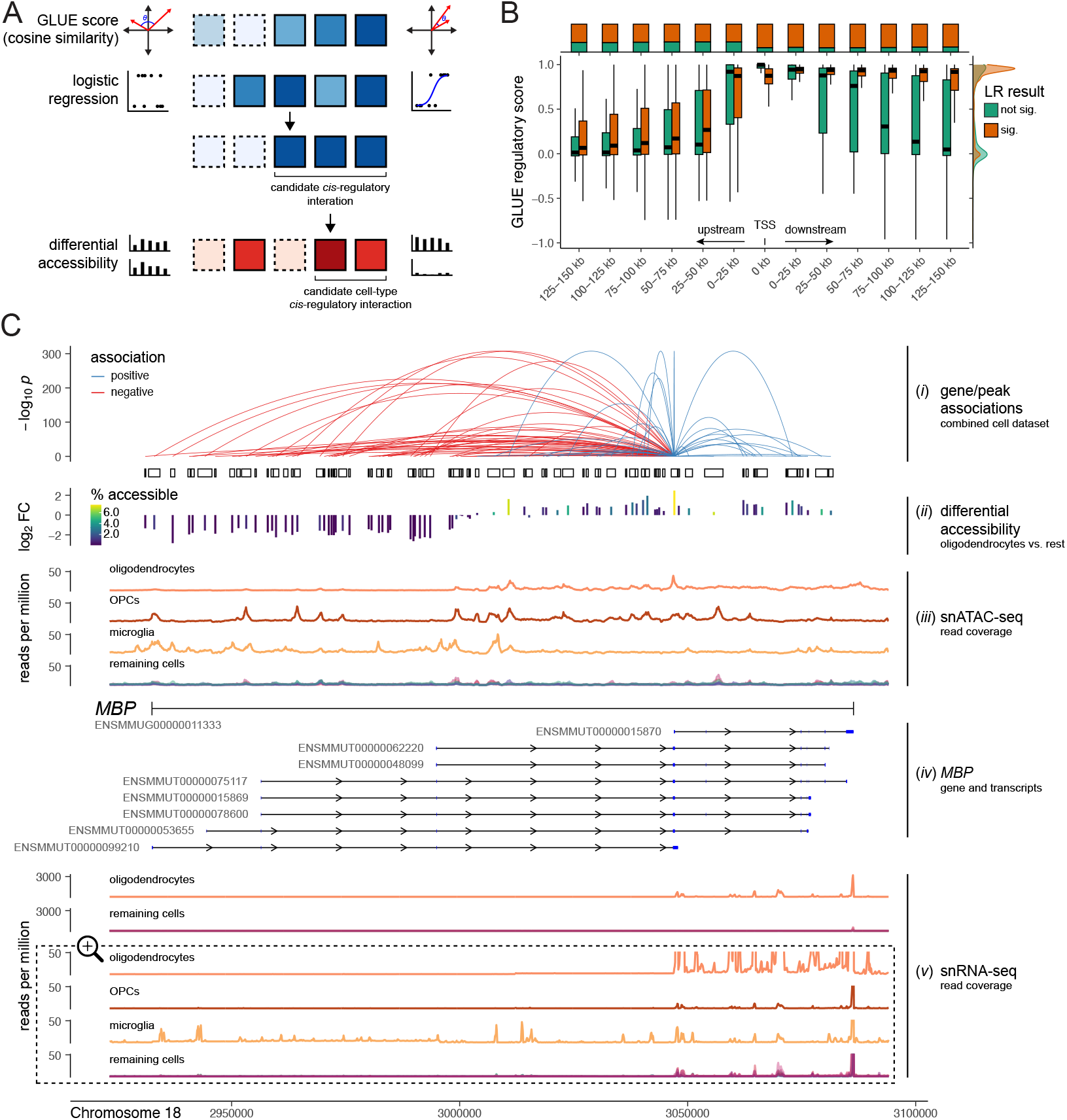
The landscape of *cis*-regulatory interactions in the Macaque Brain Atlas. **A**, Schematic outlining criteria for identification of candidate *cis*-regulatory elements (cCREs). Squares represent peak:gene pairs, darker colors symbolize stronger evidence for a given measure, and solid borders represent statistically significant measures. **B**, Distribution of gene-peak GLUE regulatory scores binned according to the minimum signed distance (left: upstream, right: downstream) between peaks and gene transcription start sites (TSS). Distributions are shown separately according to whether the gene:peak pair also exhibited a significant association (*P_adj_*<0.05) based on the metacell-based logistic regression analysis. The ratio between significant and not-significant gene-peak pairs for proximity bins according to the logistic regression model is shown in the upper margin, while the distribution of GLUE regulatory scores is shown in the right margin. **C**, Candidate regulatory elements are shown in relation to, from top to bottom, (*i*) the strength of and direction of inferred regulatory links connecting peaks to *MBP* expression (based on metacell logistic-regression analysis). The height of links represents the strength (–log_1_0 *P* values) of evidence for regulatory connections and the color symbolizes the direction of the relationship; (*ii*) the differential accessibility (–log_2_ fold change) of peaks in oligodendrocytes relative to all other cell classes; (*iii*) the distribution of normalized snATAC-seq reads by cell class; (*iv*) gene and transcript boundaries of *MBP* and its known isoforms in the rhesus macaque genome, with exons shown in blue; (v) the distribution of normalized snRNA-seq reads by cell class. Oligodendrocyte reads are shown in relation to all other cell classes on the upper portion of the plot. On the bottom portion, the y axis is magnified 60x and cropped to highlight more subtle differences among cell classes.

We focused our analysis on the 6,000 most variable genes in our snRNA-seq dataset and tested all snATAC-seq peaks that fell within 150 Kb of the gene promoter (defined as TSS extended 2 Kb upstream). In total, we tested 223,752 peak-gene pairs (151,083 unique peaks, 5,765 unique genes), of which 142,324 peak-gene pairs (63.6%) met our criteria for being considered candidate *cis*-regulatory interactions (**table S12**). 128,741 peaks (85.2%) that we evaluated were cCREs for at least one gene and 4,811 genes (83.5%) that we evaluated had at least one cCRE.

Of all peak-gene pairs, 132,805 (93.3%) involved a peak that was highly differentially accessible in at least one cell class, thereby fulfilling our criteria for being considered candidate *cis* molecular interactions regulating cell-type-specific markers. cCREs were highly differentially accessible in a maximum of 7 cell classes, with 37% exclusive to a single cell class and 88% highly differentially accessible in 1–3 cell classes.

The vast majority (133,496, or 93.8%) of candidate regulatory interactions were positively associated (i.e., had a positive effect size in the metacell logistic regression)—this held true whether peaks were upstream (13,650/14,575, or 93.7%), downstream (116,939/124,592, or 93.9%), or overlapping (2,907/3,157, or 92.1%) the gene’s transcription start site (TSS). For peak-gene pairs where the peak was upstream of the TSS, the GLUE regulatory scores were highest (indicating high similarity between peak and gene feature embeddings) when peaks were in closer proximity to the TSS (**Fig. 5B**). For peaks downstream of the TSS, GLUE regulatory scores remained high across all distances, with only a modest decrease farther from the TSS (**Fig. 5B**). This result was particularly striking for peaks that had significant, mainly positive, associations between accessibility and gene expression, likely reflecting (*i*) higher global accessibility across the gene body resulting from higher expression of the gene (as opposed to distal regulation) and/or, (*ii*) methodological limitations of using a single gene-wide TSS (i.e., the most upstream TSS of all isoforms), thereby ignoring variation in TSS positioning among isoforms, which likely vary in their usage across tissues and contexts [68].

Using the cell-class-specific gene expression and cCRE peak sets, we repeated our integration, regulatory inference, and differential accessibility workflows on each cell class individually. We tested a mean of 72,914 peak-gene pairs (range: 45,539–114,200) per cell class and identified a mean of 11,442 peak-gene pairs (range: 881–41,966) showing evidence of regulatory interactions (**fig. S17** and **table S13**).

To illustrate how these maps of putative interactions might be useful to investigate the regulatory landscape at the level of an individual locus, we focused on the myelin basic protein (*MBP)* gene (**Fig. 5C**), which encodes one of the most abundant proteins in central nervous system myelin [69, 70], has a range of splice isoforms [71], and is a canonical marker of oligodendrocytes. *MBP* is located on chromosome 18 (positions 2,932,531–3,086,873) on the rhesus macaque (Mmul_ 10) genome and has 8 annotated mRNA isoforms (Ensembl). In humans, classic MBP isoform 3 (18.5 kDa) predominates in adult myelin [71].

In our global peak set (all cells), 94 peaks fell within 150 Kb of the *MBP* promoter and were included in our analysis. Of these peaks, 83 (88.3%) were identified as candidate regulators of *MBP* (crMBP), with 38 crMBPs (45.8%) positively associated with *MBP* expression. Of all crMBPs, only one was not located within the MBP gene boundaries—it was, however, located less than 2 Kb upstream within the likely promoter region.

In accordance with the well-known status of *MBP* as an oligodendrocyte marker, we found that *MBP* was differentially expressed in oligodendrocytes, with detected expression in 80.9% of cells and 1,434-fold higher expression than all other cells averaged together. Finegrained inspection of normalized read distributions from oligodendrocyte nuclei revealed the highest densities of snRNA-seq reads corresponding to the polyadenylation site (position 3,086,373) and snATAC-seq reads corresponding to the TSS (position 3,046,976) of a single transcript, ENSMMUT00000015870, indicating that it is likely the dominant *MBP* isoform expressed in adult macaque oligodendrocytes.

By examining the genomic-distance relationships between crMBPs and the dominant *MBP* transcript in adult oligodendrocytes, we found that all 16 crMBPs that either overlapped or were downstream of the isoform’s TSS were positively associated with *MBP* expression. Among the 67 crMBPs that were located upstream of the TSS, 22 (32.8%) were positively associated with *MBP* expression while 45 (67.2%) were negatively associated. Several of these negatively associated crMBPs corresponded with sci-ATAC-seq3 peaks in other cell types, particularly oligodendrocyte precursor cells (OPCs) and microglia (**Fig. 5C**). However, the accessibility landscape of OPCs is overall more similar to that of oligodendrocytes across the region upstream of the TSS of the dominant isoform, with greater accessibility at most peaks except for that of the promoter of the dominant isoform (**Fig. 5C**). As OPCs play a critical role in myelinogenesis by giving rise to oligodendrocytes [72], these crMBPs likely serve as critical markers of the OPC-oligodendrocyte transition, during which the expression of this gene, and this isoform in particular, is massively upregulated.

#### Enrichment of disease heritability among candidate regulatory elements

Lastly, we used our cCREs to identify cell-type-associated regulatory networks that may drive polygenic disease risk. We tested for enrichment of disease trait heritability using the linkage disequilibrium score regression (LDSC) tool [73, 74], after lifting over macaque cCREs to human genome coordinates [28]. We tested a total of 53 phenotypes relevant to neurological diseases, disorders, syndromes, behaviors, or other traits (**table S14**), and examined enrichment among cell-class cCREs called separately in each of 11 cell classes.

Our results broadly recapitulated several known roles of cell classes in neurological disease (**Fig. 6** and **table S15**). For example, sites associated with cardioembolic stroke (OR=32.2) or ischemic stroke (OR=9.2) were enriched (*P_adj_* < 0.05) only in vascular cells, which play a crucial role in forming and maintaining the blood-brain barrier [75]. We also found that Alzheimer’s disease-associated sites were enriched only in microglia—a result replicated using loci from three independent genome-wide association studies (GWAS) (OR range: 13.9–15.0)—consistent with the prominent role of microglia proliferation and activation in Alzheimer’s disease [76].

**Fig. 6.**
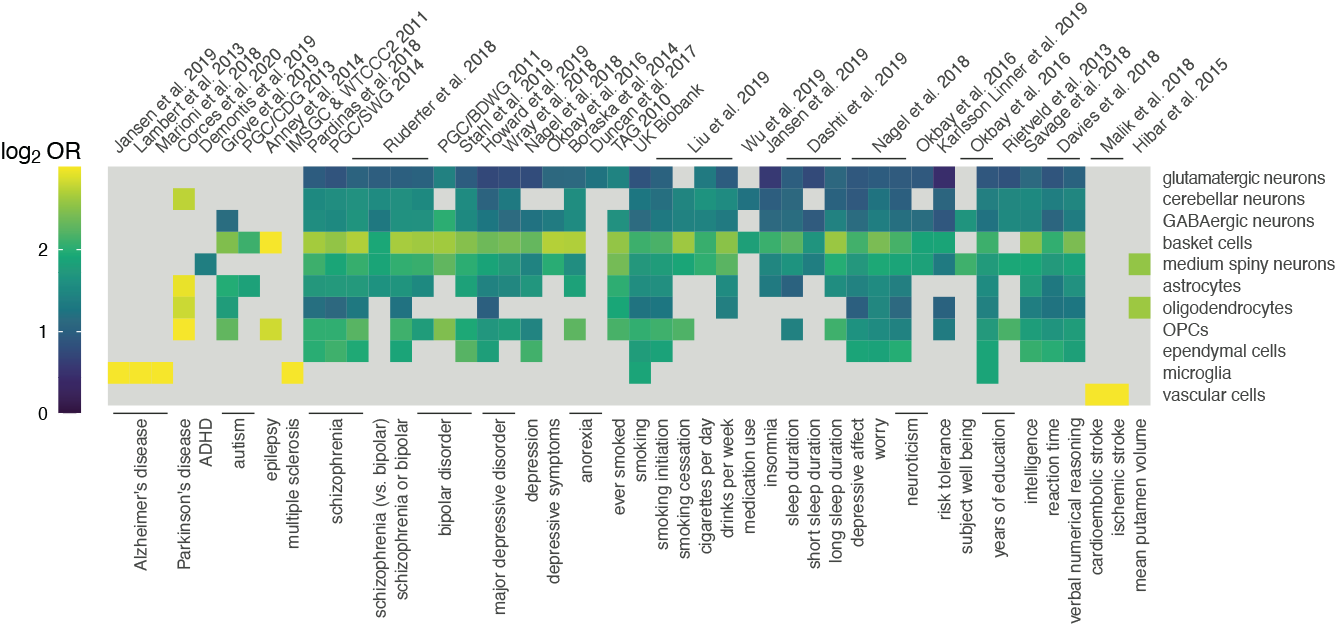
Enrichment of heritable disease-relevant sites among candidate regulatory elements. The heatmap displays heritability enrichment (log_2_ odds ratio [OR]) of diseases among cell-class snATAC-seq peaks for tested diseases, syndromes, and phenotypes. Only results passing a threshold of *P_adj_* < 0.05 are shown. The log_2_ OR color range is capped at 3.0.

Across all cell classes, basket cells were enriched for the greatest number (N=37) of GWAS phenotypes, including disorders such as schizophrenia (OR range: 5.9–6.2), bipolar disorder (OR range: 5.6–6.2), and major depressive disorder (OR range: 5.1–5.3), and, most strongly, epilepsy (OR=9.0)–a disease that basket cells have been connected to in animal models and some genetically-linked human forms of the disease [77].

Other notable results included the enrichment of multiple-sclerosis-associated sites among open regions in microglia (OR=46.6), highlighting the outsized role of these immune cells in the etiology of multiple sclerosis and as a putative therapeutic target [78, 79]. In multiple sclerosis, disease-associated microglia alter their transcriptional profiles and may contribute to neuroinflammatory processes underpinning this autoimmune disorder [79]. We also found enrichment of Parkinson’s disease-associated sites among open regions in the glial OPC, oligodendrocyte, and astrocyte cell classes (OR range: 7.0–8.4). In Parkinson’s disease, glial cells may play a major role in the progressive degeneration of dopaminergic neurons [80], a classic hallmark of Parkinson’s disease, or in alterations to glutamatergic neurotransmission [81].

Finally, we found that heritable sites associated with attention deficit/hyperactivity disorder (ADHD) in our analysis were enriched only among open regions of medium spiny neurons. While the magnitude of the enrichment was relatively mild (OR=2.6, *P_adj_*=0.031), genetic variants associated with ADHD have been historically difficult to identify, with the first risk loci only recently reported [82]. Medium spiny neurons have been linked to behavioral hyperactivity and disrupted attention via activation of astrocyte-mediated synaptogenesis [83]. Our results therefore suggest that medium spiny neurons may be a promising target for future ADHD-related study.

## Discussion

Understanding the cellular architecture of the adult primate brain is crucial both for understanding the evolution of human cognition and behavior as well as for identifying mechanisms underlying neurological disorders. In service of these goals, we used snRNA-seq and snATAC-seq to derive a molecular atlas spanning the adult rhesus macaque brain, comprising data from over 4 million cells profiled from 30 brain regions. Based on our multimodal molecular data, we identified 112 distinct molecular cell types or subtypes and characterized their distribution across the macaque brain, adding to the growing number of primate single-cell molecular brain atlases [15, 32]. The data are freely available (NeMO archive, nemo:dat-rtmm5q2) and will serve as a rich resource for the neuroscience and neurogenomics communities.

In generating a multi-region transcriptomic and epigenomic atlas of the most widely used nonhuman primate in neuroscience, we: (*i*) identified all of the major brain cell classes and many cell types that have been previously reported (**Fig. 1**, **Fig. 2**); (*ii*) quantified regional distribution of cell types and subtypes within individuals, which allowed us to identify compositional differences in samples collected at the same time and from the same animals (**Fig. 2**); (*iii*) identified rare and regionally specific cell types (e.g., Purkinje cells), which may facilitate the development of molecular tools such as cell typespecific viral vectors that, in combination with new technologies such as CellREADR (Cell access through RNA sensing by Endogenous ADAR) [84] may enable precise targeting of cell types based on their unique patterns of chromatin accessibility and gene expression; (*iv*) characterized multiple *trans*- and *cis*-regulatory mechanisms that differentiate cell classes and subtypes (**Fig. 4A-D**, Fig. 5); and (*v*) identified numerous associations between genetic risk for neurological disorders and the epigenomic states of specific cell types (**Fig. 6**).

This single-cell atlas of the adult primate is notably generated from samples collected from healthy adults. The paired nature of the dataset, with regions sampled from the same individual brains, avoids many of the inter-individual variables (e.g., genotype, environment) that can impact neurological development and function. The atlas may thus be a valuable resource for characterizing molecular features that play a role in myriad neurological disorders. The relatively few unique individuals sampled also represents a limitation of the current study—we currently know very little about how brains of healthy individuals differ in cell composition and function and what that confers for disease susceptibility and/or progression. Given continuing improvements in cost and throughput of single-cell sequencing, characterizing multi-region cellular variation across many healthy individuals is becoming not only a possibility, but also an emerging priority for the field.

To our knowledge, these data represent the largest and most comprehensive multimodal molecular atlas in a primate to date and provide a resource for exploring how the heterogeneous molecular and cellular composition of the brain gives rise to the behavioral complexity of primates including humans. We anticipate that these data will also provide a critical and much-needed molecular and neurobiological map of complex human-relevant social behavior and disease, as well as an extensive substrate for comparative analyses across animal brains.

## Acknowledgments

We thank the management and staff of the Caribbean Primate Research Center (CPRC) (particularly Angelina Ruiz-Lambides, Carlos Sariol, and Armando Burgos-Rodríguez) for maintaining the Cayo Santiago and Sabana Seca field stations. We also thank members of the CPRC (particularly Samuel Bauman, Nicole Compo, and Carlos Pacheco), the CBRU biobanking team (Christopher Walker and Jack Stylli), the Snyder-Mackler Lab (Beth Slikas, Sarah Ford, Marta Koperska, and Layla Brassington), the Platt Lab (Lora Assi), and the Tung Lab (Jenny Tung and Tawni Voyles) for assistance with sample collection and/or logistics. We are grateful to Silvia Domcke, Chengxiang Qiu, Gürkan Yardimci, Junyue Cao, Hannah Pliner, Brent Ewing, and Anne Buckley for assistance and/or feedback through various stages of this manuscript, and to Gil Speyer, Ben Readhead, the DNASU Plasmid Repository, and the ASU Research Computing team for generously sharing computational resources and/or support. The data reported here were generated via the single cell platform of the Brotman Baty Institute (BBI) for Precision Medicine.

## Author contributions

NSM, JS, MLP, MJM, LMS, KLC, and XH conceived the study. KLC, MJM, and NSM collected samples, with logistical support from CBRU and MIM. MOB performed neuroanatomical dissections, assisted by KLC, TMZ, MJM, and NSM. KLC, DRO, CHS, and TMZ performed lab work. AAG managed data and provided bioinformatic support. KLC, XH, TMZ, and NSM analyzed the data, with input from MOB, ST, MGA, LMS, MJM, MLP, and JS. KLC, XH, MOB, ST, MJM, MLP, JS, and NSM wrote the paper. All authors edited and approved the manuscript.

## Consortium authors

The members of the Cayo Biobank Research Unit are Susan C. Antón, Lauren J. N. Brent, James P. Higham, Melween I. Martínez, Amanda D. Melin, Michael J. Montague, Michael L. Platt, Jérôme Sallet, and Noah Snyder-Mackler

## Data and materials availability

The data analyzed in this study were produced through the Brain Initiative Cell Census Network (BICCN, RRID:SCR_015820) and deposited in the NeMO Archive (RRID:SCR_002001) under identifier nemo:dat-rtmm5q2 accessible at https://assets.nemoarchive.org/dat-rtmm5q2 All code for this project is available through the following GitHub repositories: https://github.com/bbi-lab/bbi-dmux (sci-RNA-seq3 demultiplexing), https://github.com/bbi-lab/bbi-sciatac-demux (sci-ATAC-seq3 demultiplexing), https://github.com/bbi-lab/bbi-sci (sci-RNA-seq3 preprocessing up to count matrix generation), https://github.com/bbi-lab/bbi-sciatac-analyze (sci-ATAC-seq3 preprocessing up to count matrix generation), and https://github.com/CayoBiobankResearchUnit/macaque-brain-atlas (remainder of the analyses).

## Methods summary

The detailed materials and methods are available in the supplementary materials. Briefly, we collected fresh-frozen brains from 5 adult rhesus macaques that were part of the free-ranging Caribbean Primate Research Center research colony on Cayo Santiago. We focused our atlas on 30 anatomically defined regions that are associated with key cognitive, behavioral, and disease traits. To allow for the profiling of multiple genomic modalities from the same representative cell populations, we pulverized all samples on dry ice to homogenize and divide tissue for single nucleus sequencing. We generated single-nucleus RNA-seq data from 2,583,967 nuclei spanning a total of 30 unique regions from both hemispheres of the brain, and paired those data with single-nucleus ATAC-seq data from 1,587,880 28 regions across 28 unique regions. These data were generated using sci-RNAseq3 [19] and sci-ATACseq3 [45] combinatorial indexing. Single-nucleus libraries were deeply sequenced and processed using a uniform protocol that included extensive QC filters (**fig. S1-S2**).

Using Leiden-clustering on snRNA-seq nuclei [19], we identified 17 primary cell classes and then iteratively clustered each cell class for deeper annotation of cell subtypes. Whenever external data were available, we validated our cell classifications using a non-negative least squares (NNLS) approach [19] to identify correlations between cell subtypes and annotated labels in reference datasets. We then identified marker genes for each cell class and subtype, characterized the regional distribution and expression of each cell class and subtype across the brain, and identified cell-specific enrichment of disease-associated genes.

To connect snATAC-seq profiles to snRNA-seq nuclei, we used the GLUE integration approach [47], which allowed us to annotate all snATAC nuclei based on the cell classes and subtypes identified in our snRNA-seq data. These connections allowed us to carry out a range of analyses, including TF binding site enrichment, linking TF enrichment to and TF expression within cell types, and identifying cell-specific regulatory links between candidate *cis*-regulatory elements (cCREs) and nearby genes. Lastly, following coordinate liftover between the primate and human genomes, we used LDSC [73, 74] to quantify enrichment of neurological disease-associated variants in cell class biased cCREs.

Raw sequencing data and the annotated count matrices are available through NeMO (RRID:SCR_ 002001), protocols for data generation are on protocols.io (DOI:10.17504/protocols.io.9yih7ue and DOI:10.17504/protocols.io.be8mjhu6); and scripts to process samples and recreate all analyses are available on GitHub (**Data and Materials Availability**).

## Online methods

### Study population and sample collection

All animals sampled in this study are rhesus macaques (*Macaca mulatta*) from the semi-free-ranging colony on the island of Cayo Santiago, Puerto Rico. Maintained by the Caribbean Primate Research Center (CPRC) within the University of Puerto Rico, the Cayo Santiago macaque colony has been largely continuously studied since its founding in 1938 [85]. All present-day macaques are descended from an initial founder population of 409 animals and have since maintained an outbred population structure despite generations of isolation [86]. Apart from being provisioned with commercial feed and occasionally subject to capture-and-release sampling, the macaques otherwise live in naturalistic conditions, subject to minimal intervention and manipulation, as approved by IACUC. The study used animals that needed to be removed from Cayo Santiago [87] and were immediately euthanized. Standardized tissue collection and sample archiving was coordinated by the Cayo Biobank Research Unit (CBRU), which provided the brain samples used in this study [88, 89].

Procedures for necropsy, brain removal, and dissection followed those previously described for this population [89] and are briefly outlined here. Following veterinary euthanasia, brains were perfused with sterile saline, removed from the cranium, and hemisected into left and right hemispheres using a long single-edge razor blade. After sectioning off the cerebellum/brainstem from each hemisphere, the cerebral hemispheres were placed on custom molds (designed either for left or right hemispheres) and coronally sectioned into 11 roughly 5-mm-thick blocks, numbered in order rostral to caudal. All 12 blocks (with the cerebellum/brainstem considered block 12) were then sealed in Whirl-pak bags, flash-frozen in liquid nitrogen vapor, and archived in ultralow −80°C freezers. The interval between euthanasia and permanent storage of frozen tissue averaged 51 minutes, with a standard deviation of 5.8.

All procedures were performed in accordance with the NIH Guide for the Care and Use of Laboratory Animals and were approved by the Institutional Animal Care and Use Committee at the University of Puerto Rico (protocol #338300). Five macaques were included in this study (**table S2**). The vast majority of the data derived from four 10-year-old macaques, which are considered middle-aged adults in this population [90, 89].

### Region selection and biopsy

Frozen brain blocks were placed on a dissection tray over dry ice in order to keep tissue frozen during biopsy collection. Individual blocks were then moved from the dry ice to a tray sitting on wet ice, allowing for tissues to be acutely warmed to the point that biopsies could be taken from targeted structures. Biopsies were made using a cutting spoon (Fine Science Tools, Inc., cat. #10360-13). Dissected brain regions are listed in **table S1** and approximate locations for biopsy are illustrated in **Fig. 1A**. For a given structure, attempts were made to minimize inclusion of off-target surrounding tissues (e.g., white matter underlying a targeted gray matter structure). Below, we document the most common block numbers where structures were located. Due to interindividual differences and/or variation in sectioning, regions of interest were sometimes identified and dissected from adjacent blocks based on neuroanatomical landmarks. Alternate block numbers are therefore also documented below.

The most anterior block sampled for this study (block 2) contained gray matter for the dorsomedial (dmPFC), ventromedial (vmPFC), dorsolateral (dlPFC), and ventrolateral prefrontal cortices (vlPFC). dmPFC and vmPFC were defined as being on the medial side of block 2. The dmPFC biopsy was pulled from the gray matter in the top half of the medial edge of the block. A space along the medial edge was left to separate dmPFC from vmPFC. The vmPFC biopsy was pulled from the medial ventral half of the tissue block. Biopsies of dlPFC came from the cortical tissues surrounding the dorsal lateral portion of the block that included the superior and inferior portions of the principal sulcus. Samples from vlPFC came from the ventral and lateral portion of the block. As was the case on the medial side, a portion of the cortex was left between each lateral biopsy to avoid overlap (**Fig. 1A**).

Block 3 (sometimes 4) contained biopsies for the anterior cingulate cortex (ACC), corpus callosum (CC), and head of the caudate nucleus (CN). The biopsy for ACC was the gray matter sitting between the CC, which is ventral to ACC and the cingulate sulcus, which sits dorsal to the cingulate gyrus. CC was defined as the white matter track sitting ventral to the ACC and medial to the lateral ventricle. The CN was the gray matter sitting ventrolateral to the lateral ventricle and surrounded on all other sides by white matter. The CN was the only biopsy in the second block that was scooped out of the block face to minimize inclusion of any white matter sitting anteriorly past the CN within the block (**Fig. 1A**).

Block 5 (sometimes 4 or 6) contained the amygdala (AMY), entorhinal cortex (EC), perirhinal cortex (PC), and nucleus accumbens (NAc). The NAc is located ventral to the caudate, internal capsule, and putamen (Pu). Furthermore, in fresh-frozen tissue, there was a slightly darker color to the NAc. The tissue making up the NAc was scooped out of the block face. Similarly, the AMY was identified as ventral to the Pu, medial to the ventral portion of the claustrum, and dorsal to the EC. The AMY was also scooped out to minimize the inadvertent collection of neurons within the hippocampus (HIP). Finally, the EC and PC were collected, the delineation between the two was the rhinal fissure (**Fig. 1A**).

Blocks 5–6 (sometimes 4 or 7) contained tissue that were biopsied to represent cortical regions primary motor cortex (M1), primary somatosensory cortex (S1), primary auditory cortex (A1), superior temporal cortex (STS), and inferior temporal cortex (IT). Subcortical structures that were biopsied included mediodorsal thalamic nucleus (mdTN), ventrolateral thalamic nucleus (vlTN), lateral geniculate nucleus (LGN), and hippocampus (HIP). The delineation between M1 and S1 was the central sulcus and were taken from the approximate central third of the lateral portion of each respective gyrus. Within a case, attempts were made to biopsy from approximately the same putative mototopic and somatotopic regions. A1 biopsies were taken from the dorsal portion of the superior temporal gyrus which is within the ls (i.e., inferior operculum). The gray matter forming the STS sits ventral to the superior temporal gyrus and dorsal to the inferior temporal gyrus. IT was defined as the gray matter forming the lateral portion of the inferior temporal cortex. mdTN sits bilaterally on midline, within the thalamus. It is bound by ventricles dorsally, laterally by the centrolateral thalamic nucleus and ventrally by the centromedial thalamic nucleus. vlTN is bound by the centrolateral thalamic nucleus medially, body of the caudate nucleus (CN) dorsally, and the reticular thalamic nucleus laterally. Biopsies for mdTN were taken from the central and central medial portions of the nucleus, while vlTN biopsies were taken from the central portion of the nucleus. In both cases, this was in an effort to avoid inclusion of other thalamic nuclei. The LGN is a 6 layered structure that is easily observed on the coronal face of fresh-frozen slabs. When observed, the biopsy was scooped out. Like the LGN, the HIP was defined by its classic cytoarchitectonic features within the medial temporal lobe. For biopsies, efforts were made to not include EC, which sits ventral and ventromedial to HIP (**Fig. 1A**).

Block 7 (sometimes 6 or 8) contained tissues representing the superior posterior parietal (SPP), inferior posterior parietal (IPP), and area MT (MT). SPP biopsies were from the gray matter of the superior lobule. The intraparietal sulcus sits between SPP and IPP. Therefore, IPP biopsies were taken from the gray matter of the second, more lateral lobule. Finally, area MT was defined by the gray matter of the insular cortex, bound on its medial edge by white matter of the extreme external capsule and laterally by the superior and inferior operculum divided by the superior temporal sulcus (**Fig. 1A**) [91, 92].

The final cerebral block, block 11, contained the visual cortex. Biopsies from primary visual cortex (V1) were taken from the dorsolateral surface gray matter above the external calcarine sulcus (**Fig. 1A**).

The hemisected cerebellum/brainstem block was dissected as follows. First, the cerebellum was dissected off and the cerebellar vermis (CV) was separated from the lateral cerebellar cortex (lCb). Next, the remaining brainstem was dissected such that the midbrain (MB) block was separated by making a cut from just behind the inferior colliculus to the top of the basilar pons. Next, the pons was separated from the medulla by making a cut from the stria medullaris (approximate center of the fourth ventricle) to the base of the pons. A final cut at the base of the fourth ventricle to separate the open medulla (MdO) from the closed medulla (MdC) (**Fig. 1A**).

To allow for the profiling of multiple genomic modalities from the same representative cell populations, we pulverized all biopsies on dry ice to homogenize and divide tissue for downstream experiments. We followed the tissue pulverization procedures described by Domcke et al. [93] to achieve a powder consistency on a sterile aluminum foil work surface. Once sufficiently pulverized, we stirred the sample thoroughly, then divided the sample using the folded edge of foil as a funnel into new 1.5 ml pre-chilled and pre-labeled microcentrifuge tubes. Foil and tubes were set on aluminum trays or tube racks set on dry ice to keep powdered tissue frozen throughout this process. We divided samples into roughly a 2:1 ratio given the expected efficiencies/yields for single-nucleus RNA-seq and single-nucleus ATAC-seq protocols, respectively. Pulverized tissue was stored at −80°C up until processing for downstream library preparation procedures.

### snRNA-seq data generation

To profile single-nucleus gene expression, we performed single-nucleus RNA-seq (snRNA-seq) using the three-level single-cell combinatorial indexing RNA-seq (sci-RNA-seq3) approach [19], which is the improved version of the original sci-RNA-seq protocol [18].

For two out of the three experimental batches in our dataset, we used a protocol closely adhering to the sci-RNA-seq3 protocol described by Cao *et al.* [19]. For the third batch, we used the improved protocol (“tiny sci”) described by Martin *et al.* [21]. Sample order was randomized between the first two batches, and within the third batch, to minimize batch effects and other technical artifacts.

For the first two batches, we slightly modified the protocol described by Cao *et al.* [19, 20] for a different tissue type and smaller input amounts. Briefly, we added 50 μl of cell lysis buffer to pulverized tissue in a 1.5 ml microcentrifuge tube, then homogenized the tissue using 5–10 strokes with a disposable RNase-free plastic pestle (Fisherbrand, cat. #12-141-364). We then added another 950 μl of cell lysis buffer, mixed by pipette, then transferred the suspension through a 70 μm cell strainer (pluriSelect cat. #43-10070-70) into a 15 ml conical tube containing 5 ml ice-cold 4% paraformaldehyde. Nuclei were fixed in 4% paraformaldehyde for 15 min with occasional mixing, washed once in 1 ml ice-cold nuclei wash buffer, then suspended in 200 μl nuclei wash buffer. Nuclei were counted by mixing with 1 μM of YOYO-1 iodide (TheroFisher cat. #Y3601) using a Countess II FL automated cell counter (Life Technologies), divided into tubes in 100 μl aliquots, then flash-frozen in liquid nitrogen.

For nuclei fixed with paraformaldehyde, library construction was similar to the sci-RNA-seq3 method from Cao *et al.* [19] with minor modifications including the substitution of Quick Ligase (NEB) for 10 minutes at 25°C for the second index step, instead of T4 DNA ligase (NEB) for 180 minutes at 16°C. For tagmentation, we used N7 adaptor-loaded Tn5 from QB3 MacroLab at the University of California Berkeley in tagmentation buffer (2X TD) as previously described in Corces *et al.* [94]: 20 mM Tris-HCl, pH 7.5, 10 mM MgCl_2_, 20% (vol/vol) dimethylformamide (DMF). Libraries were sequenced on a NextSeq or NovaSeq platform (Illumina) (read 1: 34 cycles, read 2: 100 cycles, index 1:10 cycles, index 2: 10 cycles).

For the DSP/MeOH nuclei isolations and library construction based on Martin *et al.* [21], we used hypotonic lysis buffer solution B (with BSA) for small volume tiny sci-RNA-seq3 nuclei isolation methods. For sci-RNA-seq3 library construction, we loaded 20,000 nuclei per index 1 reverse transcriptase (RT) well in a 384 RT-well experiment with mouse and human brain added as separate quality control nuclei and nuclei from cell lines HEK293T (RRID:CVCL_0063) and NIH/3T3 (RRID:CVCL_0594) combined as barnyard controls per RT plate. Nuclei from all RT plates were pooled and redistributed to ligation plates for the second index as previously published; after the addition of the second index, nuclei were again re-pooled for their final distribution of 4,000 nuclei per well prior to second strand synthesis, protease digestion, tagmentation and PCR all on this final third index plate.

### snRNA-seq pre-processing

snRNA-seq sequencing reads were processed into a gene-by-nucleus expression matrix of unique molecular index (UMI) counts following the methods described by Cao *et al.* [19]. We used largely an identical pipeline which, briefly, (1) converts base calls to fastq files with bcl2fastq/v.2.20 (RRID:SCR_015058) (Illumina), (2) removes adapter sequences using Trim Galore/v.0.6.7 (RRID:SCR_011847) [95], (3) aligns trimmed reads to a reference genome with STAR/v.2.7.6 (RRID:SCR_ 004463) [96], (4) extracts mapped reads, (5) removes duplicates, and (6) generates UMI counts for exonic and intronic regions of each gene, tabulated according to the unique three-level barcode design in sci-RNA-seq3. We used the rhesus macaque reference genome (Mmul_ 10) [97] and annotation, obtained from Ensembl (version 101) (RRID:SCR_002344). We extended the 3’ UTR annotations of genes and transcripts by 500 bp to avoid misclassifying genic reads as intergenic. The remainder of our pipeline followed the procedures described by Cao *et al.* [19]. After generating the count matrix, we removed all nuclei with UMI counts < 100.

For each sample, we imported gene-by-nucleus count matrices into the AnnData/v.0.8.0 (RRID:SCR_ 018209) [98] framework, then ran Scrublet/v.0.2.3 (RRID:SCR_018098) [99] (expected_doublet_rate=0.05) to calculate doublet scores. We marked nuclei as doublets if they had Scrublet doublet scores > 0.20. For each sample, we additionally marked nuclei as doublets using per-sample thresholds determined by Scrublet and adjusted by eye as necessary in order to separate bimodal peaks visualized on the Scrublet doublet score histogram (**fig. S1**).

To further identify potential doublet nuclei, we employed an iterative clustering strategy [100] implemented with Scanpy/v.1.9.1 (RRID:SCR_018139) [101]. First, we combined all nuclei into a single AnnData object and filtered nuclei to those with UMI ≥ 100, number of expressed genes < 2,500, and a percentage of reads mapping to the mitochondrial genome < 5%. We then removed all non-autosomal genes, genes located on unplaced scaffolds, and unexpressed genes. Next, we normalized the data to the total UMI per nucleus, logarithmized the data, and subsetted the data to the 10,000 most variable genes. For each cell, we regressed out total UMI counts per nucleus, then mean-centered and scaled the data. The dimensionality of the data was then reduced by PCA (50 components). To further reduce the dimensionality, we ran a UMAP (using umap-learn/v.0.5.2) (RRID:SCR_018217) analysis [102] with BBKNN/v.1.5.1 (RRID:SCR_022807) [103] to simultaneously correct for batch differences. For the BBKNN integration, we set neighbors_within_batch=10 (given three batches, tantamount to UMAP n_neighbors=30), used the cosine distance metric, and used the PyNNDescent/v.0.5.6 algorithm (RRID:SCR_022806) [104]. We then ran UMAP using the settings min_dist=0, spread=1.0, and n_components=10 to facilitate clustering (https://umap-learn.readthedocs.io/en/latest/clustering.html). For data visualization only (not clustering), we ran a similar BBKNN/UMAP pipeline with neighbors_within_ batch=5 (for three batches, tantamount to UMAP n_ neighbors=15), min_dist=0.25, spread=1.0, and n_ components=2. To cluster the data, we exported and imported the 10-dimensional UMAP matrix into Monocle3/v.1.2.9 (RRID:SCR_018685) [19] in R/v.4.0.2 (RRID:SCR_001905) [105], then implemented the Leiden-clustering workflow in Monocle3 with a relatively high-resolution setting (resolution=1 ×10^-4^). For each cluster, we then calculated the mean Scrublet doublet score and marked all clusters with a mean Scrublet doublet score > 0.15 as doublet clusters (**fig. S1**).

After identifying doublets as described above, we removed all marked doublets and repeated the normalization, dimensionality, and clustering procedures almost exactly as described above, with the only difference being a coarser cluster resolution setting in Monocle3 (resolution=1 ×10^-5^). We confirmed adequate removal of doublet cells by observing the clean separation of distinct cell types and the absence of clusters expressing obviously ambiguous marker gene profiles (**fig. S1**).

#### Removal of sci-RNA-seq cell contamination

During the course of cell-type identification (see following section), we observed the presence of two distinct clusters of cells (**fig. S2A**) with expression profiles resembling embryonic progenitors (markers genes, unknown cluster 1: *ASPM, CENPE, CENPF, MKI67*; unknown cluster 2: *COL1A1, COL1A2, FN1, VIM*), an unusual finding in adult primate brain samples. Because these were present in relatively large proportions in some samples (25%)—but at low levels overall (2.2%)—and because our sci-RNA-seq experiments included control samples of exogenous (i.e., non-macaque brain) origin (specifically, a fetal mouse brain positive control and a “barnyard” sample consisting of mixed human HEK293T and mouse NIH/3T3 cells), we tested for the presence of contaminating nuclei of exogenous origin. We identified and removed contaminating cells as follows.

Because the only non-macaque samples included in all experiments were the control samples of either human or mouse origin, we used BBSplit/v.38.38 (RRID:SCR_016965) [106] to assign reads to the macaque, human, or mouse genomes. BBSplit is a competitive aligner that maps to several references simultaneously, assigning reads to the genome with the best unambiguous match. We used the following reference assemblies from Ensembl version 101: Mmul_ 10 (macaque), GRCh38.p13 (human), and GRCm38.p6 (mouse). After indexing the three references simultaneously using BBSplit, we aligned 10 million randomly sampled unique (de-duplicated) reads for each sample using default settings in BBSplit, which partitioned reads assigned to each genome into separate fastq files. Unmapped and ambiguous reads were directed to additional fastq files that were not used. Using a similar demultiplexing workflow to the sci-RNA-seq3 preprocessing pipeline, we tabulated reads-per-cell for each of the three genomes and calculated summary statistics.

After filtering to only cells with ≥10 unambiguously assigned reads by BBSplit, we observed that discernible fractions of exogenous reads (reads unambiguously assigned to human or mouse) were specific to certain barcodes from the first round of sci-RNA-seq barcoding (reverse transcription), indicating that a low level of cross-well contamination of cells or barcoded primers likely occurred at this stage (**fig. S2C**). We also observed that, after filtering to cells passing all previous quality control filters, our clustering and annotation workflow had partitioned exogenous cells into two clusters corresponding to human and mouse cells respectively, with no discernible exogenous contamination in other annotated cell types (**fig. S2B**). After removing the entirety of the two exogenous clusters from the dataset (N=58,443 cells), we re-examined the distribution of exogenous read fractions across reverse-transcription barcodes and confirmed that human and mouse cells were effectively removed (**fig. S2C**).

### snRNA-seq cell-type and cell-subtype identification

To identify cell types, we visualized the expression of canonical marker genes (**table S3**) on normalized, log-transformed gene expression data using Scanpy. Most clusters were readily assigned to well-characterized cell types in this manner. To aid in the classification of more nuanced cell types, we determined top marker genes using logistic-regression and *t*-test marker-gene methods implemented via the ‘rank_genes_groups‘ function in Scanpy. For each discrete cell type, we ran marker gene tests by testing gene expression in a given cell type against gene expression in all other cells in our dataset.

Based on canonical markers and data-derived marker genes, we identified 17 parent cell types (not including the two cells of exogenous origin, see section above), which we refer to as cell classes. In all but two cases, our parent cell types corresponded with partitions identified through our clustering using Monocle3 (q-value threshold=0.05). In two cases, we considered clusters assigned to the same partition to be discrete parent cell types because they exhibited clear separation in our global analysis while clearly expressing canonical markers of known cell types (dopaminergic and serotonergic neurons; **table S3**), yet did not effectively segregate when their assigned partition (the partition also including GABAergic neurons) was analyzed separately.

To identify cell subtypes, we partitioned the data by cell class and reanalyzed each data partition individually. For each cell-class-specific analysis, we repeated a preprocessing, dimensionality reduction, and clustering analysis that largely followed the pipeline described above for our global analysis, with the following exceptions. After normalizing and log-transforming the data, we identified the 2,000 most variable genes for each given cell type and subset the data to those highly variable genes. Because we observed that differences in total UMI among batches resulted in artifactual clusters being identified downstream (even after batch-correction with BBKNN, a problem we did not observe in our global analysis), we regressed out total UMI counts per nucleus separately for each batch. We then combined residual values from all batches before mean-centering and scaling for PCA and UMAP analysis. For Leiden clustering, we used the same resolution parameter (resolution=1 × 10^-5^) for most cell types, but in four cases defaulted to partitions identified using Monocle3 (q-value threshold=0.05) after observing small clusters with unusually high UMI. We considered clusters/partitions identified in this manner to be cell subtypes.

As with our global (all cell classes combined) analysis, for each cell subtype we identified top marker genes using logistic-regression and *t*-test marker gene methods implemented in Scanpy. Additionally we used a nonnegative least squares (NNLS) approach [19] to identify correlations between cell subtypes and annotated labels in reference datasets (**table S5**, **fig. S8**).

Additionally, we scanned for gene-disease associations that were enriched among the top 100 marker genes for each cell subtype. We used gene-disease associations from the DISEASES database (RRID:SCR_ 015664) [59] and used Fisher’s exact test to identify overrepresented disease associations among the top 100 marker genes for a given cell subtype, using all macaque genes in our analysis as the background (**table S10**).

### Cell composition and regional heterogeneity analysis

To assess the specificity of cell classes and/or subtypes, we calculated the Jensen-Shannon divergence statistic using the ‘JSD‘ function from the philentropy package (RRID:SCR_022805) [107] in R. We calculated the Jensen-Shannon divergence by comparing, for a given cell class or cell subtype, the cell type’s count distribution across brain regions to the count distribution (combining all cell types per region) of the entire dataset combined [29].

To measure regional heterogeneity within cell types, we extended our recently developed statistic, lochNESS [41], to quantitatively measure enrichment of each region subclass or region within each cell’s neighborhood. For each cell type, we define lochNESS of *cell_n_* for *region_m_* as:

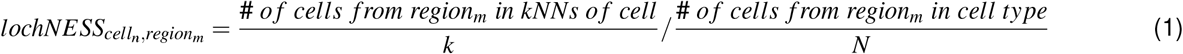

where *N* is the total number of cells in the the cell type and *k* is the number of nearest neighbors for *cell_n_*. For each cell type the calculation results in a cell × region matrix, where each row can be separately visualized. For a summarizing visualization, each cell can be colored by the region with the largest lochNESS. Additionally, when we focus on a subset of regions (e.g. just the cortical regions), we calculate a normalized lochNESS that is comparable across the regions of interest:

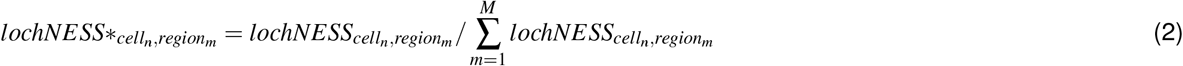

where *M* is the number of regions or region subclasses of interest.

To identify genes that are expressed with regional bias, we fit a regression model for each gene to identify regions with significant non-zero correlation with gene expression as implemented in Monocle3 [19]. The model for each cell type is:

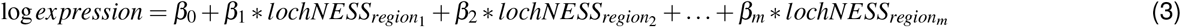

where *β*_0_ is the intercept and *lochNESS*_*region*_1__ is a vector of lochNESS across all cells in the cell type.

### Hierarchical clustering of cells and regions

We used Scanpy to cluster cell classes and brain regions based on the top 50 principal components (PCs) of gene expression. Because of our use of BBKNN for batch correction in our main workflow, our PCA was not actually corrected for batch. To rectify this, we first used the harmonpy/v.0.0.5 (RRID:SCR_022798) implementation of Harmony (RRID:SCR_022206) [108] to generate a batch-corrected PCA matrix (convergence after 2 generations). We then used the Scanpy ‘dendrogram‘ function to perform hierarchical clustering using the batch-corrected PCA embedding. To visualize uncertainty, we performed 1,000 bootstrap iterations in which we resampled cells randomly with replacement and computed new dendrograms. We then used the ‘DensiTree‘ function [109] implemented in the phangorn/v.2.6.3 (RRID:SCR_ 017302) [110] R package to visualize trees. We performed this procedure using both cell class and brain region as labels (**fig. S5A-B**).

For brain regions, we also performed hierarchical clustering using the cell proportion (cell class×brain region) matrix. We used the ‘hclust‘ function in R to cluster using the ‘complete’ method based on Euclidean distances. To again visualize uncertainty, we resampled all cells in our dataset 1,000× with replacement, then repeated calculation of cell class proportions and hierarchical clustering. We visualized the final tree with ‘DensiTree‘ (**fig. S5B**).

### snATAC-seq data generation

To profile single-nucleus chromatin accessibility, we performed single-nucleus ATAC-seq (snATAC-seq) using the three-level single-cell combinatorial indexing ATAC-seq (sci-ATAC-seq3) approach [45], which is the improved version of the original sci-ATAC-seq protocol [44]. We followed the protocol of Domcke *et al.* [93], with slight modifications. Briefly, we added 50 μl of Omni-ATAC lysis buffer to pulverized tissue and homogenized the tissue with 5–10 strokes with a disposable RNase-free plastic pestle (Fisherbrand, cat. #12-141-364). We then added another 950 μl of Omni-ATAC lysis buffer, mixed by pipette, incubated on ice for 3 min, then transferred the suspension to a new 15 ml conical tube containing 5 ml ATAC-RSB with 0.1% Tween-20. We then pelleted the nuclei, removed the supernatant, and resuspended the pellet in 1 ml of 1x DPBS. We then transferred the suspension through a 70 μm cell strainer (pluriSelect cat. #43-10070-70) into a 15 ml conical tube containing 4 ml of 1x DPBS and 140 μl of 37% formaldehyde (final concentration 1% formaldehyde). We then incubated the nuclei for 10 min with occasional mixing. The fixation was then quenched with 250 μl 2.5 M glycine, incubated for 5 min at room temperature, then incubated for another 15 min on ice. We then pelleted the nuclei, removed the supernatant, and resuspended the pellet in 2 ml freezing buffer. Nuclei were counted by mixing with 1 μM of YOYO-1 iodide (TheroFisher cat. #Y3601) using a Countess II FL automated cell counter (Life Technologies), divided into tubes in 50 μl aliquots, then flash-frozen in liquid nitrogen

Frozen fixed nuclei were prepared for the sci-ATAC-seq3 library similar to Domcke *et al.* [45]. Omni-ATAC lysis buffer (10 mM NaCl, 3 mM MgCl_2_, 10 mM Tris-HCl pH 7.4, 0.1% IGEPAL CA-630, 0.1% Tween 20 and 0.01% Digitonin) was used to permeabilize fixed nuclei before diluting samples with ATAC-RSB (10 mM NaCl, 3 mM MgCl_2_, 10 mM Tris-HCl pH 7.4) supplemented with 0.1% Tween-20. Approximately 200,000 nuclei per sample was spread across 4 wells for tagmentation as previously described. Barnyard control for each set of experiments included mouse cell line (CH12-LX; RRID:CVCL_0211) and human pancreas as a quality control tissue.

Our combined snATAC-seq dataset encompasses data prepared using five sci-ATAC-seq3 experimental runs (i.e., library preparation/sequencing batches). Sample order was randomized between batches to ensure balance of brain regions, sex, and hemispheres between runs and to minimize batch effects.

### snATAC-seq pre-processing

snATAC-seq sequencing reads were processed into a peak-by-nucleus count matrix following the methods described by Domcke *et al.* [45]. We followed largely an identical pipeline which, briefly, (1) converts base calls to fastq files with bcl2fastq/v.2.20 (Illumina), (2) removes adapter sequences using Trimmomatic/v.0.39 (RRID:SCR_011848) [111], (3) aligns trimmed reads to a reference genome with bowtie2/v.2.4.1 (RRID:SCR_016368) [112], (4) calculates nonduplicate fragment endpoints for each cell, (5) calls peaks using MACS2/v.2.2.7.1 (RRID:SCR_013291) [113, 114] and merges peaks across samples to create a merged BED file, (6) tabulates reads from merged peaks and annotated TSSs (±1 Kb around each TSS) for quality control (QC), (7) separate cell barcodes from background barcodes by fitting a mixture of two negative binomials (noise vs. signal), and (8) assembles sparse matrix tabulating reads per cell barcode falling within the master set of peaks and within gene bodies extended by 2 Kb upstream. We used the rhesus macaque reference genome (Mmul_10) [97] and annotation, obtained from Ensembl (version 101), and merged peaks across all samples (encompassing five library preparation and sequencing batches) to create a global set of peaks. After binarizing UMI counts, we filtered the peak-by-nucleus matrix to include only nuclei with ≥ 1000 binarized UMI, less than 100,000 binarized UMI, and ≥ 30% fraction of reads in peaks (FRIP) (**fig. S11**).

We identified and removed doublets using a similar iterative clustering approach to that described for our single-nucleus RNA dataset (**fig. S11**). Briefly, we ran Scrublet/v.0.2.3 [99] on each sample individually and marked doublets using both a universal threshold (Scrublet doublet score > 0.20) and a per-sample threshold determined by Scrublet and checked and adjusted (if necessary) by eye. We then performed a similar preprocessing, dimensionality reduction, and clustering pipeline to identify clusters with relatively high Scrublet doublet scores (mean Scrublet doublet score > 0.15). We finally removed all nuclei marked as doublets based on the described criteria before concatenating all singlet nuclei across all samples together.

Our snATAC-seq preprocessing, dimensionality reduction, and clustering pipeline likewise tracked closely to our single-nucleus RNA-seq analysis, with minor modifications to accommodate best practices for ATAC-seq data. Briefly, we filtered the data to remove peaks that were not accessible in a minimum of 5 cells as well as peaks that were located on non-autosomal or unplaced scaffolds in the macaque genome. We then filtered the data to the top 100,000 variable features. We performed latent semantic analysis (LSI) on the resulting peak-by-cell matrix to reduce the dimensionality of the data. We performed term frequency/inverse document frequency (TF-IDF) normalization followed by singular value decomposition (SVD) following previously described procedures [45] to reduce the data to 50 PCA dimensions. L2 normalization was then performed on the last 49 principal components, thereby excluding the first principal component, which tends to capture read depth [45]. TF-IDF, SVD, and L2-normalization procedures were implemented using scikit-learn/v.1.0.2 (RRID:SCR_002577) [115]. The L2-normalized PCA matrix was then reduced further and clustered using an identical BBKNN/UMAP/Monocle3 approach to that used for our single-nucleus RNA-seq data. Doublet-derived clusters were also marked for removal using an identical threshold (mean Scrublet doublet score > 0.15).

After marking and removing doublets from our data, we repeated our preprocessing, dimensionality reduction, and clustering pipeline. After observing clear separation of distinct cell classes, we used MUON/v.0.1.2 (RRID:SCR_022804) [116] to calculate promoter accessibility scores by tabulating binarized UMI counts within the region 2,000 bp upstream of a transcriptional start site (TSS). Because at the time of this analysis MUON did not factor in DNA strand information, we ran the function ‘count_fragments_features‘ separately for + and – strand genes, using the “upstream_bp” or “downstream_ bp” arguments as necessary to tabulate counts in the correct upstream region (extending from the TSS to [TSS – 2,000 bp] or [TSS + 2,000 bp], respectively) (https://github.com/scverse/muon/issues/59). We used Scanpy to normalize and visualize resulting promoter accessibility scores (**Fig. 3B**). We provisionally classified nuclei based on promoter accessibility scores of known marker genes.

### Integration of snRNA-seq and snATAC-seq data

We used Graph Linked Unified Embedding (GLUE) implemented in scglue/v.0.2.3 (RRID:SCR_022803) [47] to integrate our snRNA-seq and snATAC-seq datasets. To run scglue, we followed preprocessing procedures in Scanpy recommended by the scglue authors for both our snRNA-seq and snATAC-seq data, after filtering out doublets as described above. For snRNA-seq data, we identified the top 2,000 most variable genes, then normalized, log-transformed, and scaled the data using default parameters in Scanpy. We then reduced the dimensionality of the data to the top 100 principal components using PCA, based on the top 2,000 variable genes and the automatic SVD solver selected by Scanpy. For snATAC-seq data, we used the LSI implementation in scglue to reduce the data to the top 100 principal components, with the number of power iterations set to 15.

We then used scglue to compute a prior guidance graph and propagated highly variable snRNA-seq features (genes) to identify highly variable snATAC-seq features (peaks) based on the guidance graph. We then built and trained the GLUE integration model using the PCA and LSI embeddings, respectively, as the first encoding transformation, modeling raw counts of both snRNA-seq and snATAC-seq data using the negative binomial model, using the batch-correction option to correct for sequencing batches, and using the previously computed prior guidance graph as input. As all nuclei from this study were included (totaling over 4 million nuclei), this analysis was particularly computationally demanding. We performed this analysis on a machine with 1.5 TB RAM, accelerated by 4 Tesla V100 (NVIDIA) GPUs.

After training a GLUE model, we validated effective integration by calculating integration consistency scores using scglue (**fig. S12A**). We then calculated integrated cell and feature embeddings for both snRNA-seq and snATAC-seq data using scglue. After projecting all cells to a unified embedding, we performed UMAP dimensionality reduction using the same procedures as described previously, with one exception. Because the unified GLUE embedding was already batch-corrected, we computed the neighborhood graph using the Scanpy ‘neighbors‘ function rather than BBKNN, with n_neighbors=15.

To transfer cell-class labels from our snRNA-seq data to our snATAC-seq data, we used the ‘transfer_labels‘ function in scglue, which computes shared nearest neighbors between reference (snRNA-seq) and query (snATAC-seq) nuclei, weighted by the Jaccard index. Jaccard indices are then normalized per query nucleus to form a mapping matrix, which is then multiplied by one-hot-encoded reference labels. The reference label with the highest score is then assigned as the predicted cell type, with the highest score retained as the confidence score. For label transfer, because a subset of our snRNA-seq data was derived from samples that were unprofiled in our snATAC-seq data, we limited our reference RNA-seq dataset to only those nuclei deriving from samples profiled in both snRNA-seq and snATAC-seq experiments. We then retained 100,000 nuclei from withheld (unmatched) snRNA-seq samples as a query dataset to evaluate label transfer accuracy. For snATAC-seq label transfer, we used all snATAC-seq nuclei as a query dataset. We used previously assigned parent cell types for our snRNA-seq dataset as reference labels. For our snATAC-seq query nuclei, we retained all predicted cell-class labels with a label transfer confidence score ≥ 0.95. At this threshold, the error rate in our evaluation dataset was 0.43% (**fig. S12B**).

### Identification of candidate *cis*-regulatory elements

To scan for candidate *cis*-regulatory elements underlying differential expression among brain cells, we used two complementary approaches. First, we used the integrative GLUE regulatory inference approach implemented in scglue/v.2.0.3 [47], which calculates regulatory scores based on the cosine similarities between multi-omics data features in an integrated space. Second, we used a metacell approach to construct multi-omic samples (determined via k-means clustering in integrated space) with aggregated (pseudobulk) gene expression and chromatin accessibility counts, which we then modeled using logistic regression. Finally, we calculated differentially accessible peaks using a similar workflow to our snRNA-seq marker gene analysis.

To calculate GLUE regulatory scores, we performed a second integration of our snRNA-seq and snATAC-seq datasets, following an identical pipeline except including the top 6,000 most variable genes (rather than 2,000). This allowed us to identify putative gene:peak regulatory connections and to generate an integrated feature embedding for a greater number of genes and genomic regions. We constructed a window graph between inferred promoters—which we calculated as the region from the strand-specific transcription start site extended upstream 2,000 bp—and peaks using the ‘window_graph‘ function, with the window size set to 150 Kb and a distance-decaying weight, as recommended by the scglue authors. We then used the previously computed window graph and feature embeddings to perform the regulatory inference analysis using the ‘regulatory_ inference‘ function, with the alternative hypothesis set to “greater” in order to perform a one-sided test.

In order to determine the directionality of putative regulatory relationships, we used a second approach based on metacell identification and logistic regression (**Fig. 5A**). We use the ‘get_metacells‘ function to generate multi-omic (snRNA-seq/snATAC-seq) metacells based on k-means clustering of their integrated cell embeddings. As our snRNA-seq dataset included 2,583,967 single-cell transcriptomes, we set k (n_meta) to 10,335 in order to target a mean of roughly 250 RNA transcriptomes per metacell. After identifying metacells in this manner, we summed (pseudobulked) gene expression per metacell. For each gene:peak pair tested in our GLUE regulatory inference, we then performed a logistic regression modeling accessibility of each individual snATAC-seq cell in a given metacell (1: open, 0: closed) as a function of log_2_CPM-normalized gene expression for that snATAC-seq cell’s respective metacell. Logistic regressions were performed in R/v.4.0.2.

We considered candidate *cis*-regulatory relationships to be gene:peak pairs for which FDR-adjusted *P* < 0.05 for both the GLUE regulatory inference and metacell-based logistic regression tests. We classified candidate *cis*-regulatory relationships as positive or negative relationships based on the sign of their logistic regression coefficients (*β* values) (**fig. S15**).

For all peaks, we also tested for marker peaks (peaks with differentially accessibility) using logistic-regression and *t*-test marker-gene methods implemented via the ‘rank_genes_groups‘ function in Scanpy. Similar to our snRNA-seq marker genes analysis, we ran marker peak tests by testing chromatin accessibility in a given cell type against accessibility in all other cells in our dataset. Additionally, to validate marker peaks, we used a second logistic-regression approach implemented via the ‘FindMarkers‘ function in Seurat/v.4.1.1 (RRID:SCR_016341) [117]. In contrast to the logistic regression in Scanpy, the Seurat implementation is not a regularized procedure and is thus able to control for latent variables and to calculate *P* values. To reduce computational burden, we ran ‘FindMarkers‘ on a dataset with 1,000 cells per cell class. As we found that output statistics (regularized logistic regression coefficient in Scanpy and log foldchange in Seurat) were highly concordant (**fig. S16**), we report Scanpy results here as they included all possible cells. We considered peaks to be differentially accessible if the regularized logistic regression coefficient > 0, the log fold change > 0, and the *t*-test *P_adj_* < 0.05.

### snATAC-seq cell subtype analysis

To mitigate peak-calling biases while allowing us to probe more nuanced regulatory variation within cell populations, we called a new set of cell-class-specific peaks for each cell type with assigned cells, skipping rarer cell types for which no snATAC-seq nuclei passed our prediction threshold above.

Following scglue integration and assignment of snATAC-seq cells to cell classes, we created cell-class-specific pseudobulk epigenomes by aggregating all nonduplicate fragment endpoints for each cell class. These cell-class-level ATAC-seq data were then used for peak-calling using MACS3/v.3.0.0a6 [113, 114], with the same peak calling parameters that we used for each sample and batch described in the “snATAC-seq pre-processing” section above (‘-g 2.7e9 –call-summits – nomodel‘). For each cell class, we repeated steps from our snATAC-seq data generation pipeline to tabulate reads from newly called peaks and to assemble sparse count matrices matrix tabulating reads per cell barcode falling within the master set of peaks and within gene bodies extended by 2 Kb upstream. We then imported peak-by-nucleus count matrices into the AnnData/v.0.8.0 [98] framework.

To assign cell subtypes for our snATAC-seq data (**fig. S13**), we repeated preprocessing, data integration, label transfer, and regulatory inference procedures described above on each cell class individually. In contrast to our global joint analysis, we only included snRNA-seq nuclei deriving from samples that were profiled in both snRNA-seq and snATAC-seq experiments, and used the top 6,000 most variable genes in our snRNA-seq analysis, and used the snATAC-seq peak sets specific to each cell type. The remainder of our preprocessing and data integration procedures followed the same pipeline described previously for our global integration analysis. For label transfer, we also followed largely the same procedures as for our global label transfer pipeline. We did not, however, use a label transfer confidence score threshold under the assumption that snATAC-seq nuclei would, on average, be assigned to the correct cell subtype and, if incorrect, would be assigned to a closely related cell subtype (i.e., a neighboring subtype in the integrated multidimensional cell space).

For metacell-based regulatory inference, we varied the settings for k based on dataset size in order to target a mean of 50 transcriptomes per metacell.

### Transcription factor binding site enrichment

For enrichment analyses at the cell-class level, we focused on peaks that were deemed accessible in one and only one cell class, which we called “cell-class-unique peaks”. We identified these peaks using BedTools/v.2.30.0 (‘intersect -v‘) (RRID:SCR_006646) [118] to find all peaks in a cell class that did not overlap with any peak called in another cell class. The number of peaks identified in this manner ranged from 655 (in ependymal cells) to 71,049 (in glutamatergic neurons). We tested for enrichment of TF binding motifs in cellclass-unique peaks compared to the background of the rhesus macaque genome while controlling for GC content, implemented in the monaLisa/v.1.3.1 (RRID:SCR_ 022802) [119] in R/v.4.1.0 (**table S8**). We used the JASPAR 2018 (RRID:SCR_003030) non-redundant vertebrate core position weight matrices [120].

At the cell-subtype level, we tested for enrichment using the top differentially accessible peaks among sub-types of the same cell class, excluding peaks with regularized logistic regression coefficients < 0 (**table S9**). We retained the top first percentile of marker peaks, ranked according to their regularized logistic regression coefficients.

### Disease heritability enrichment

We calculated enrichment of disease-associated variants in cell-class-specific accessible chromatin regions using linkage disequilibrium score regression, LDSC (RRID:SCR_022801) [73, 74] (**Fig. 6** and **table S15**). Because the trait-associated loci are annotated in the human genome, we converted all peaks (at the combined level as well as each individual cell-class level) from the rhesus macaque genome coordinates to GRCh37 using UCSC’s liftOver/v.302 (RRID:SCR_018160) tool [121]. We followed the standard pipeline using the 1000 Genomes baseline model and precomputed .sumstats files. A list of phenotypes tested can be found in **table S14**.

## References

[1] Dunbar, R. I. M. and Shultz, S. Evolution in the social brain. Science 317(5843), 1344–1347 (2007).

[2] Navarrete, A., van Schaik, C. P., and Isler, K. Energetics and the evolution of human brain size. Nature 480(7375), 91–93 (2011).

[3] Darwin, C. The Descent of Man, and Selection in Relation to Sex. John Murray, London, UK (1871).

[4] Herculano-Houzel, S., Collins, C. E., Wong, P., and Kaas, J. H. Cellular scaling rules for primate brains. Proc. Natl. Acad. Sci. U. S. A. 104(9), 3562–3567 (2007).

[5] Grasby, K. L., Jahanshad, N., Painter, J. N., Colodro-Conde, L., Bralten, J., Hibar, D. P., Lind, P. A., Pizzagalli, F., Ching, C. R. K., McMahon, M. A. B., Shatokhina, N., Zsembik, L. C. P., Thomopoulos, S. I., Zhu, A. H., Strike, L. T., Agartz, I., Alhusaini, S., Almeida, M. A. A., Alnæs, D., Amlien, I. K., Andersson, M., Ard, T., Armstrong, N. J., Ashley-Koch, A., Atkins, J. R., Bernard, M., Brouwer, R. M., Buimer, E. E. L., Bülow, R., Bürger, C., Cannon, D. M., Chakravarty, M., Chen, Q., Cheung, J. W., Couvy-Duchesne, B., Dale, A. M., Dalvie, S., de Araujo, T. K., de Zubicaray, G. I., de Zwarte, S. M. C., den Braber, A., Doan, N. T., Dohm, K., Ehrlich, S., Engelbrecht, H.-R., Erk, S., Fan, C. C., Fedko, I. O., Foley, S. F., Ford, J. M., Fukunaga, M., Garrett, M. E., Ge, T., Giddaluru, S., Goldman, A. L., Green, M. J., Groenewold, N. A., Grotegerd, D., Gurholt, T. P., Gutman, B. A., Hansell, N. K., Harris, M. A., Harrison, M. B., Haswell, C. C., Hauser, M., Herms, S., Heslenfeld, D. J., Ho, N. F., Hoehn, D., Hoffmann, P., Holleran, L., Hoogman, M., Hottenga, J.-J., Ikeda, M., Janowitz, D., Jansen, I. E., Jia, T., Jockwitz, C., Kanai, R., Karama, S., Kasperaviciute, D., Kaufmann, T., Kelly, S., Kikuchi, M., Klein, M., Knapp, M., Knodt, A. R., Krämer, B., Lam, M., Lancaster, T. M., Lee, P. H., Lett, T. A., Lewis, L. B., Lopes-Cendes, I., Luciano, M., Macciardi, F., Marquand, A. F., Mathias, S. R., Melzer, T. R., Milaneschi, Y., Mirza-Schreiber, N., Moreira, J. C. V., Mühleisen, T. W., Müller-Myhsok, B., Najt, P., Nakahara, S., Nho, K., Olde Loohuis, L. M., Orfanos, D. P., Pearson, J. F., Pitcher, T. L., Pütz, B., Quidé, Y., Ragothaman, A., Rashid, F. M., Reay, W. R., Redlich, R., Reinbold, C. S., Repple, J., Richard, G., Riedel, B. C., Risacher, S. L., Rocha, C. S., Mota, N. R., Salminen, L., Saremi, A., Saykin, A. J., Schlag, F., Schmaal, L., Schofield, P. R., Secolin, R., Shapland, C. Y., Shen, L., Shin, J., Shumskaya, E., Sønderby, I. E., Sprooten, E., Tansey, K. E., Teumer, A., Thalamuthu, A., Tordesillas-Gutiérrez, D., Turner, J. A., Uhlmann, A., Vallerga, C. L., van der Meer, D., van Donkelaar, M. M. J., van Eijk, L., van Erp, T. G. M., van Haren, N. E. M., van Rooij, D., van Tol, M.-J., Veldink, J. H., Verhoef, E., Walton, E., Wang, M., Wang, Y., Wardlaw, J. M., Wen, W., Westlye, L. T., Whelan, C. D., Witt, S. H., Wittfeld, K., Wolf, C., Wolfers, T., Wu, J. Q., Yasuda, C. L., Zaremba, D., Zhang, Z., Zwiers, M. P., Artiges, E., Assareh, A. A., Ayesa-Arriola, R., Belger, A., Brandt, C. L., Brown, G. G., Cichon, S., Curran, J. E., Davies, G. E., Degenhardt, F., Dennis, M. F., Dietsche, B., Djurovic, S., Doherty, C. P., Espiritu, R., Garijo, D., Gil, Y., Gowland, P. A., Green, R. C., Häusler, A. N., Heindel, W., Ho, B.-C., Hoffmann, W. U., Holsboer, F., Homuth, G., Hosten, N., Jack, Jr, C. R., Jang, M., Jansen, A., Kimbrel, N. A., Kolskår, K., Koops, S., Krug, A., Lim, K. O., Luykx, J. J., Mathalon, D. H., Mather, K. A., Mattay, V. S., Matthews, S., Mayoral Van Son, J., McEwen, S. C., Melle, I., Morris, D. W., Mueller, B. A., Nauck, M., Nordvik, J. E., Nöthen, M. M., O’Leary, D. S., Opel, N., Martinot, M.-L. P., Pike, G. B., Preda, A., Quinlan, E. B., Rasser, P. E., Ratnakar, V., Reppermund, S., Steen, V. M., Tooney, P. A., Torres, F. R., Veltman, D. J., Voyvodic, J. T., Whelan, R., White, T., Yamamori, H., Adams, H. H. H., Bis, J. C., Debette, S., Decarli, C., Fornage, M., Gudnason, V., Hofer, E., Ikram, M. A., Launer, L., Longstreth, W. T., Lopez, O. L., Mazoyer, B., Mosley, T. H., Roshchupkin, G. V., Satizabal, C. L., Schmidt, R., Seshadri, S., Yang, Q., Alzheimer’s Disease Neuroimaging Initiative, CHARGE Consortium, EPIGEN Consortium, IMAGEN Consortium, SYS Consortium, Parkinson’s Progression Markers Initiative, Alvim, M. K. M., Ames, D., Anderson, T. J., Andreassen, O. A., Arias-Vasquez, A., Bastin, M. E., Baune, B. T., Beckham, J. C., Blangero, J., Boomsma, D. I., Brodaty, H., Brunner, H. G., Buckner, R. L., Buitelaar, J. K., Bustillo, J. R., Cahn, W., Cairns, M. J., Calhoun, V., Carr, V. J., Caseras, X., Caspers, S., Cavalleri, G. L., Cendes, F., Corvin, A., Crespo-Facorro, B., Dalrymple-Alford, J. C., Dannlowski, U., de Geus, E. J. C., Deary, I. J., Delanty, N., Depondt, C., Desrivières, S., Donohoe, G., Espeseth, T., Fernández, G., Fisher, S. E., Flor, H., Forstner, A. J., Francks, C., Franke, B., Glahn, D. C., Gollub, R. L., Grabe, H. J., Gruber, O., Håberg, A. K., Hariri, A. R., Hartman, C. A., Hashimoto, R., Heinz, A., Henskens, F. A., Hillegers, M. H. J., Hoekstra, P. J., Holmes, A. J., Hong, L. E., Hopkins, W. D., Hulshoff Pol, H. E., Jernigan, T. L., Jönsson, E. G., Kahn, R. S., Kennedy, M. A., Kircher, T. T. J., Kochunov, P., Kwok, J. B. J., Le Hellard, S., Loughland, C. M., Martin, N. G., Martinot, J.-L., McDonald, C., McMahon, K. L., Meyer-Lindenberg, A., Michie, P. T., Morey, R. A., Mowry, B., Nyberg, L., Oosterlaan, J., Ophoff, R. A., Pantelis, C., Paus, T., Pausova, Z., Penninx, B. W. J. H., Polderman, T. J. C., Posthuma, D., Rietschel, M., Roffman, J. L., Rowland, L. M., Sachdev, P. S., Sämann, P. G., Schall, U., Schumann, G., Scott, R. J., Sim, K., Sisodiya, S. M., Smoller, J. W., Sommer, I. E., St Pourcain, B., Stein, D. J., Toga, A. W., Trollor, J. N., Van der Wee, N. J. A., van ’t Ent, D., Völzke, H., Walter, H., Weber, B., Weinberger, D. R., Wright, M. J., Zhou, J., Stein, J. L., Thompson, P. M., Medland, S. E., and Enhancing NeuroImaging Genetics through Meta-Analysis Consortium (ENIGMA)—Genetics working group. The genetic architecture of the human cerebral cortex. Science 367(6484), eaay6690 (2020).

[6] Zeng, J., Konopka, G., Hunt, B. G., Preuss, T. M., Geschwind, D., and Yi, S. V. Divergent wholegenome methylation maps of human and chimpanzee brains reveal epigenetic basis of human regulatory evolution. Am. J. Hum. Genet. 91(3), 455–465 (2012).

[7] Preuss, T. M. Human brain evolution: from gene discovery to phenotype discovery. Proc. Natl. Acad. Sci. U. S. A. 109 Suppl 1, 10709–10716 (2012).

[8] Cajal, S. R. Y. The Croonian lecture: la fine structure des centres nerveux. Proc. R. Soc. Lond. 55(331-335), 444–468 (1894).

[9] Striedter, G. F. Principles of Brain Evolution. Sinauer Associates Incorporated, (2005).

[10] Allman, J. M., Watson, K. K., Tetreault, N. A., and Hakeem, A. Y. Intuition and autism: a possible role for Von Economo neurons. Trends Cogn. Sci. 9(8), 367–373 (2005).

[11] Rizzolatti, G., Fabbri-Destro, M., and Cattaneo, L. Mirror neurons and their clinical relevance. Nat. Clin. Pract. Neurol. 5(1), 24–34 (2009).

[12] Boldog, E., Bakken, T. E., Hodge, R. D., Novotny, M., Aevermann, B. D., Baka, J., Bordé, S., Close, J. L., Diez-Fuertes, F., Ding, S.-L., Faragó, N., Kocsis, Á. K., Kovács, B., Maltzer, Z., McCorrison, J. M., Miller, J. A., Molnár, G., Oláh, G., Ozsvár, A., Rózsa, M., Shehata, S. I., Smith, K. A., Sunkin, S. M., Tran, D. N., Venepally, P., Wall, A., Puskás, L. G., Barzó, P., Steemers, F. J., Schork, N. J., Scheuermann, R. H., Lasken, R. S., Lein, E. S., and Tamás, G. Transcriptomic and morphophysiological evidence for a specialized human cortical GABAergic cell type. Nat. Neurosci. 21(9), 1185–1195 (2018).

[13] Nowakowski, T. J., Bhaduri, A., Pollen, A. A., Alvarado, B., Mostajo-Radji, M. A., Di Lullo, E., Haeussler, M., Sandoval-Espinosa, C., Liu, S. J., Velmeshev, D., Ounadjela, J. R., Shuga, J., Wang, X., Lim, D. A., West, J. A., Leyrat, A. A., Kent, W. J., and Kriegstein, A. R. Spatiotemporal gene expression trajectories reveal developmental hierarchies of the human cortex. Science 358(6368), 1318–1323 (2017).

[14] Hodge, R. D., Bakken, T. E., Miller, J. A., Smith, K. A., Barkan, E. R., Graybuck, L. T., Close, J. L., Long, B., Johansen, N., Penn, O., Yao, Z., Eggermont, J., Höllt, T., Levi, B. P., Shehata, S. I., Aevermann, B., Beller, A., Bertagnolli, D., Brouner, K., Casper, T., Cobbs, C., Dalley, R., Dee, N., Ding, S.-L., Ellenbogen, R. G., Fong, O., Garren, E., Goldy, J., Gwinn, R. P., Hirschstein, D., Keene, C. D., Keshk, M., Ko, A. L., Lathia, K., Mahfouz, A., Maltzer, Z., McGraw, M., Nguyen, T. N., Nyhus, J., Ojemann, J. G., Oldre, A., Parry, S., Reynolds, S., Rimorin, C., Shapovalova, N. V., Somasundaram, S., Szafer, A., Thomsen, E. R., Tieu, M., Quon, G., Scheuermann, R. H., Yuste, R., Sunkin, S. M., Lelieveldt, B., Feng, D., Ng, L., Bernard, A., Hawrylycz, M., Phillips, J. W., Tasic, B., Zeng, H., Jones, A. R., Koch, C., and Lein, E. S. Conserved cell types with divergent features in human versus mouse cortex. Nature 573(7772), 61–68 (2019).

[15] Bakken, T. E., Jorstad, N. L., Hu, Q., Lake, B. B., Tian, W., Kalmbach, B. E., Crow, M., Hodge, R. D., Krienen, F. M., Sorensen, S. A., Eggermont, J., Yao, Z., Aevermann, B. D., Aldridge, A. I., Bartlett, A., Bertagnolli, D., Casper, T., Castanon, R. G., Crichton, K., Daigle, T. L., Dalley, R., Dee, N., Dembrow, N., Diep, D., Ding, S.-L., Dong, W., Fang, R., Fischer, S., Goldman, M., Goldy, J., Graybuck, L. T., Herb, B. R., Hou, X., Kancherla, J., Kroll, M., Lathia, K., van Lew, B., Li, Y. E., Liu, C. S., Liu, H., Lucero, J. D., Mahurkar, A., McMillen, D., Miller, J. A., Moussa, M., Nery, J. R., Nicovich, P. R., Niu, S.-Y., Orvis, J., Osteen, J. K., Owen, S., Palmer, C. R., Pham, T., Plongthongkum, N., Poirion, O., Reed, N. M., Rimorin, C., Rivkin, A., Romanow, W. J., Sedeño-Cortés, A. E., Siletti, K., Somasundaram, S., Sulc, J., Tieu, M., Torkelson, A., Tung, H., Wang, X., Xie, F., Yanny, A. M., Zhang, R., Ament, S. A., Behrens, M. M., Bravo, H. C., Chun, J., Dobin, A., Gillis, J., Hertzano, R., Hof, P. R., Höllt, T., Horwitz, G. D., Keene, C. D., Kharchenko, P. V., Ko, A. L., Lelieveldt, B. P., Luo, C., Mukamel, E. A., Pinto-Duarte, A., Preissl, S., Regev, A., Ren, B., Scheuermann, R. H., Smith, K., Spain, W. J., White, O. R., Koch, C., Hawrylycz, M., Tasic, B., Macosko, E. Z., McCarroll, S. A., Ting, J. T., Zeng, H., Zhang, K., Feng, G., Ecker, J. R., Linnarsson, S., and Lein, E. S. Comparative cellular analysis of motor cortex in human, marmoset and mouse. Nature 598(7879), 111–119 (2021).

[16] Ecker, J. R., Geschwind, D. H., Kriegstein, A. R., Ngai, J., Osten, P., Polioudakis, D., Regev, A., Sestan, N., Wickersham, I. R., and Zeng, H. The BRAIN Initiative Cell Census Consortium: lessons learned toward generating a comprehensive brain cell atlas. Neuron 96(3), 542–557 (2017).

[17] Gibbs, R. A., Rogers, J., Katze, M. G., Bumgarner, R., Weinstock, G. M., Mardis, E. R., Remington, K. A., Strausberg, R. L., Venter, J. C., Wilson, R. K., Batzer, M. A., Bustamante, C. D., Eichler, E. E., Hahn, M. W., Hardison, R. C., Makova, K. D., Miller, W., Milosavljevic, A., Palermo, R. E., Siepel, A., Sikela, J. M., Attaway, T., Bell, S., Bernard, K. E., Buhay, C. J., Chandrabose, M. N., Dao, M., Davis, C., Delehaunty, K. D., Ding, Y., Dinh, H. H., Dugan-Rocha, S., Fulton, L. A., Gabisi, R. A., Garner, T. T., Godfrey, J., Hawes, A. C., Hernandez, J., Hines, S., Holder, M., Hume, J., Jhangiani, S. N., Joshi, V., Khan, Z. M., Kirkness, E. F., Cree, A., Fowler, R. G., Lee, S., Lewis, L. R., Li, Z., Liu, Y.-S., Moore, S. M., Muzny, D., Nazareth, L. V., Ngo, D. N., Okwuonu, G. O., Pai, G., Parker, D., Paul, H. A., Pfannkoch, C., Pohl, C. S., Rogers, Y.-H., Ruiz, S. J., Sabo, A., Santibanez, J., Schneider, B. W., Smith, S. M., Sodergren, E., Svatek, A. F., Utterback, T. R., Vattathil, S., Warren, W., White, C. S., Chinwalla, A. T., Feng, Y., Halpern, A. L., Hillier, L. W., Huang, X., Minx, P., Nelson, J. O., Pepin, K. H., Qin, X., Sutton, G. G., Venter, E., Walenz, B. P., Wallis, J. W., Worley, K. C., Yang, S.-P., Jones, S. M., Marra, M. A., Rocchi, M., Schein, J. E., Baertsch, R., Clarke, L., Csürös, M., Glasscock, J., Harris, R. A., Havlak, P., Jackson, A. R., Jiang, H., Liu, Y., Messina, D. N., Shen, Y., Song, H. X.-Z., Wylie, T., Zhang, L., Birney, E., Han, K., Konkel, M. K., Lee, J., Smit, A. F. A., Ullmer, B., Wang, H., Xing, J., Burhans, R., Cheng, Z., Karro, J. E., Ma, J., Raney, B., She, X., Cox, M. J., Demuth, J. P., Dumas, L. J., Han, S.-G., Hopkins, J., Karimpour-Fard, A., Kim, Y. H., Pollack, J. R., Vinar, T., Addo-Quaye, C., Degenhardt, J., Denby, A., Hubisz, M. J., Indap, A., Kosiol, C., Lahn, B. T., Lawson, H. A., Marklein, A., Nielsen, R., Vallender, E. J., Clark, A. G., Ferguson, B., Hernandez, R. D., Hirani, K., Kehrer-Sawatzki, H., Kolb, J., Patil, S., Pu, L.-L., Ren, Y., Smith, D. G., Wheeler, D. A., Schenck, I., Ball, E. V., Chen, R., Cooper, D. N., Giardine, B., Hsu, F., Kent, W. J., Lesk, A., Nelson, D. L., O’brien, W. E., Prüfer, K., Stenson, P. D., Wallace, J. C., Ke, H., Liu, X.-M., Wang, P., Xiang, A. P., Yang, F., Barber, G. P., Haussler, D., Karolchik, D., Kern, A. D., Kuhn, R. M., Smith, K. E., Zwieg, A. S., and Rhesus Macaque Genome Sequencing and Analysis Consortium. Evolutionary and biomedical insights from the rhesus macaque genome. Science 316(5822), 222–234 (2007).

[18] Cao, J., Packer, J. S., Ramani, V., Cusanovich, D. A., Huynh, C., Daza, R., Qiu, X., Lee, C., Furlan, S. N., Steemers, F. J., Adey, A., Waterston, R. H., Trapnell, C., and Shendure, J. Comprehensive single-cell transcriptional profiling of a multicellular organism. Science 357(6352), 661–667 (2017).

[19] Cao, J., Spielmann, M., Qiu, X., Huang, X., Ibrahim, D. M., Hill, A. J., Zhang, F., Mundlos, S., Christiansen, L., Steemers, F. J., Trapnell, C., and Shendure, J. The single-cell transcriptional landscape of mammalian organogenesis. Nature 566(7745), 496–502 (2019).

[20] Cao, J. and Shendure, J. sci-RNA-seq3. protocols.io (2020).

[21] Martin, B. K., Qiu, C., Nichols, E., Phung, M., Green-Gladden, R., Srivatsan, S., Blecher-Gonen, R., Beliveau, B. J., Trapnell, C., Cao, J., and Shendure, J. An optimized protocol for single cell transcriptional profiling by combinatorial indexing. (2021).

[22] Kamath, T., Abdulraouf, A., Burris, S. J., Langlieb, J., Gazestani, V., Nadaf, N. M., Balderrama, K., Vanderburg, C., and Macosko, E. Z. Single-cell genomic profiling of human dopamine neurons identifies a population that selectively degenerates in Parkinson’s disease. Nat. Neurosci. 25(5), 588–595 (2022).

[23] Ren, J., Isakova, A., Friedmann, D., Zeng, J., Grutzner, S. M., Pun, A., Zhao, G. Q., Kolluru, S. S., Wang, R., Lin, R., Li, P., Li, A., Raymond, J. L., Luo, Q., Luo, M., Quake, S. R., and Luo, L. Single-cell transcriptomes and whole-brain projections of serotonin neurons in the mouse dorsal and median raphe nuclei. eLife 8, e49424 (2019).

[24] Poewe, W., Seppi, K., Tanner, C. M., Halliday, G. M., Brundin, P., Volkmann, J., Schrag, A.-E., and Lang, A. E. Parkinson disease. Nat Rev Dis Primers 3, 17013 (2017).

[25] Fakhoury, M. Revisiting the Serotonin Hypothesis: implications for major depressive disorders. Mol. Neurobiol. 53(5), 2778–2786 (2016).

[26] Yang, A. C., Vest, R. T., Kern, F., Lee, D. P., Agam, M., Maat, C. A., Losada, P. M., Chen, M. B., Schaum, N., Khoury, N., Toland, A., Calcuttawala, K., Shin, H., Pálovics, R., Shin, A., Wang, E. Y., Luo, J., Gate, D., Schulz-Schaeffer, W. J., Chu, P., Siegenthaler, J. A., McNerney, M. W., Keller, A., and Wyss-Coray, T. A human brain vascular atlas reveals diverse mediators of Alzheimer’s risk. Nature 603(7903), 885–892 (2022).

[27] Hao, Z.-Z., Wei, J.-R., Xiao, D., Liu, R., Xu, N., Tang, L., Huang, M., Shen, Y., Xing, C., Huang, W., Liu, X., Xiang, M., Liu, Y., Miao, Z., and Liu, S. Single-cell transcriptomics of adult macaque hippocampus reveals neural precursor cell populations. Nat. Neurosci. 25(6), 805–817 (2022).

[28] Cusanovich, D. A., Hill, A. J., Aghamirzaie, D., Daza, R. M., Pliner, H. A., Berletch, J. B., Filippova, G. N., Huang, X., Christiansen, L., DeWitt, W. S., Lee, C., Regalado, S. G., Read, D. F., Steemers, F. J., Disteche, C. M., Trapnell, C., and Shendure, J. A single-cell atlas of in vivo mammalian chromatin accessibility. Cell 174(5), 1309–1324.e18 (2018).

[29] Li, Y. E., Preissl, S., Hou, X., Zhang, Z., Zhang, K., Qiu, Y., Poirion, O. B., Li, B., Chiou, J., Liu, H., Pinto-Duarte, A., Kubo, N., Yang, X., Fang, R., Wang, X., Han, J. Y., Lucero, J., Yan, Y., Miller, M., Kuan, S., Gorkin, D., Gaulton, K. J., Shen, Y., Nunn, M., Mukamel, E. A., Behrens, M. M., Ecker, J. R., and Ren, B. An atlas of gene regulatory elements in adult mouse cerebrum. Nature 598(7879), 129–136 (2021).

[30] Berg, J., Sorensen, S. A., Ting, J. T., Miller, J. A., Chartrand, T., Buchin, A., Bakken, T. E., Budzillo, A., Dee, N., Ding, S.-L., Gouwens, N. W., Hodge, R. D., Kalmbach, B., Lee, C., Lee, B. R., Alfiler, L., Baker, K., Barkan, E., Beller, A., Berry, K., Bertagnolli, D., Bickley, K., Bomben, J., Braun, T., Brouner, K., Casper, T., Chong, P., Crichton, K., Dalley, R., de Frates, R., Desta, T., Lee, S. D., D’Orazi, F., Dotson, N., Egdorf, T., Enstrom, R., Farrell, C., Feng, D., Fong, O., Furdan, S., Galakhova, A. A., Gamlin, C., Gary, A., Glandon, A., Goldy, J., Gorham, M., Goriounova, N. A., Gratiy, S., Graybuck, L., Gu, H., Hadley, K., Hansen, N., Heistek, T. S., Henry, A. M., Heyer, D. B., Hill, D., Hill, C., Hupp, M., Jarsky, T., Kebede, S., Keene, L., Kim, L., Kim, M.-H., Kroll, M., Latimer, C., Levi, B. P., Link, K. E., Mallory, M., Mann, R., Marshall, D., Maxwell, M., McGraw, M., McMillen, D., Melief, E., Mertens, E. J., Mezei, L., Mihut, N., Mok, S., Molnar, G., Mukora, A., Ng, L., Ngo, K., Nicovich, P. R., Nyhus, J., Olah, G., Oldre, A., Omstead, V., Ozsvar, A., Park, D., Peng, H., Pham, T., Pom, C. A., Potekhina, L., Rajanbabu, R., Ransford, S., Reid, D., Rimorin, C., Ruiz, A., Sandman, D., Sulc, J., Sunkin, S. M., Szafer, A., Szemenyei, V., Thomsen, E. R., Tieu, M., Torkelson, A., Trinh, J., Tung, H., Wakeman, W., Waleboer, F., Ward, K., Wilbers, R., Williams, G., Yao, Z., Yoon, J.-G., Anastassiou, C., Arkhipov, A., Barzo, P., Bernard, A., Cobbs, C., de Witt Hamer, P. C., Ellenbogen, R. G., Esposito, L., Ferreira, M., Gwinn, R. P., Hawrylycz, M. J., Hof, P. R., Idema, S., Jones, A. R., Keene, C. D., Ko, A. L., Murphy, G. J., Ng, L., Ojemann, J. G., Patel, A. P., Phillips, J. W., Silbergeld, D. L., Smith, K., Tasic, B., Yuste, R., Segev, I., de Kock, C. P. J., Mansvelder, H. D., Tamas, G., Zeng, H., Koch, C., and Lein, E. S. Human neocortical expansion involves glutamatergic neuron diversification. Nature 598(7879), 151–158 (2021).

[31] Trevino, A. E., Müller, F., Andersen, J., Sundaram, L., Kathiria, A., Shcherbina, A., Farh, K., Chang, H. Y., Paşca, A. M., Kundaje, A., Paşca, S. P., and Greenleaf, W. J. Chromatin and gene-regulatory dynamics of the developing human cerebral cortex at single-cell resolution. Cell 184(19), 5053–5069.e23 (2021).

[32] Krienen, F. M., Goldman, M., Zhang, Q., C H Del Rosario, R., Florio, M., Machold, R., Saunders, A., Levandowski, K., Zaniewski, H., Schuman, B., Wu, C., Lutservitz, A., Mullally, C. D., Reed, N., Bien, E., Bortolin, L., Fernandez-Otero, M., Lin, J. D., Wysoker, A., Nemesh, J., Kulp, D., Burns, M., Tkachev, V., Smith, R., Walsh, C. A., Dimidschstein, J., Rudy, B., S Kean, L., Berretta, S., Fishell, G., Feng, G., and McCarroll, S. A. Innovations present in the primate interneuron repertoire. Nature 586(7828), 262–269 (2020).

[33] Barres, B. A. The mystery and magic of glia: a perspective on their roles in health and disease. Neuron 60(3), 430–440 (2008).

[34] Valori, C. F., Guidotti, G., Brambilla, L., and Rossi, D. Astrocytes: emerging therapeutic targets in neurological disorders. Trends Mol. Med. 25(9), 750–759 (2019).

[35] Wiebe, S., Nagpal, A., Truong, V. T., Park, J., Skalecka, A., He, A. J., Gamache, K., Khoutorsky, A., Gantois, I., and Sonenberg, N. Inhibitory interneurons mediate autism-associated behaviors via 4E-BP2. Proc. Natl. Acad. Sci. U. S. A. 116(36), 18060–18067 (2019).

[36] Miyoshi, G., Ueta, Y., Natsubori, A., Hiraga, K., Osaki, H., Yagasaki, Y., Kishi, Y., Yanagawa, Y., Fishell, G., Machold, R. P., and Miyata, M. FoxG1 regulates the formation of cortical GABAergic circuit during an early postnatal critical period resulting in autism spectrum disorder-like phenotypes. Nat. Commun. 12(1), 3773 (2021).

[37] Bakken, T. E., van Velthoven, C. T., Menon, V., Hodge, R. D., Yao, Z., Nguyen, T. N., Graybuck, L. T., Horwitz, G. D., Bertagnolli, D., Goldy, J., Yanny, A. M., Garren, E., Parry, S., Casper, T., Shehata, S. I., Barkan, E. R., Szafer, A., Levi, B. P., Dee, N., Smith, K. A., Sunkin, S. M., Bernard, A., Phillips, J., Hawrylycz, M. J., Koch, C., Murphy, G. J., Lein, E., Zeng, H., and Tasic, B. Single-cell and single-nucleus RNA-seq uncovers shared and distinct axes of variation in dorsal LGN neurons in mice, non-human primates, and humans. eLife 10, e64875 (2021).

[38] Zhang, Q., Sano, C., Masuda, A., Ando, R., Tanaka, M., and Itohara, S. Netrin-G1 regulates fear-like and anxiety-like behaviors in dissociable neural circuits. Sci. Rep. 6, 28750 (2016).

[39] Mathys, H., Davila-Velderrain, J., Peng, Z., Gao, F., Mohammadi, S., Young, J. Z., Menon, M., He, L., Abdurrob, F., Jiang, X., Martorell, A. J., Ransohoff, R. M., Hafler, B. P., Bennett, D. A., Kellis, M., and Tsai, L.-H. Single-cell transcriptomic analysis of Alzheimer’s disease. Nature 570(7761), 332–337 (2019).

[40] Batiuk, M. Y., Martirosyan, A., Wahis, J., de Vin, F., Marneffe, C., Kusserow, C., Koeppen, J., Viana, J. F., Oliveira, J. F., Voet, T., Ponting, C. P., Belgard, T. G., and Holt, M. G. Identification of regionspecific astrocyte subtypes at single cell resolution. Nat. Commun. 11 (1), 1220 (2020).

[41] Huang, X., Henck, J., Qiu, C., Sreenivasan, V. K. A., Balachandran, S., Behncke, R., Chan, W.-L., Despang, A., Dickel, D. E., Haag, N., Hägerling, R., Hansmeier, N., Hennig, F., Marshall, C., Rajderkar, S., Ringel, A., Robson, M., Saunders, L., Srivatsan, S. R., Ulferts, S., Wittler, L., Zhu, Y., Kalscheuer, V. M., Ibrahim, D., Kurth, I., Kornak, U., Beier, D. R., Visel, A., Pennacchio, L. A., Trapnell, C., Cao, J., Shendure, J., and Spielmann, M. Single cell, whole embryo phenotyping of pleiotropic disorders of mammalian development. bioRxiv, 2022.08.03.500325 (2022).

[42] Kirdajova, D., Valihrach, L., Valny, M., Kriska, J., Krocianova, D., Benesova, S., Abaffy, P., Zucha, D., Klassen, R., Kolenicova, D., Honsa, P., Kubista, M., and Anderova, M. Transient astrocyte-like NG2 glia subpopulation emerges solely following permanent brain ischemia. Glia 69(11), 2658–2681 (2021).

[43] Shu, Q., Lennemann, N. J., Sarkar, S. N., Sadovsky, Y., and Coyne, C. B. ADAP2 Is an Interferon Stimulated Gene That Restricts RNA Virus Entry. PLoS Pathog. 11(9), e1005150 (2015).

[44] Cusanovich, D. A., Daza, R., Adey, A., Pliner, H. A., Christiansen, L., Gunderson, K. L., Steemers, F. J., Trapnell, C., and Shendure, J. Multiplex single cell profiling of chromatin accessibility by combinatorial cellular indexing. Science 348(6237), 910–914 (2015).

[45] Domcke, S., Hill, A. J., Daza, R. M., Cao, J., O’Day, D. R., Pliner, H. A., Aldinger, K. A., Pokholok, D., Zhang, F., Milbank, J. H., Zager, M. A., Glass, I. A., Steemers, F. J., Doherty, D., Trapnell, C., Cusanovich, D. A., and Shendure, J. A human cell atlas of fetal chromatin accessibility. Science 370(6518), eaba7612 (2020).

[46] Stuart, T., Butler, A., Hoffman, P., Hafemeister, C., Papalexi, E., Mauck, 3rd, W. M., Hao, Y., Stoeckius, M., Smibert, P., and Satija, R. Comprehensive integration of single-cell data. Cell 177(7), 1888–1902.e21 (2019).

[47] Cao, Z.-J. and Gao, G. Multi-omics single-cell data integration and regulatory inference with graph-linked embedding. Nat. Biotechnol. (2022).

[48] Ishibashi, M., Ang, S. L., Shiota, K., Nakanishi, S., Kageyama, R., and Guillemot, F. Targeted disruption of mammalian hairy and Enhancer of split homolog-1 (HES-1) leads to up-regulation of neural helix-loop-helix factors, premature neurogenesis, and severe neural tube defects. Genes Dev. 9(24), 3136–3148 (1995).

[49] Collignon, J., Sockanathan, S., Hacker, A., Cohen-Tannoudji, M., Norris, D., Rastan, S., Stevanovic, M., Goodfellow, P. N., and Lovell-Badge, R. A comparison of the properties of Sox-3 with Sry and two related genes, Sox-1 and Sox-2. Development 122(2), 509–520 (1996).

[50] Gaiano, N., Nye, J. S., and Fishell, G. Radial glial identity is promoted by Notch1 signaling in the murine forebrain. Neuron 26(2), 395–404 (2000).

[51] Palma, V. and Ruiz i Altaba, A. Hedgehog-GLI signaling regulates the behavior of cells with stem cell properties in the developing neocortex. Development 131(2), 337–345 (2004).

[52] Kitamura, Y., Shimohama, S., Ota, T., Matsuoka, Y., Nomura, Y., and Taniguchi, T. Alteration of transcription factors NF-*κ*B and STAT1 in Alzheimer’s disease brains. Neurosci. Lett. 237(1), 17–20 (1997).

[53] Citron, B. A., Dennis, J. S., Zeitlin, R. S., and Echeverria, V. Transcription factor Sp1 dysregulation in Alzheimer’s disease. J. Neurosci. Res. 86(11), 2499–2504 (2008).

[54] Tiwari, P. C. and Pal, R. The potential role of neuroinflammation and transcription factors in Parkinson disease. Dialogues Clin. Neurosci. 19(1), 71–80 (2017).

[55] PsychENCODE Consortium, Akbarian, S., Liu, C., Knowles, J. A., Vaccarino, F. M., Farnham, P. J., Crawford, G. E., Jaffe, A. E., Pinto, D., Dracheva, S., Geschwind, D. H., Mill, J., Nairn, A. C., Abyzov, A., Pochareddy, S., Prabhakar, S., Weissman, S., Sullivan, P. F., State, M. W., Weng, Z., Peters, M. A., White, K. P., Gerstein, M. B., Amiri, A., Armoskus, C., Ashley-Koch, A. E., Bae, T., Beckel-Mitchener, A., Berman, B. P., Coetzee, G. A., Coppola, G., Francoeur, N., Fromer, M., Gao, R., Grennan, K., Herstein, J., Kavanagh, D. H., Ivanov, N. A., Jiang, Y., Kitchen, R. R., Kozlenkov, A., Kundakovic, M., Li, M., Li, Z., Liu, S., Mangravite, L. M., Mattei, E., Markenscoff-Papadimitriou, E., Navarro, F. C. P., North, N., Omberg, L., Panchision, D., Parikshak, N., Poschmann, J., Price, A. J., Purcaro, M., Reddy, T. E., Roussos, P., Schreiner, S., Scuderi, S., Sebra, R., Shibata, M., Shieh, A. W., Skarica, M., Sun, W., Swarup, V., Thomas, A., Tsuji, J., van Bakel, H., Wang, D., Wang, Y., Wang, K., Werling, D. M., Willsey, A. J., Witt, H., Won, H., Wong, C. C. Y., Wray, G. A., Wu, E. Y., Xu, X., Yao, L., Senthil, G., Lehner, T., Sklar, P., and Sestan, N. The PsychENCODE project. Nat. Neurosci. 18(12), 1707–1712 (2015).

[56] Prater, K. E., Green, K. J., Sun, W., Smith, C. L., Chiou, K. L., Heath, L., Rose, S., Dirk Keene, C., Kwon, R. Y., Snyder-Mackler, N., Blue, E. E., Young, J. E., Shojaie, A., Logsdon, B., Garden, G. A., and Jayadev, S. Transcriptomic profiling of myeloid cells in Alzheimer’s Disease brain illustrates heterogeneity of microglia endolysosomal subtypes (2021).

[57] Gerrits, E., Giannini, L. A. A., Brouwer, N., Melhem, S., Seilhean, D., Le Ber, I., Brainbank Neuro-CEB Neuropathology Network, Kamermans, A., Kooij, G., de Vries, H. E., Boddeke, E. W. G. M., Seelaar, H., van Swieten, J. C., and Eggen, B. J. L. Neurovascular dysfunction in GRN-associated frontotemporal dementia identified by single-nucleus RNA sequencing of human cerebral cortex. Nat. Neurosci. 25(8), 1034–1048 (2022).

[58] Pieper, A., Rudolph, S., Wieser, G. L., Götze, T., Mießner, H., Yonemasu, T., Yan, K., Tzvetanova, I., Castillo, B. D., Bode, U., Bormuth, I., Wadiche, J. I., Schwab, M. H., and Goebbels, S. NeuroD2 controls inhibitory circuit formation in the molecular layer of the cerebellum. Sci. Rep. 9(1), 1448 (2019).

[59] Pletscher-Frankild, S., Pallejà, A., Tsafou, K., Binder, J. X., and Jensen, L. J. DISEASES: text mining and data integration of disease-gene associations. Methods 74, 83–89 (2015).

[60] Nguyen, A., Rauch, T. A., Pfeifer, G. P., and Hu, V. W. Global methylation profiling of lymphoblastoid cell lines reveals epigenetic contributions to autism spectrum disorders and a novel autism candidate gene, RORA, whose protein product is reduced in autistic brain. FASEB J. 24(8), 3036–3051 (2010).

[61] Sayad, A., Noroozi, R., Omrani, M. D., Taheri, M., and Ghafouri-Fard, S. Retinoic acid-related orphan receptor alpha (RORA) variants are associated with autism spectrum disorder. Metab. Brain Dis. 32(5), 1595–1601 (2017).

[62] Liu, X., Han, D., Somel, M., Jiang, X., Hu, H., Guijarro, P., Zhang, N., Mitchell, A., Halene, T., Ely, J. J., Sherwood, C. C., Hof, P. R., Qiu, Z., Pääbo, S., Akbarian, S., and Khaitovich, P. Disruption of an evolutionarily novel synaptic expression pattern in autism. PLoS Biol. 14(9), e1002558 (2016).

[63] Hirayama, T., Yuuki, K., Tarusawa, E., Saito, S., Nakayama, H., Hoshino, N., Nakama, S., Fukuishi, T., Kawanishi, Y., Umeshima, H., Tomita, K., Yoshimura, Y., Galjart, N., Hashimoto, K., Ohno, N., and Yagi, T. CTCF loss induces giant lamellar bodies in Purkinje cell dendrites (2022).

[64] Chang, J., Gilman, S. R., Chiang, A. H., Sanders, S. J., and Vitkup, D. Genotype to phenotype relationships in autism spectrum disorders. Nat. Neurosci. 18(2), 191–198 (2015).

[65] Calderon, D., Blecher-Gonen, R., Huang, X., Secchia, S., Kentro, J., Daza, R. M., Martin, B., Dulja, A., Schaub, C., Trapnell, C., Larschan, E., O’Connor-Giles, K. M., Furlong, E. E. M., and Shendure, J. The continuum of Drosophila embryonic development at single-cell resolution. Science 377(6606), eabn5800 (2022).

[66] Efthymiou, A. G. and Goate, A. M. Late onset Alzheimer’s disease genetics implicates microglial pathways in disease risk. Mol. Neurodegener. 12(1), 43 (2017).

[67] Fang, R., Preissl, S., Li, Y., Hou, X., Lucero, J., Wang, X., Motamedi, A., Shiau, A. K., Zhou, X., Xie, F., Mukamel, E. A., Zhang, K., Zhang, Y., Behrens, M. M., Ecker, J. R., and Ren, B. Comprehensive analysis of single cell ATAC-seq data with Snap-ATAC. Nat. Commun. 12(1), 1337 (2021).

[68] Leung, S. K., Jeffries, A. R., Castanho, I., Jordan, B. T., Moore, K., Davies, J. P., Dempster, E. L., Bray, N. J., O’Neill, P., Tseng, E., Ahmed, Z., Collier, D. A., Jeffery, E. D., Prabhakar, S., Schalkwyk, L., Jops, C., Gandal, M. J., Sheynkman, G. M., Hannon, E., and Mill, J. Full-length transcript sequencing of human and mouse cerebral cortex identifies widespread isoform diversity and alternative splicing. Cell Rep. 37(7), 110022 (2021).

[69] Boggs, J. M. Myelin basic protein: a multifunctional protein. Cell. Mol. Life Sci. 63(17), 1945–1961 (2006).

[70] Harauz, G., Ladizhansky, V., and Boggs, J. M. Structural polymorphism and multifunctionality of myelin basic protein. Biochemistry 48(34), 8094–8104 (2009).

[71] Harauz, G. and Boggs, J. M. Myelin management by the 18.5-kDa and 21.5-kDa classic myelin basic protein isoforms. J. Neurochem. 125(3), 334–361 (2013).

[72] Dietz, K. C., Polanco, J. J., Pol, S. U., and Sim, F. J. Targeting human oligodendrocyte progenitors for myelin repair. Exp. Neurol. 283(Pt B), 489–500 (2016).

[73] Bulik-Sullivan, B. K., Loh, P.-R., Finucane, H. K., Ripke, S., Yang, J., Schizophrenia Working Group of the Psychiatric Genomics Consortium, Patterson, N., Daly, M. J., Price, A. L., and Neale, B. M. LD Score regression distinguishes confounding from polygenicity in genome-wide association studies. Nat. Genet. 47(3), 291–295 (2015).

[74] Finucane, H. K., Bulik-Sullivan, B., Gusev, A., Trynka, G., Reshef, Y., Loh, P.-R., Anttila, V., Xu, H., Zang, C., Farh, K., Ripke, S., Day, F. R., ReproGen Consortium, Schizophrenia Working Group of the Psychiatric Genomics Consortium, RACI Consortium, Purcell, S., Stahl, E., Lindstrom, S., Perry, J. R. B., Okada, Y., Raychaudhuri, S., Daly, M. J., Patterson, N., Neale, B. M., and Price, A. L. Partitioning heritability by functional annotation using genome-wide association summary statistics. Nat. Genet. 47(11), 1228–1235 (2015).

[75] Yang, C., Hawkins, K. E., Doré, S., and Candelario-Jalil, E. Neuroinflammatory mechanisms of bloodbrain barrier damage in ischemic stroke. Am. J. Physiol. Cell Physiol. 316(2), C135–C153 (2019).

[76] Bohlen, C. J., Friedman, B. A., Dejanovic, B., and Sheng, M. Microglia in brain development, homeostasis, and neurodegeneration. Annu. Rev. Genet. 53, 263–288 (2019).

[77] Jiang, X., Lachance, M., and Rossignol, E. Involvement of cortical fast-spiking parvalbumin-positive basket cells in epilepsy. Prog. Brain Res. 226, 81–126 (2016).

[78] Kamma, E., Lasisi, W., Libner, C., Ng, H. S., and Plemel, J. R. Central nervous system macrophages in progressive multiple sclerosis: relationship to neurodegeneration and therapeutics. J. Neuroinflammation 19(1), 45 (2022).

[79] Voet, S., Prinz, M., and van Loo, G. Microglia in central nervous system inflammation and multiple sclerosis pathology. Trends Mol. Med. 25(2), 112–123 (2019).

[80] Domingues, A. V., Pereira, I. M., Vilaça-Faria, H., Salgado, A. J., Rodrigues, A. J., and Teixeira, F. G. Glial cells in Parkinson’s disease: protective or deleterious? Cell. Mol. Life Sci. 77(24), 5171–5188 (2020).

[81] Iovino, L., Tremblay, M. E., and Civiero, L. Glutamate-induced excitotoxicity in Parkinson’s disease: the role of glial cells. J. Pharmacol. Sci. 144(3), 151–164 (2020).

[82] Demontis, D., Walters, R. K., Martin, J., Mattheisen, M., Als, T. D., Agerbo, E., Baldursson, G., Belliveau, R., Bybjerg-Grauholm, J., Bækvad-Hansen, M., Cerrato, F., Chambert, K., Churchhouse, C., Dumont, A., Eriksson, N., Gandal, M., Goldstein, J. I., Grasby, K. L., Grove, J., Gudmundsson, O. O., Hansen, C. S., Hauberg, M. E., Hollegaard, M. V., Howrigan, D. P., Huang, H., Maller, J. B., Martin, A. R., Martin, N. G., Moran, J., Pallesen, J., Palmer, D. S., Pedersen, C. B., Pedersen, M. G., Poterba, T., Poulsen, J. B., Ripke, S., Robinson, E. B., Satterstrom, F. K., Stefansson, H., Stevens, C., Turley, P., Walters, G. B., Won, H., Wright, M. J., ADHD Working Group of the Psychiatric Genomics Consortium (PGC), Early Lifecourse & Genetic Epidemiology (EAGLE) Consortium, 23andMe Research Team, Andreassen, O. A., Asherson, P., Burton, C. L., Boomsma, D. I., Cormand, B., Dalsgaard, S., Franke, B., Gelernter, J., Geschwind, D., Hakonarson, H., Haavik, J., Kranzler, H. R., Kuntsi, J., Langley, K., Lesch, K.-P., Middeldorp, C., Reif, A., Rohde, L. A., Roussos, P., Schachar, R., Sklar, P., Sonuga-Barke, E. J. S., Sullivan, P. F., Thapar, A., Tung, J. Y., Waldman, I. D., Medland, S. E., Stefansson, K., Nordentoft, M., Hougaard, D. M., Werge, T., Mors, O., Mortensen, P. B., Daly, M. J., Faraone, S. V., Børglum, A. D., and Neale, B. M. Discovery of the first genome-wide significant risk loci for attention deficit/hyperactivity disorder. Nat. Genet. 51 (1), 63–75 (2019).

[83] Nagai, J., Rajbhandari, A. K., Gangwani, M. R., Hachisuka, A., Coppola, G., Masmanidis, S. C., Fanselow, M. S., and Khakh, B. S. Hyperactivity with disrupted attention by activation of an astrocyte synaptogenic cue. Cell 177(5), 1280–1292.e20 (2019).

[84] Qian, Y., Li, J., Zhao, S., Matthews, E., Adoff, M., Zhong, W., An, X., Yeo, M., Park, C., Wang, B.-S., Southwell, D., and Josh Huang, Z. Programmable RNA Sensing for Cell Monitoring and Manipulation (2022).

[85] Kessler, M. J. and Rawlins, R. G. A 75-year pictorial history of the Cayo Santiago rhesus monkey colony. Am. J. Primatol. 78(1), 6–43 (2016).

[86] Widdig, A., Muniz, L., Minkner, M., Barth, Y., Bley, S., Ruiz-Lambides, A., Junge, O., Mundry, R., and Kulik, L. Low incidence of inbreeding in a long-lived primate population isolated for 75 years. Behav. Ecol. Sociobiol. 71 (1), 18 (2017).

[87] Hernandez-Pacheco, R., Delgado, D. L., Rawlins, R. G., Kessler, M. J., Ruiz-Lambides, A. V., Maldonado, E., and Sabat, A. M. Managing the Cayo Santiago rhesus macaque population: the role of density. Am. J. Primatol. 78(1), 167–181 (2016).

[88] Testard, C., Brent, L. J. N., Andersson, J., Chiou, K. L., Valle, J. E. N.-D., DeCasien, A. R., Acevedo-Ithier, A., Stock, M. K., Antón, S. C., Gonzalez, O., Walker, C. S., Foxley, S., Compo, N. R., Bauman, S., Ruiz-Lambides, A. V., Martinez, M. I., Skene, J. H. P., Horvath, J. E., Unit, C. B. R., Higham, J. P., Miller, K. L., Snyder-Mackler, N., Montague, M. J., Platt, M. L., and Sallet, J. Social connections predict brain structure in a multidimensional free-ranging primate society. Sci. Adv. 8(15), eabl5794 (2022).

[89] Chiou, K. L., DeCasien, A. R., Rees, K. P., Testard, C., Spurrell, C. H., Gogate, A. A., Pliner, H. A., Tremblay, S., Mercer, A., Whalen, C. S., Negrón-Del Valle, J. E., Janiak, M. C., Bauman Surratt, S. E., González, O., Compo, N. R., Stock, M. K., Ruiz-Lambides, A. V., Martínez, M. I., Cayo Biobank Research Unit, Wilson, M. A., Melin, A. D., Antón, S. C., Walker, C. S., Sallet, J., Newbern, J. M., Starita, L. M., Shendure, J., Higham, J. P., Brent, L. J. N., Montague, M. J., Platt, M. L., and Snyder-Mackler, N. Multi-region transcriptomic profiling of the primate brain reveals signatures of aging and the social environment. Nat. Neurosci. (2022).

[90] Chiou, K. L., Montague, M. J., Goldman, E. A., Watowich, M. M., Sams, S. N., Song, J., Horvath, J. E., Sterner, K. N., Ruiz-Lambides, A. V., Martínez, M. I., Higham, J. P., Brent, L. J. N., Platt, M. L., and Snyder-Mackler, N. Rhesus macaques as a tractable physiological model of human ageing. Philos. Trans. R. Soc. Lond. B Biol. Sci. 375(1811), 20190612 (2020).

[91] Desimone, R. and Ungerleider, L. G. Multiple visual areas in the caudal superior temporal sulcus of the macaque. J. Comp. Neurol. 248(2), 164–189 (1986).

[92] Born, R. T. and Bradley, D. C. Structure and function of visual area MT. Annu. Rev. Neurosci. 28, 157–189 (2005).

[93] Domcke, S., Hill, A. J., Daza, R. M., Trapnell, C., Cusanovich, D. A., and Shendure, J. sci-ATAC-seq3. protocols.io (2020).

[94] Corces, M. R., Trevino, A. E., Hamilton, E. G., Greenside, P. G., Sinnott-Armstrong, N. A., Vesuna, S., Satpathy, A. T., Rubin, A. J., Montine, K. S., Wu, B., Kathiria, A., Cho, S. W., Mumbach, M. R., Carter, A. C., Kasowski, M., Orloff, L. A., Risca, V. I., Kundaje, A., Khavari, P. A., Montine, T. J., Greenleaf, W. J., and Chang, H. Y. An improved ATAC-seq protocol reduces background and enables interrogation of frozen tissues. Nat. Methods 14(10), 959–962 (2017).

[95] Krueger, F., James, F., Ewels, P., Afyounian, E., and Schuster-Boeckler, B. TrimGalore (2021).

[96] Dobin, A., Davis, C. A., Schlesinger, F., Drenkow, J., Zaleski, C., Jha, S., Batut, P., Chaisson, M., and Gingeras, T. R. STAR: ultrafast universal RNA-seq aligner. Bioinformatics 29(1), 15–21 (2013).

[97] Warren, W. C., Harris, R. A., Haukness, M., Fiddes, I. T., Murali, S. C., Fernandes, J., Dishuck, P. C., Storer, J. M., Raveendran, M., Hillier, L. W., Porubsky, D., Mao, Y., Gordon, D., Vollger, M. R., Lewis, A. P., Munson, K. M., DeVogelaere, E., Armstrong, J., Diekhans, M., Walker, J. A., Tomlinson, C., Graves-Lindsay, T. A., Kremitzki, M., Salama, S. R., Audano, P. A., Escalona, M., Maurer, N. W., Antonacci, F., Mercuri, L., Maggiolini, F. A. M., Catacchio, C. R., Underwood, J. G., O’Connor, D. H., Sanders, A. D., Korbel, J. O., Ferguson, B., Kubisch, H. M., Picker, L., Kalin, N. H., Rosene, D., Levine, J., Abbott, D. H., Gray, S. B., Sanchez, M. M., Kovacs-Balint, Z. A., Kemnitz, J. W., Thomasy, S. M., Roberts, J. A., Kinnally, E. L., Capitanio, J. P., Skene, J. H. P., Platt, M., Cole, S. A., Green, R. E., Ventura, M., Wiseman, R. W., Paten, B., Batzer, M. A., Rogers, J., and Eichler, E. E. Sequence diversity analyses of an improved rhesus macaque genome enhance its biomedical utility. Science 370(6523), eabc6617 (2020).

[98] Virshup, I., Rybakov, S., Theis, F. J., Angerer, P., and Alexander Wolf, F. anndata: Annotated data (2021).

[99] Wolock, S. L., Lopez, R., and Klein, A. M. Scrublet: computational identification of cell doublets in single-cell transcriptomic data. Cell Syst 8(4), 281–291.e9 (2019).

[100] Qiu, C., Cao, J., Martin, B. K., Li, T., Welsh, I. C., Srivatsan, S., Huang, X., Calderon, D., Noble, W. S., Disteche, C. M., Murray, S. A., Spielmann, M., Moens, C. B., Trapnell, C., and Shendure, J. Systematic reconstruction of cellular trajectories across mouse embryogenesis. Nat. Genet. 54(3), 328–341 (2022).

[101] Wolf, F. A., Angerer, P., and Theis, F. J. SCANPY: large-scale single-cell gene expression data analysis. Genome Biol. 19(1), 15 (2018).

[102] McInnes, L. and Healy, J. UMAP: Uniform Manifold Approximation and Projection for Dimension Reduction. (2018).

[103] Polański, K., Young, M. D., Miao, Z., Meyer, K. B., Teichmann, S. A., and Park, J.-E. BBKNN: fast batch alignment of single cell transcriptomes. Bioinformatics 36(3), 964–965 (2020).

[104] Dong, W., Moses, C., and Li, K. Efficient k-nearest neighbor graph construction for generic similarity measures. In Proceedings of the 20th International Conference on World Wide Web, WW ’11, 577–586 (Association for Computing Machinery, York, NY, USA, 2011).

[105] R Core Team. R: a language and environment for statistical computing. R Foundation for Statistical Computing. Vienna, Austria. http://www.R-project.org, (2013).

[106] Bushnell, B. BBMap: a fast, accurate, splice-aware aligner. Technical report, Lawrence Berkeley National Lab, (2014).

[107] Drost, H.-G. Philentropy: information theory and distance quantification with R. J. Open Source Softw. 3(26), 765 (2018).

[108] Korsunsky, I., Millard, N., Fan, J., Slowikowski, K., Zhang, F., Wei, K., Baglaenko, Y., Brenner, M., Loh, P.-R., and Raychaudhuri, S. Fast, sensitive and accurate integration of single-cell data with Harmony. Nat. Methods 16(12), 1289–1296 (2019).

[109] Bouckaert, R. R. DensiTree: making sense of sets of phylogenetic trees. Bioinformatics 26(10), 1372–1373 (2010).

[110] Schliep, K. P. phangorn: phylogenetic analysis in R. Bioinformatics 27(4), 592–593 (2011).

[111] Bolger, A. M., Lohse, M., and Usadel, B. Trimmomatic: a flexible trimmer for Illumina sequence data. Bioinformatics 30(15), 2114–2120 (2014).

[112] Langmead, B. and Salzberg, S. L. Fast gapped-read alignment with Bowtie 2. Nat. Methods 9(4), 357–359 (2012).

[113] Zhang, Y., Liu, T., Meyer, C. A., Eeckhoute, J., Johnson, D. S., Bernstein, B. E., Nusbaum, C., Myers, R. M., Brown, M., Li, W., and Liu, X. S. Modelbased analysis of ChIP-Seq (MACS). Genome Biol. 9(9), R137 (2008).

[114] Gaspar, J. M. Improved peak-calling with MACS2 (2018).

[115] Pedregosa, F., Varoquaux, G., Gramfort, A., Michel, V., Thirion, B., Grisel, O., Blondel, M., Prettenhofer, P., Weiss, R., Dubourg, V., Vanderplas, J., Passos, A., Cournapeau, D., Brucher, M., Perrot, M., and Duchesnay, É. Scikit-learn: machine learning in Python. J. Mach. Learn. Res. 12, 2825–2830 (2011).

[116] Bredikhin, D., Kats, I., and Stegle, O. MUON: multimodal omics analysis framework. Genome Biol. 23(1), 42 (2022).

[117] Satija, R., Farrell, J. A., Gennert, D., Schier, A. F., and Regev, A. Spatial reconstruction of single-cell gene expression data. Nat. Biotechnol. 33(5), 495–502 (2015).

[118] Quinlan, A. R. and Hall, I. M. BEDTools: a flexible suite of utilities for comparing genomic features. Bioinformatics 26(6), 841–842 (2010).

[119] Machlab, D., Burger, L., Soneson, C., Rijli, F. M., Schübeler, D., and Stadler, M. B. monaLisa: an R/Bioconductor package for identifying regulatory motifs. Bioinformatics 38(9), 2624–2625 (2022).

[120] Khan, A., Fornes, O., Stigliani, A., Gheorghe, M., Castro-Mondragon, J. A., van der Lee, R., Bessy, A., Chèneby, J., Kulkarni, S. R., Tan, G., Baranasic, D., Arenillas, D. J., Sandelin, A., Vandepoele, K., Lenhard, B., Ballester, B., Wasserman, W. W., Parcy, F., and Mathelier, A. JASPAR 2018: update of the open-access database of transcription factor binding profiles and its web framework. Nucleic Acids Res. 46(D1), D260–D266 (2018).

[121] Hinrichs, A. S., Karolchik, D., Baertsch, R., Barber, G. P., Bejerano, G., Clawson, H., Diekhans, M., Furey, T. S., Harte, R. A., Hsu, F., Hillman-Jackson, J., Kuhn, R. M., Pedersen, J. S., Pohl, A., Raney, B. J., Rosenbloom, K. R., Siepel, A., Smith, K. E., Sugnet, C. W., Sultan-Qurraie, A., Thomas, D. J., Trumbower, H., Weber, R. J., Weirauch, M., Zweig, A. S., Haussler, D., and Kent, W. J. The UCSC Genome Browser Database: update 2006. Nucleic Acids Res. 34(Database issue), D590–8 (2006).

