## Supplemental Figures for "A single-cell multi-omic atlas spanning the adult rhesus macaque brain"

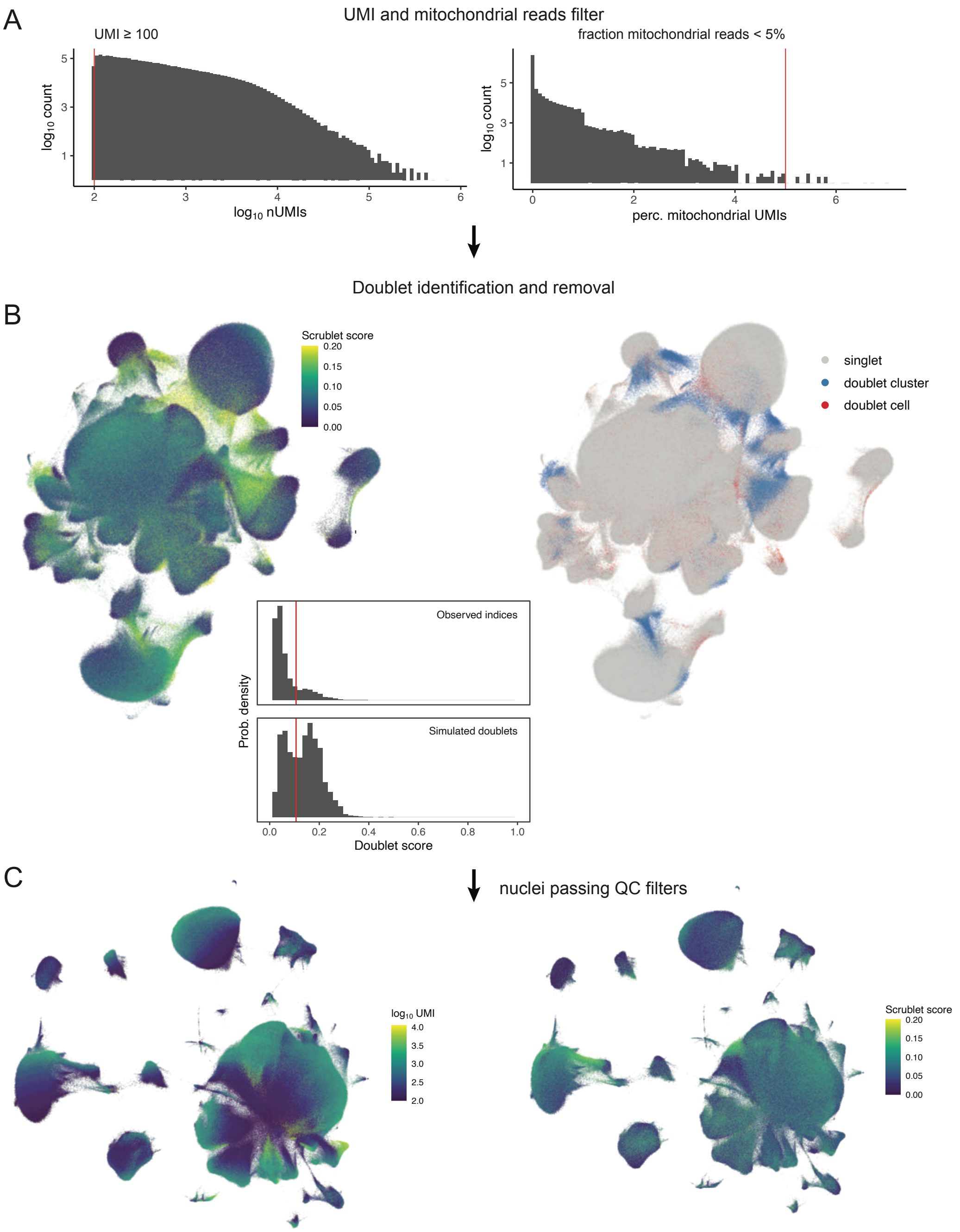

**fig. S1. Schematic depicting snRNA-seq quality control pipeline. A**, Nuclei (combinatorial indices) with fewer than 100 UMIs and greater than 5% reads mapping to the mitochondrial genome were removed. **B**, Scrublet *k*-nearest-neighbor (kNN) doublet scores were calculated per-sample and doublets with scores > 0.20 were marked (using automated Scrublet thresholds with manual adjustment) but not removed. All nuclei, including doublets, were then jointly preprocessed and clustered. Clusters with mean doublet scores > 0.15 were then removed along with previously marked doublets. **C**, UMI counts and Scrublet doublet-detection scores visualized on the post-quality-control dataset.

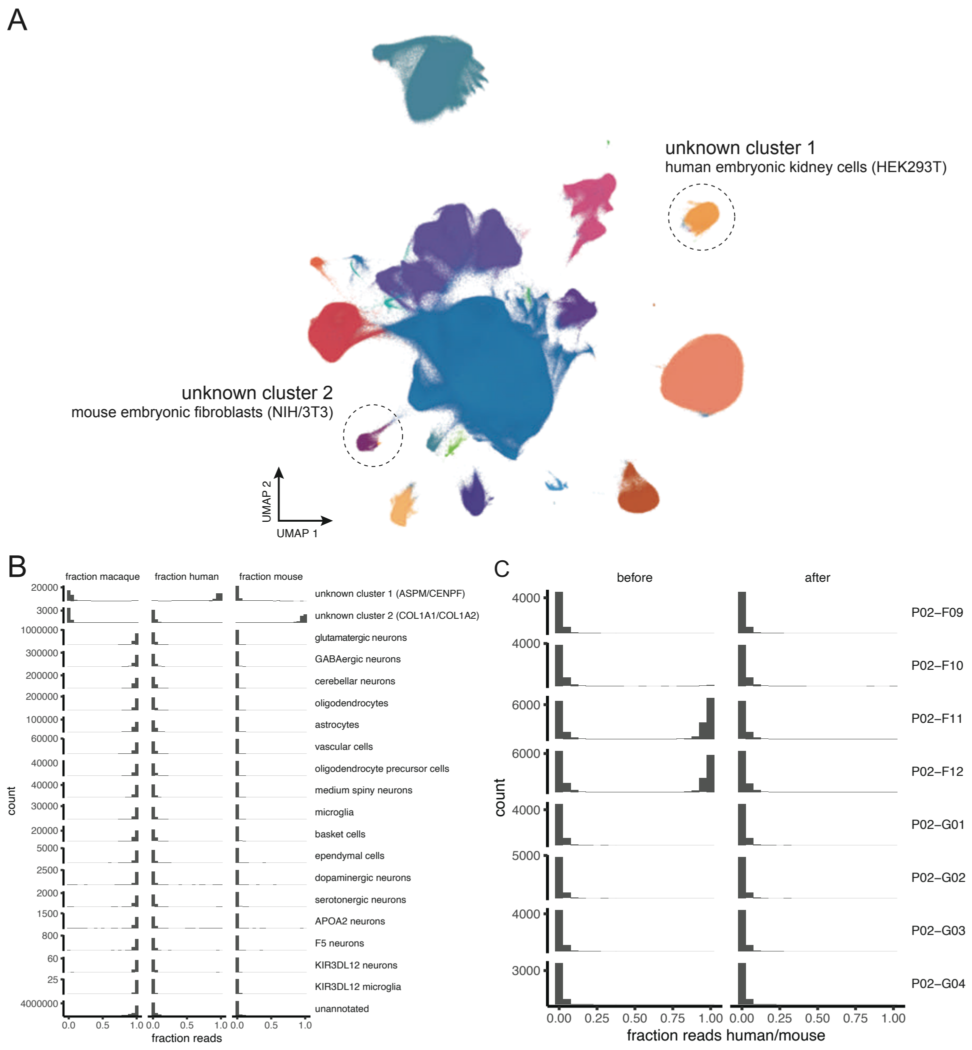

**fig. S2. Identification and removal of exogenous nuclei.** **A**, Two anomalous clusters were identified during the course of cell-type annotation had marker gene profiles (unknown cluster 1: *ASPM*, *CENPE*, *CENPF*, *MKI67*; unknown cluster 2: *COL1A1*, *COL1A2*, *FN1*, *VIM*) characteristic of stem cell progenitors. **B**, Using the BBSplit multi-genome mapping strategy, reads were assigned to either the rhesus macaque, human, or mouse genomes. Histograms showed that exogenous (human or mouse) reads were specific to the two anomalous clusters and identified them as human-derived (unknown cluster 1) and mouse-derived (unknown cluster 2), respectively. **C**, Histograms of exogenous read fractions reveal that exogenous reads were specific to particular reverse-transcription (RT) barcodes. The 8 barcodes shown (named according to plate number and position in 96-well plate) were assigned to equal aliquots of a single tissue sample, the right SPP from individual 2C0. Reads associated with two barcodes (P02-F11 and P02-F12) showed clear evidence of contamination (notably, a human-mouse barnyard control sample was loaded in adjacent wells P02-G11 and P02-G12). After the two anomalous clusters were removed from the entire dataset, these two barcodes no longer showed discernible evidence of exogenous contamination, indicating that human- and mouse-derived nuclei had been effectively partitioned and removed from the dataset. We observed similar patterns with some other samples, though with much lower degrees of contamination.

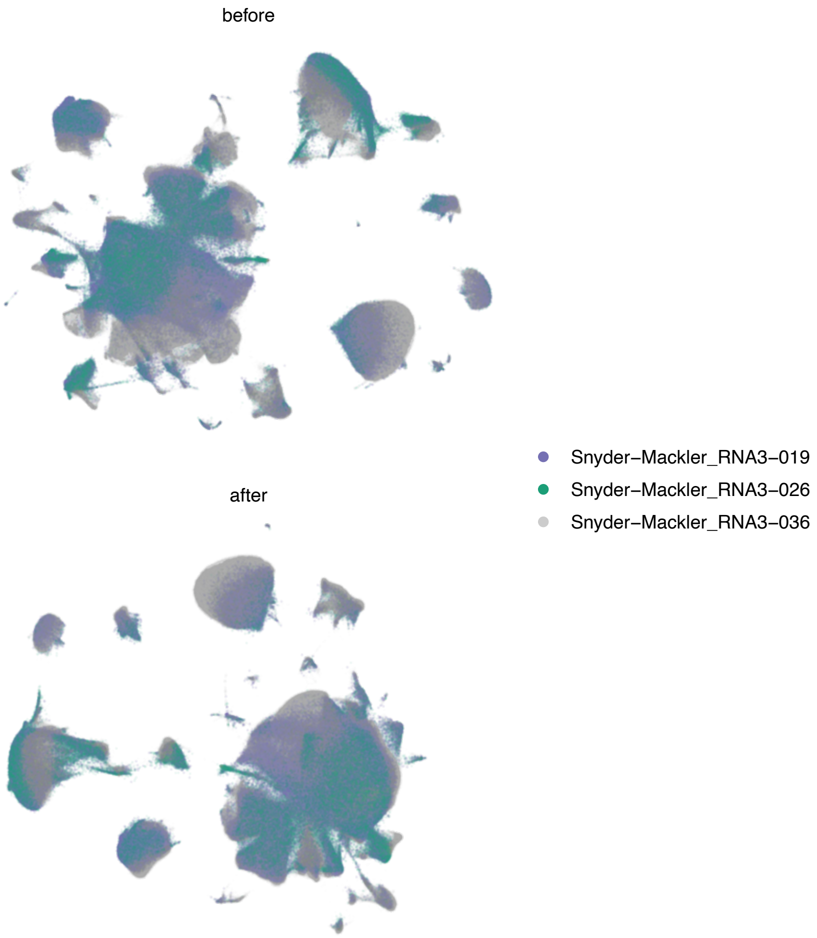

**fig. S3. Comparison of UMAP projections before batch and after batch correction.** The UMAP projection prior to batch correction was generated using the Scanpy ‘neighbors‘ function to build a neighborhood-graph while the UMAP projection with batch correction used BBKNN in place of ‘neighbors‘. Colors highlight the three library-preparation/sequencing batches, with the third batch shown in a lighter gray color with increased transparency due to the higher nuclei numbers from this batch implementing protocol improvements.

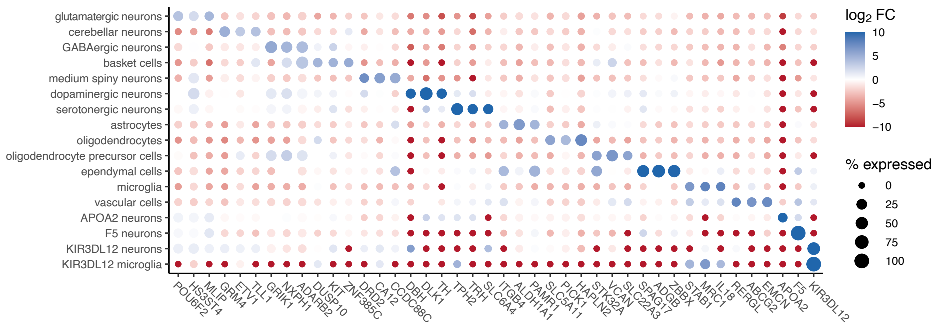

**fig. S4. Dot plot showing the top three marker genes (ranked by regularized logistic regression coefficient) for each cell class identified by our analysis.** Marker genes with percent expression < 10% were excluded, as were novel genes lacking gene names. For the four rarest cell classes, only the top marker is shown (*KIR3DL1/2* was the top marker gene for two rare cell classes). $\text{Log}_2$ fold-change (FC) color ranges are capped at $\pm 10$.

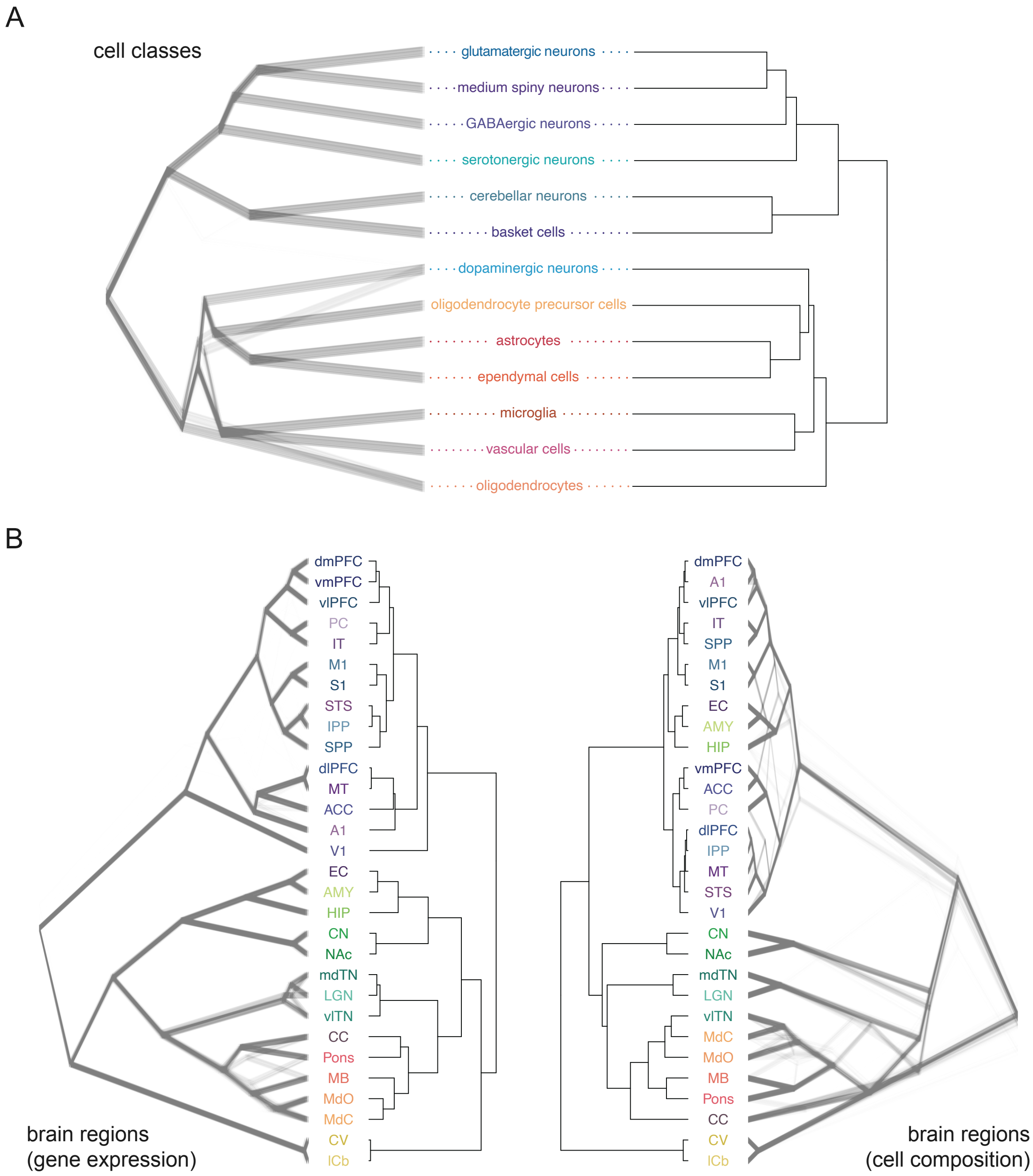

**fig. S5. Unsupervised clustering of cell classes and brain regions. A**, Dendrograms showing unsupervised hierarchical clustering of cell classes by the top 50 principal components of gene expression. The consensus tree is shown on the right, opposite an uncertainty tree derived from 1,000 bootstrap replicates. **B**, Dendrograms showing unsupervised clustering of brain regions by, left, the top 50 principal components of gene expression and, right, relative proportions of cell classes. Consensus trees are shown opposite uncertainty trees which were also each derived from 1,000 bootstrap replicates.

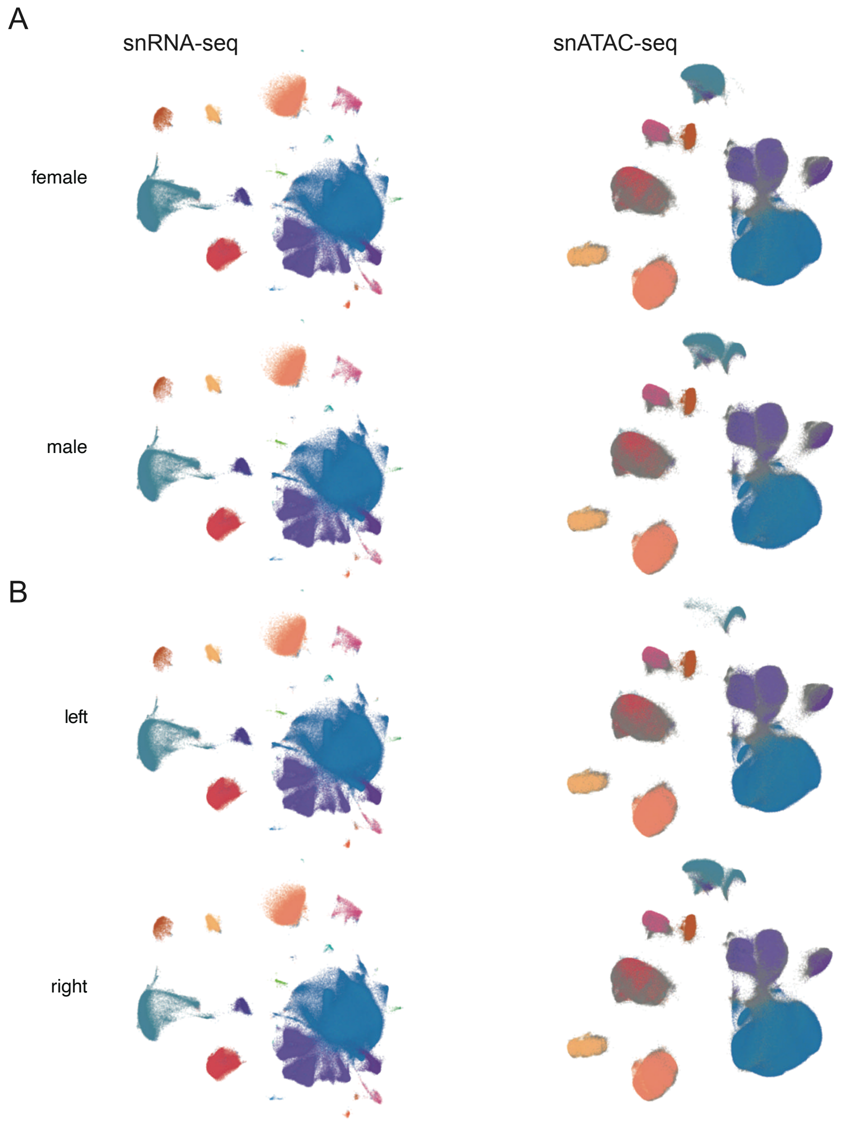

**fig. S6. Sex and hemisphere balance.** Comparison of UMAP embeddings of nuclei derived from samples of different **A**, biological sex and **B**, brain hemisphere. snRNA-seq data are shown on the left and snATAC-seq data are shown on the right. For the snATAC-seq dataset, nuclei lacking cell-class assignments are shown in gray. All other colors follow the color scale in **Fig. 1B** and **Fig. 2G**. For hemisphere comparisons, nuclei from the cerebellar vermis (Vrm) and midbrain (MB) are not shown because the structures are located on the midline and were sampled from either or both hemisphere(s) depending on where they were located.

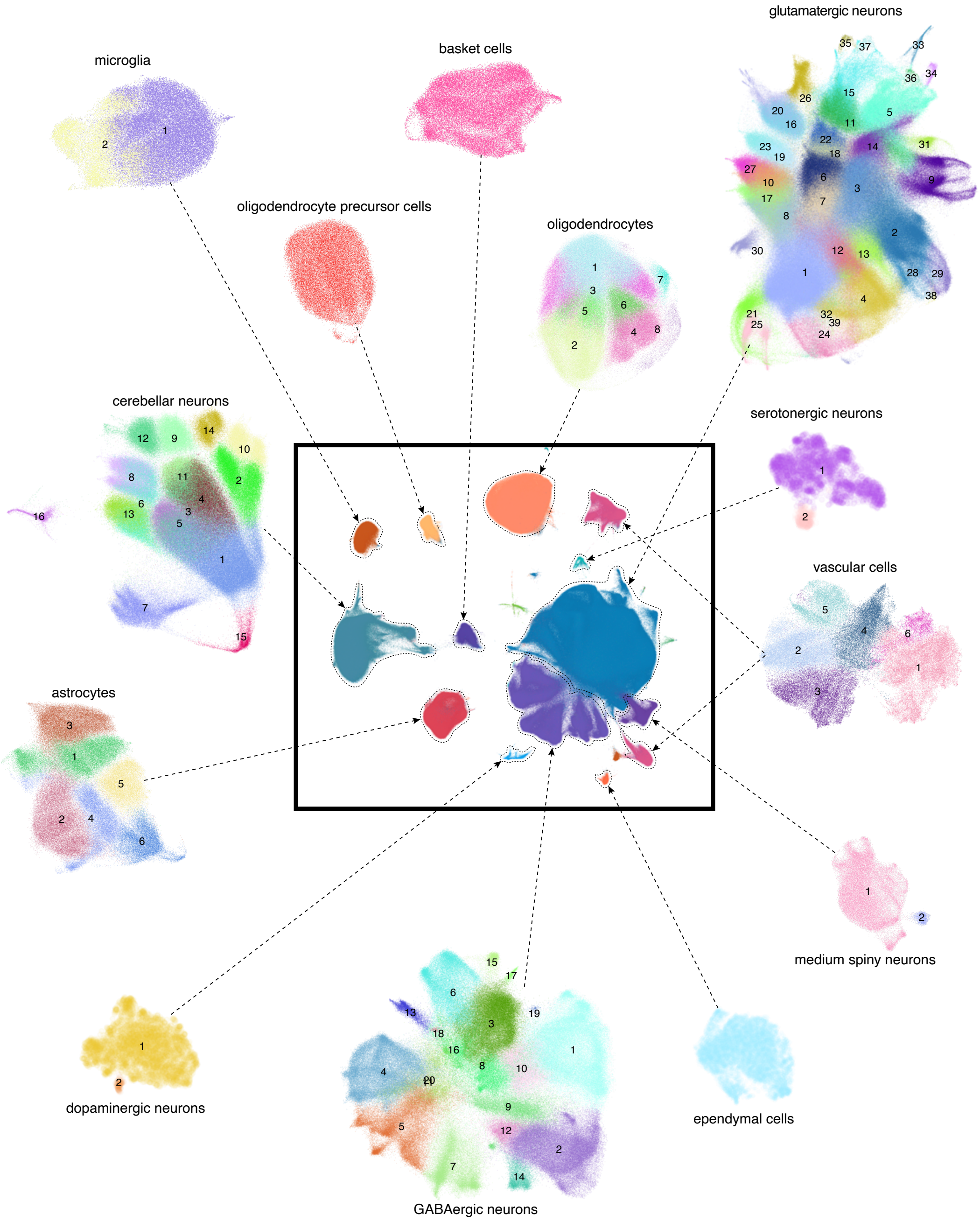

**fig. S7. Cell-class specific UMAP projections colored and labeled according to identified cell subtypes.** To identify cell subtypes, the dataset was partitioned by cell class and preprocessing, clustering, and annotation steps were repeated on each partition separately. Cell subtype colors were generated separately for each cell class partition using the randomcoloR/v.1.1.0.1 package in R.

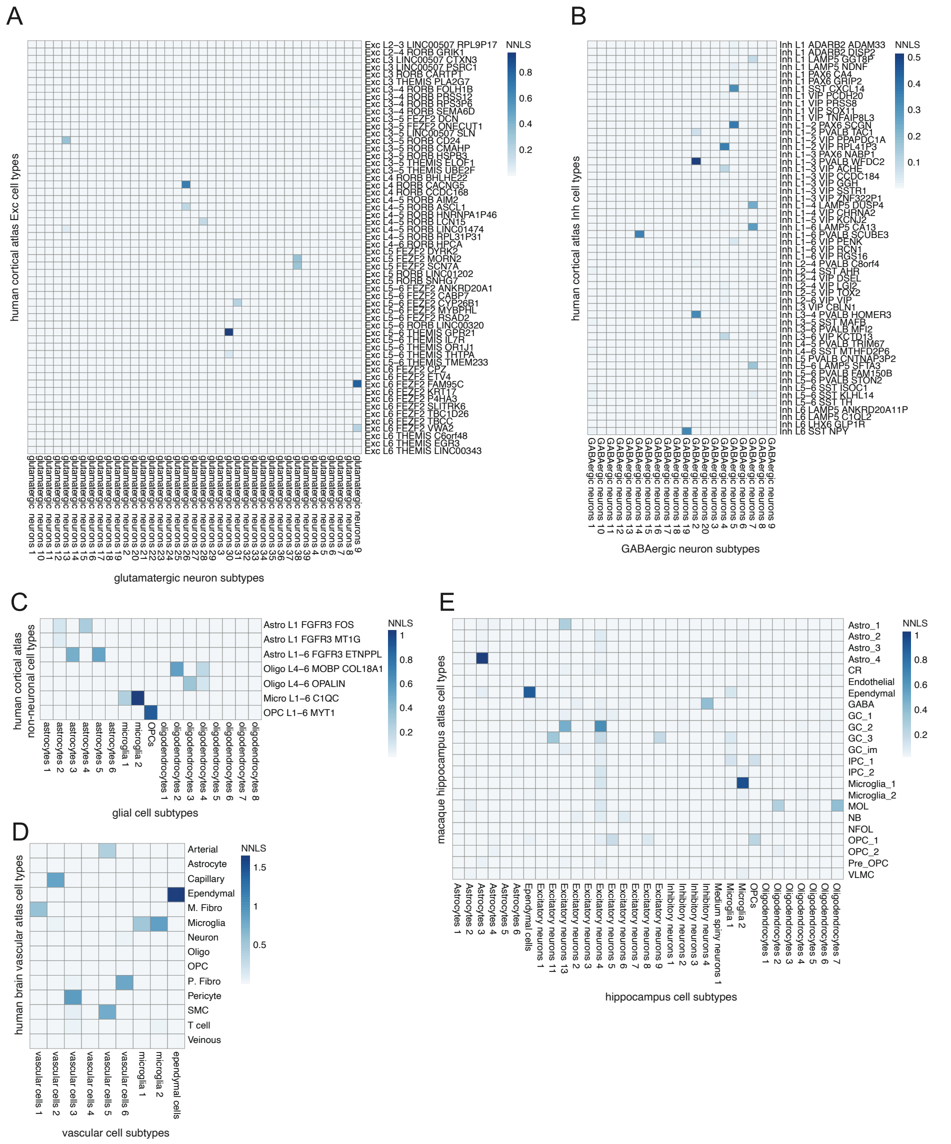

**fig. S8. Cell subtype concordance with human cortical, human brain vascular cell and macaque hippocampus atlases.** **A**-**C**, Correlations between cell subtypes and annotated labels in a cortical human brain atlas. Combined $\beta$ values from bi-directional non-negative least squares (NNLS) regression are shown for adult macaque cell subtypes (x axis) and cortical cell types annotated in the Allen human cortex dataset comprising several cortical brain regions (y axis). Glutamatergic neuron, GABAergic neuron and glial subtypes are shown in three separate panels. **D**, Correlations between cell subtypes and annotated labels in a human brain vascular cell atlas. Combined $\beta$ values from bi-directional NNLS regression are shown for adult macaque cell subtypes (vascular, myeloid, and ependymal cells, x axis) and reference cell types in the vascular cell atlas (y axis) . **E**, Correlations between cell subtypes and annotated labels in a macaque hippocampus cell atlas. Combined $\beta$ values from bi-directional NNLS regression are shown for adult macaque glutamatergic neuron, GABAergic neuron and glial cell subtypes that are sufficiently abundant in the hippocampus ($N$>100, x axis) and reference cell types in the macaque hippocampus cell atlas (y axis).

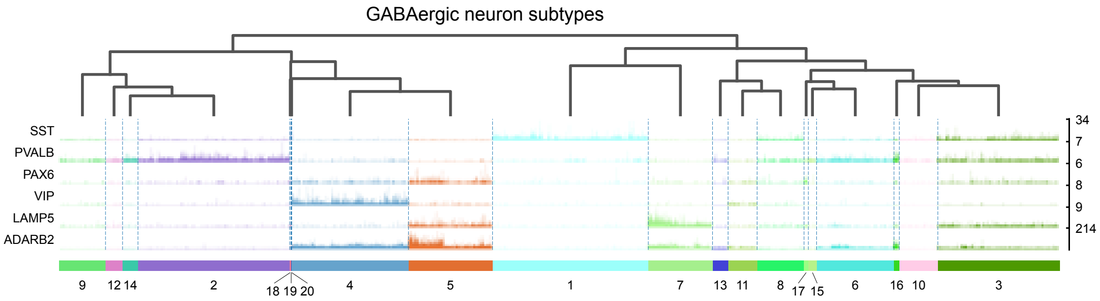

**fig. S9. “Tracks plot” showing snRNA-seq read counts for cells assigned to each of 20 GABAergic neuron subtypes for six known GABAergic neuron markers.** Dendrogram is based on hierarchical clustering of the top 50 principal components of gene expression in the GABAergic neuron class dataset partition.

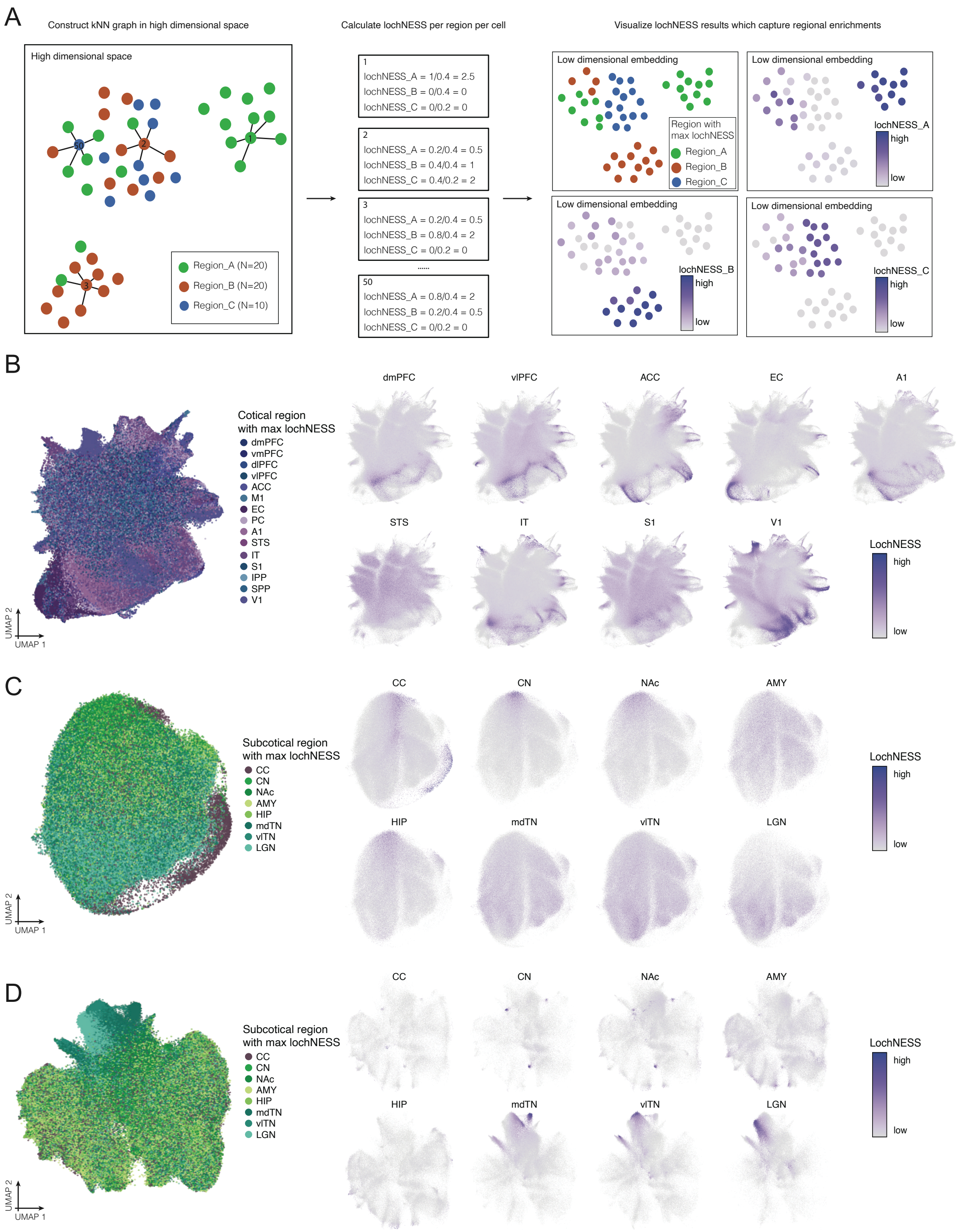

**fig. S10. Schematic of extended lochNESS and additional lochNESS examples. A**, Schematic of the lochNESS analysis. **B**, UMAP visualizations of glutamatergic neurons colored by the cortical region with the highest lochNESS (left). LochNESS distributions in a subset of cortical regions are shown in separate panels (right). **C**, UMAP visualizations of oligodendrocytes colored by the subcortical region with the highest lochNESS (left). LochNESS distribution in subcortical regions are shown in separate panels (right). **D**, UMAP visualizations of GABAergic neurons colored by the subcortical region with the highest lochNESS (left). LochNESS distribution in subcortical regions are shown in separate panels (right).

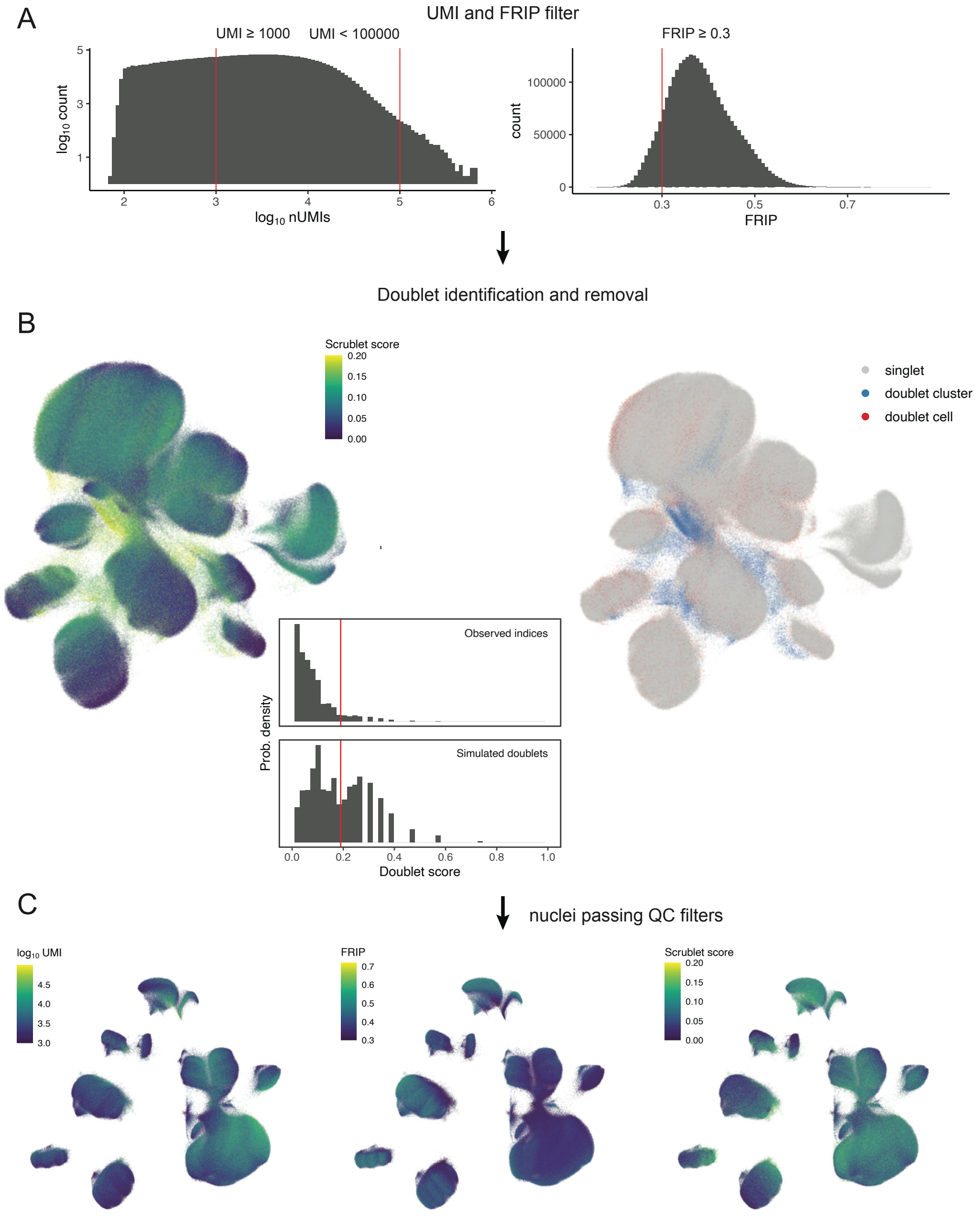

**fig. S11. Schematic depicting snATAC-seq quality control pipeline. A**, Nuclei (combinatorial indices) with fewer than 100 or greater than 100,000 UMIs were removed, as were nuclei with fractions of reads in peaks (FRIP) < 0.3. **B**, Scrublet *k*-nearest-neighbor (kNN) doublet scores were calculated per-sample and doublets with scores > 0.20 were marked (using automated Scrublet thresholds with manual adjustment) but not removed. All nuclei, including doublets, were then jointly preprocessed and clustered. Clusters with mean doublet scores > 0.15 were then removed along with previously marked doublets. **C**, UMI counts, FRIP, and Scrublet doublet-detection scores visualized on the post-quality-control dataset.

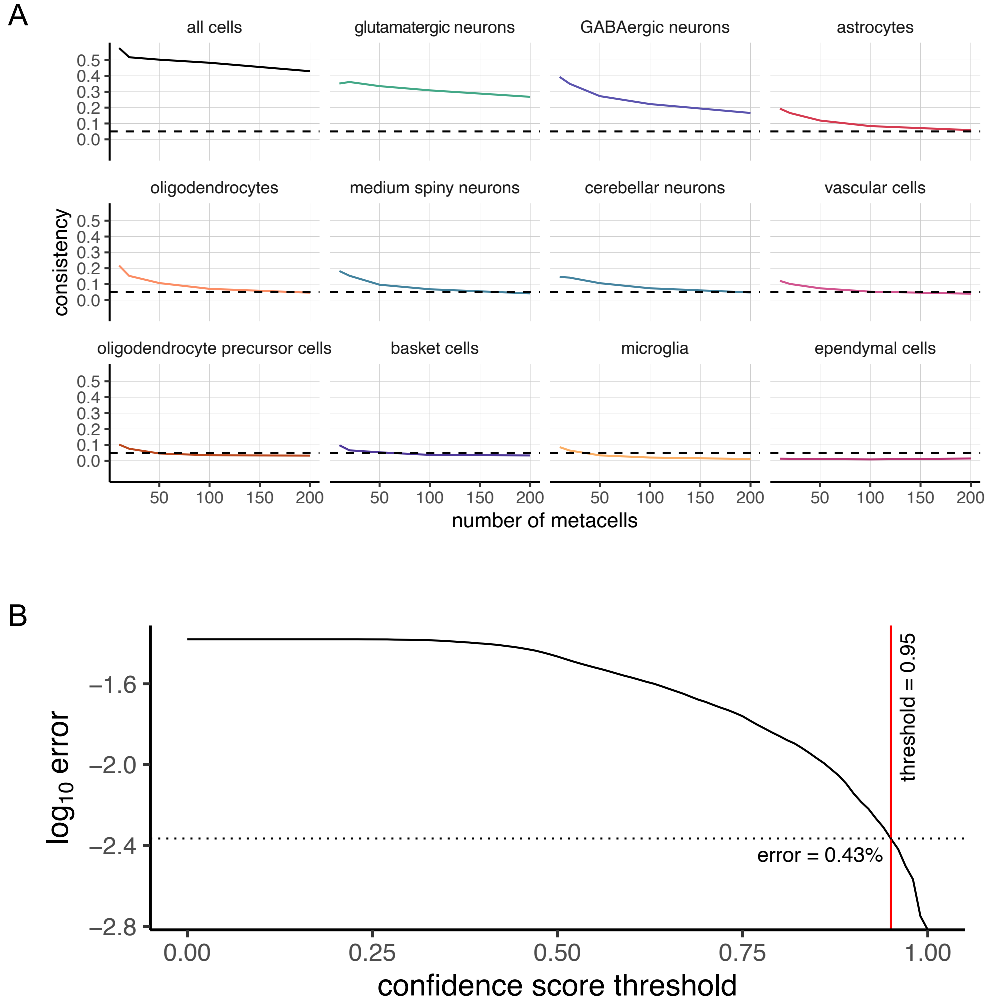

**fig. S12. Assessment of snRNA-seq/snATAC-seq integration quality. A**, Integration consistency scores—calculated by grouping neighboring cells into "metacells" and computing correlations—were calculated using glue and are plotted here. These plots include both integrations performed at the dataset-wide level ("all cells") and at the cell-class-specific level. Integrations are considered more reliable the higher the curve is. We observed that larger cell classes tended to have higher integration consistency scores. **B**, Cell-class prediction accuracy was calculated using an evaluation dataset of 100,000 snRNA-seq cells not used in the reference dataset. A range of confidence score thresholds were then tested. At a confidence threshold of 0.95 (the chosen threshold), the prediction error (percentage of incorrectly predicted cell-class labels) was 0.43%.

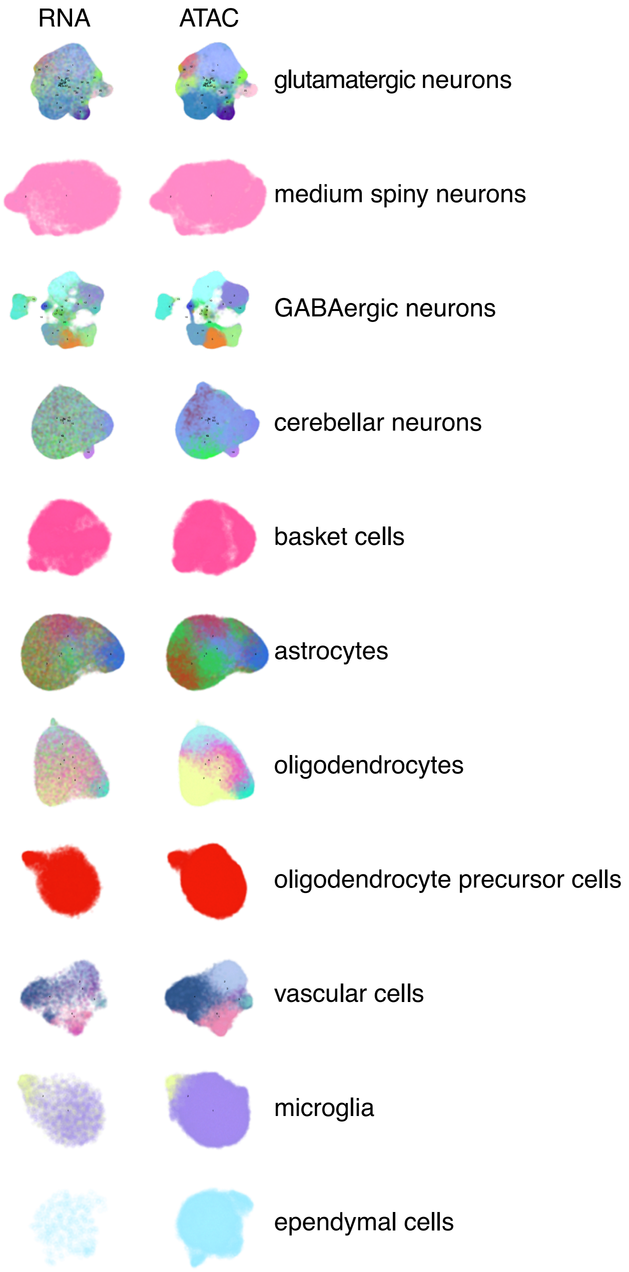

**fig. S13. UMAP embeddings of snRNA-seq and snATAC-seq data integrated separately across cell classes.** Cells are colored according to annotated or predicted cell subtypes and match the colors in **fig. S7**.

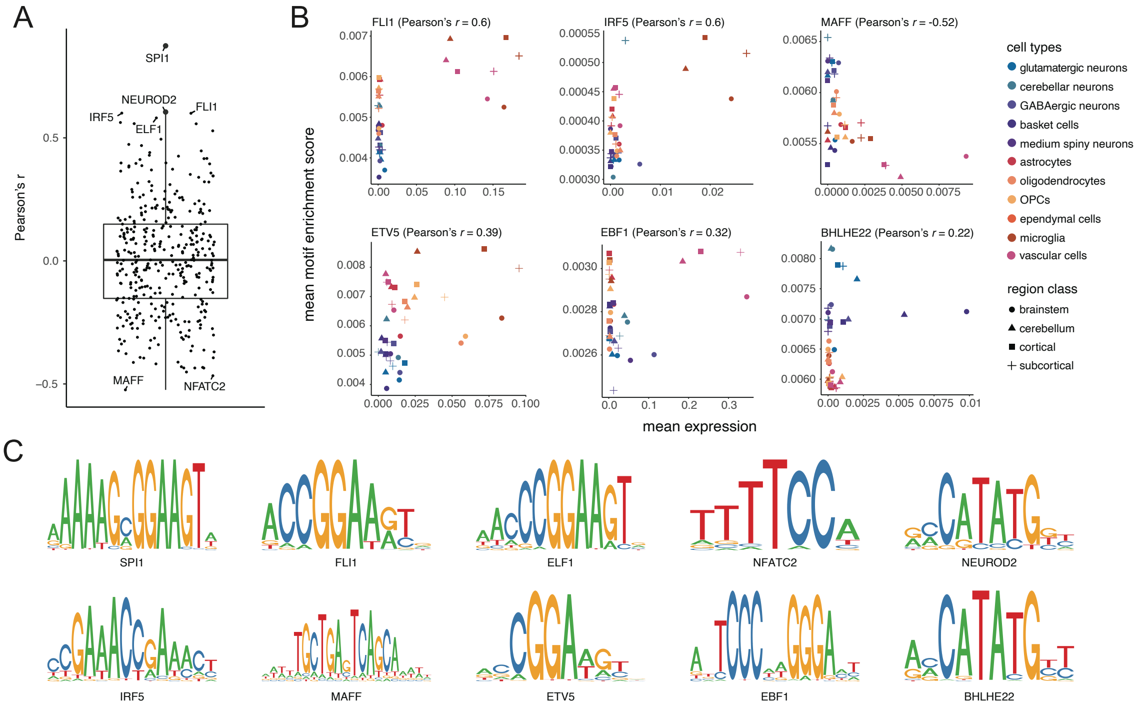

**fig. S14. Correlation of TF expression and motif enrichments.** **A**, Boxplot showing distribution of Pearson's correlation coefficients with TFs with largest coefficient values labeled. **B**, Scatterplots showing correlation between snATAC-seq accessibility of TF binding motifs and snRNA-seq gene expression of corresponding TF genes within cell classes in regional classes for six TFs (in addition to examples in **Fig. 4D**). These TFs with cell-class-specific activating and repressing effects were selected either by systematically screening for large coefficient values (top row) or manual inspection (bottom row). **C**, Position weight matrices of the ten TF motifs shown in the TF expression and motif enrichment correlation analysis.

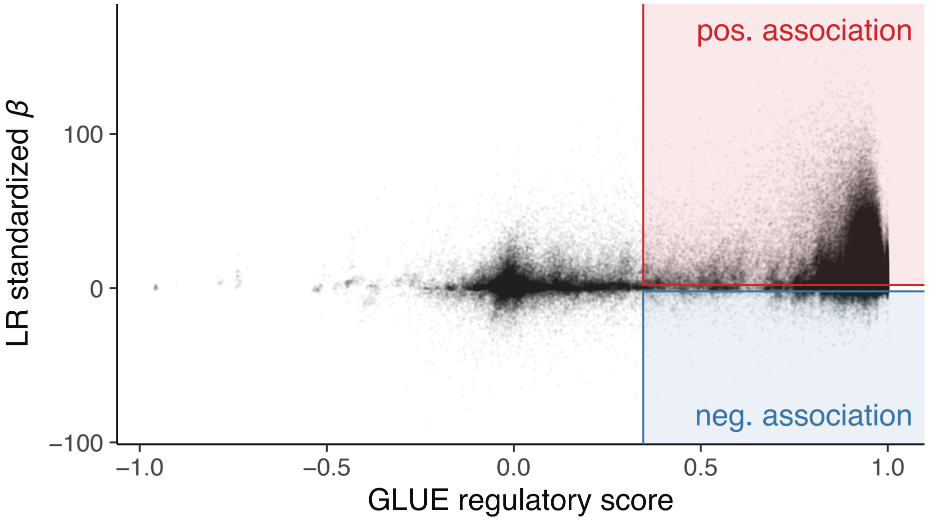

**fig. S15. Concordance between GLUE regulatory scores—which measure the cosine similarity between genes and peaks in the integrated multidimensional embedding—and logistic regression standardized effect sizes (standardized $\beta$), calculated based on meta-cells.** Shaded areas encompass peak-gene pairs that are identified as candidate regulatory interactions ($P_{adj} < 0.05$ for both statistics), with the color depicting the direction of the association based on the sign of the logistic regression $\beta$ estimate.

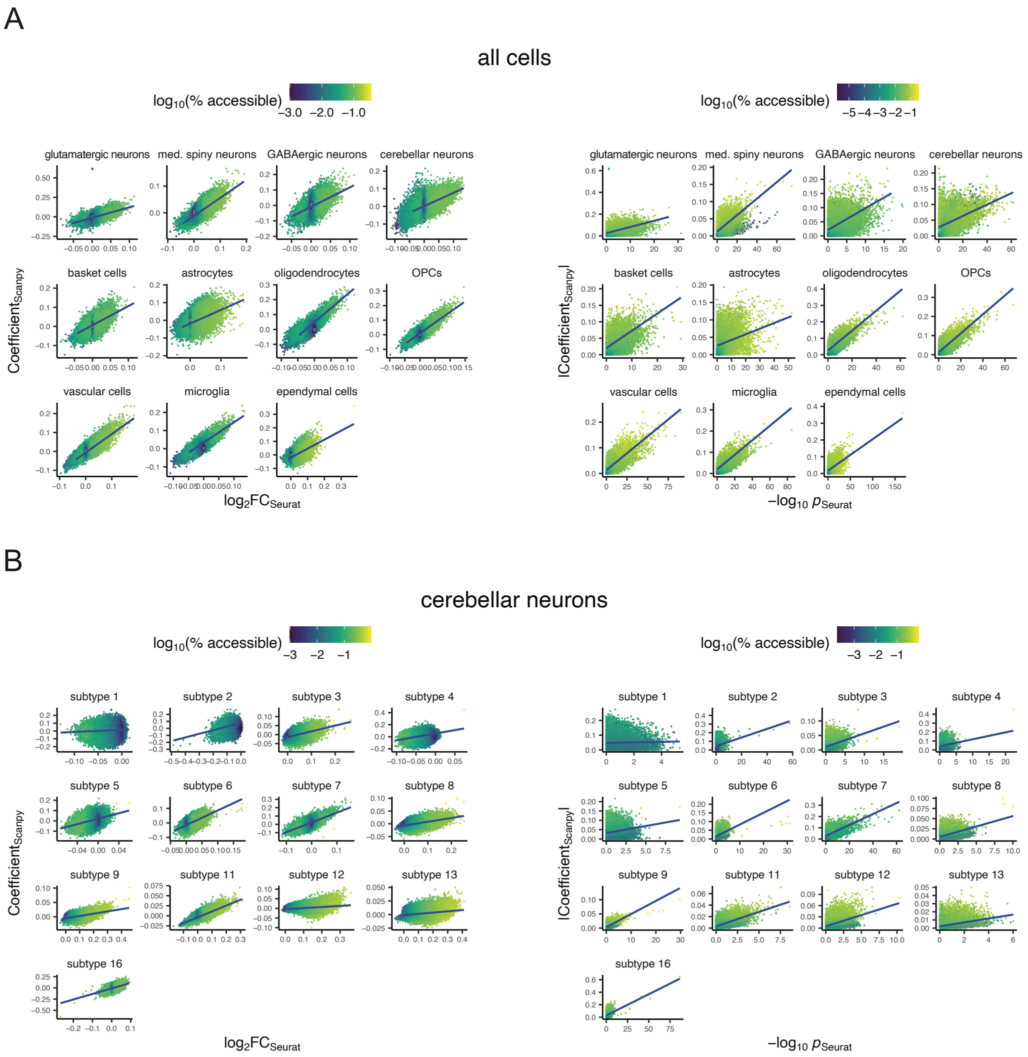

**fig. S16. Validation of marker peaks.** Due to the large size of the data, the relatively computationally tractable regularized logistic regression method implemented in Scanpy was used to calculate differential accessibility across peaks. The regularized logistic regression, however, is unable to calculate *P* values or to control for covariates such as UMI counts. We therefore subsampled our dataset (1,000 cells per cell class or subtype) and repeated the analysis using the Seurat implementation ('FindMarkers') of logistic regression, which allowed us to control for UMI counts and to calculate *P* values. We confirmed that results were similar, both at the **A**, class, and **B**, subtype levels (a representative subtype-level analysis is shown here for the cerebellar neuron data partition).

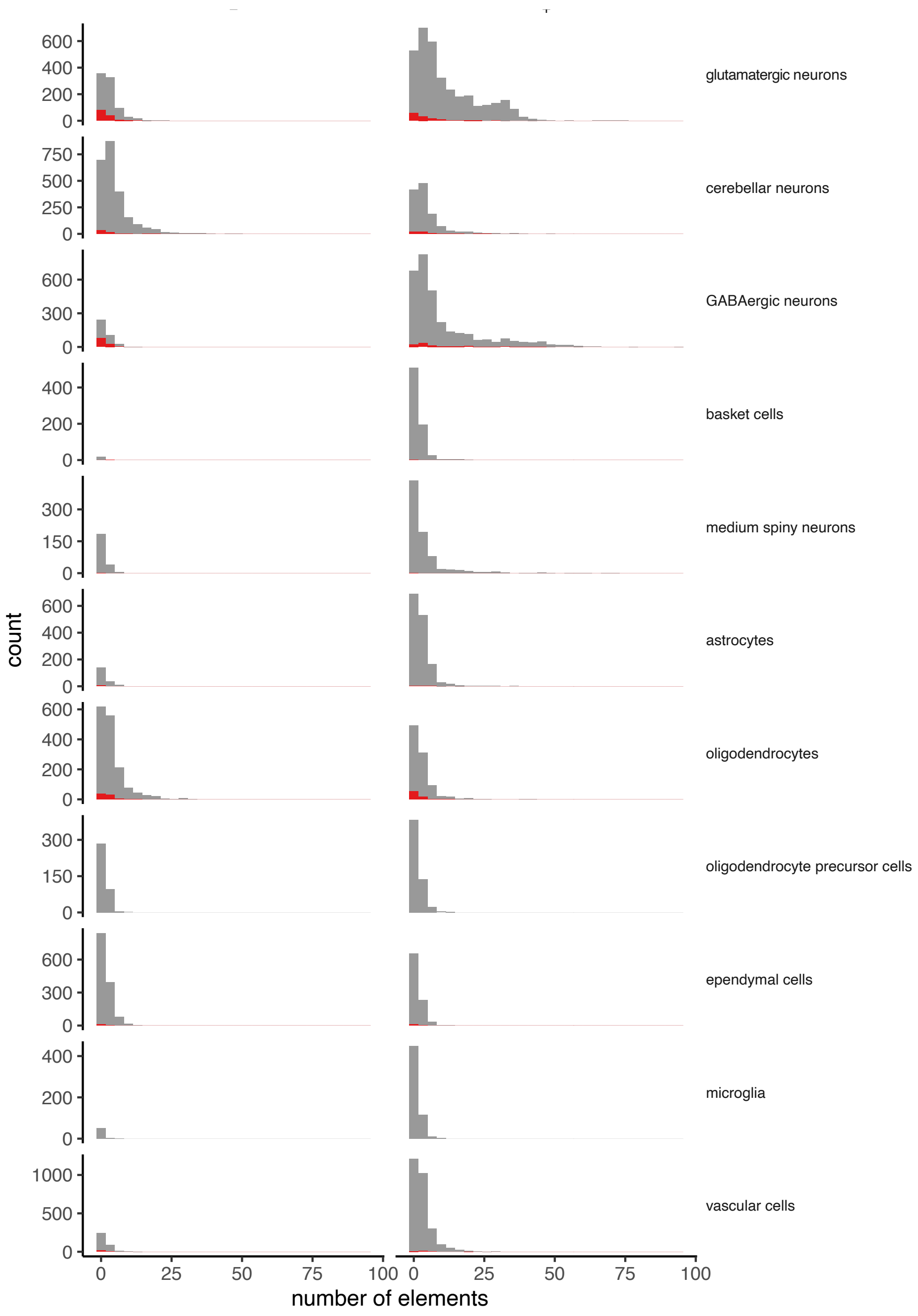

**fig. S17. Histograms showing the number of cCREs (peaks) identified as interacting with a gene for each of 11 cell classes. Interactions are grouped separately into positive and negative interactions.** Genes having both positively and negatively associated cCREs are shaded in red.
